# Synaptic connectome of a neurosecretory network in the *Drosophila* brain

**DOI:** 10.1101/2024.08.28.609616

**Authors:** Theresa H. McKim, Jayati Gera, Ariana J. Gayban, Kevin Christie, Joel Shin, Nils Reinhard, Giulia Manoli, Selina Hilpert, Patrick Callaerts, Floris van Breugel, Charlotte Helfrich-Förster, Meet Zandawala

## Abstract

Hormones mediate inter-organ signaling which is crucial in orchestrating diverse behaviors and physiological processes including sleep and activity, feeding, growth, metabolism and reproduction. The pars intercerebralis and pars lateralis in insects represent major hubs which contain neurosecretory cells (NSC) that produce various peptide hormones. To obtain insight into how hormonal signaling is regulated, we have characterized the synaptic connectome of NSC in the adult *Drosophila* brain. Identification of neurons providing inputs to multiple NSC classes implicate diuretic hormone 44-expressing NSC as a major coordinator of physiology and behavior. Surprisingly, despite most NSC having dendrites in the subesophageal zone (primary taste processing center), inputs from peripheral gustatory neurons to NSC are largely indirect. We also deciphered pathways via which diverse olfactory inputs are relayed to NSC. Linear dynamical modeling of signal propagation through the connectome identifies enteric neurons as the strongest influencers of NSC activity compared to other sensory modalities. Further, our analyses revealed substantial inputs from brain descending neurons to NSC, suggesting that descending neurons regulate both endocrine and motor output to synchronize physiological changes with appropriate behaviors. In contrast to NSC inputs, synaptic output from NSC is sparse and mostly mediated by corazonin NSC. We show that both corazonin-expressing NSC and their downstream synaptic partner DNg27 influence egg-laying. We additionally explore putative paracrine interconnectivity between NSC classes and peptide hormone pathways from NSC to peripheral tissues by analyzing single-cell transcriptomic datasets. Our comprehensive characterization of the *Drosophila* neurosecretory network connectome provides a platform to understand complex hormonal networks and how they orchestrate animal behaviors and physiology.

## Introduction

The endocrine or hormonal systems in animals play a pivotal role in regulating development and a multitude of physiological processes including growth, metabolism, and reproduction (Edgar, 2006; Nässel and Zandawala, 2019; Aref *et al*., 2024). In addition, peptide hormones can target neuronal circuits to modulate diverse behaviors ranging from feeding and locomotion to courtship and aggression (Bargmann, 2012; Marder, 2012; Taghert and Nitabach, 2012; Kim *et al*., 2017; Schoofs *et al*., 2017; Nässel and Zandawala, 2022). Peptide hormones also enable organisms to adapt to changing external environments and internal states by permitting communication between the nervous system and peripheral tissues (Friedman and Halaas, 1998; Murphy and Bloom, 2006). This inter-organ signaling is crucial in orchestrating the functions of different tissues to attain homeostasis (Droujinine and Perrimon, 2016; Castillo-Armengol *et al*., 2019; Koyama *et al*., 2025). Given its importance, it is not surprising that disrupted endocrine signaling can result in several disorders including obesity, diabetes, hypertension, infertility and growth defects amongst others (Golden *et al*., 2009; Castillo-Armengol *et al*., 2019). Understanding the regulation of endocrine signaling can thus provide insights into the prevention or treatment of endocrine-related disorders.

Although peptide hormones can be produced by several tissues, the nervous system represents a major source of hormones. In vertebrates, the hypothalamus and pituitary gland contain neurosecretory cells (NSC) that are the source of several neuropeptides/hormones (Meister, 1993). These hormones, the tissues producing them, and their target tissues are categorized into different axes which regulate distinct functions. For example, the hypothalamic-pituitary-adrenal (HPA) axis utilizes corticotropin-releasing hormone (CRH), adrenocorticotropic hormone, and cortisol to primarily regulate the stress response (Herman *et al*., 2016). The hypothalamic-pituitary-thyroid (HPT) axis controls growth and metabolism whereas the hypothalamic-pituitary-gonad (HPG) axis regulates reproductive processes, both of which utilize several different hormones (Brent, 2012; Plant, 2015). Interestingly, there is also interaction between these systems. For instance, stress-regulating CRH can act on the HPG axis to reduce the production of sex hormones and suppress gonadal function (Ferin, 1999; Maeda and Tsukamura, 2006). Hence, these axes are interconnected, which underscores the regulatory complexity of the vertebrate endocrine system.

In contrast to vertebrates, the *Drosophila* brain contains a small number of NSC in the pars intercerebralis, pars lateralis, and subesophageal zone (SEZ). These NSC project their axons towards the corpora cardiaca (CC) and corpora allata (CA), a set of endocrine glands closely associated with the aorta and anterior gut **(Figure 1A)**. Despite the large evolutionary timescale separating vertebrates and insects, there are similarities between their neuroendocrine systems (Nässel and Zandawala, 2020). These systems share significant similarities in structure, signaling pathways, and cell fate determinants during development (Hartenstein, 2006; Tessmar-Raible, 2007). Hence, the pars intercerebralis and CC are analogs of the vertebrate hypothalamus and pituitary, respectively. Strikingly, there is also conservation in some of the neuropeptides utilized by these systems. For instance, CRH is homologous to diuretic hormone 44 (DH44), and both hormones regulate stress responses (Furuya *et al*., 1995; Zandawala *et al*., 2018a; Nässel and Zandawala, 2019). Similarly, homologs of other hypothalamic neuropeptides such as prolactin-releasing peptide, neuromedin U, and gonadotropin-releasing hormone that regulate hormone release are also expressed in the *Drosophila* neuroendocrine system (Melcher *et al*., 2006; Kapan *et al*., 2012; Terhzaz *et al*., 2012; Zandawala *et al*., 2018b; Yanez-Guerra *et al*., 2020). Therefore, *Drosophila* with its smaller neuroendocrine system is an attractive model to unravel evolutionary conserved pathways which regulate peptide hormone signaling. While previous studies have characterized the connectomes of neuroendocrine centers in the larvae of *Drosophila* and *Platynereis dumerilii* (Williams *et al*., 2017; Huckesfeld *et al*., 2021), the neuroendocrine connectome of an adult animal is lacking. Given the expansion and transformation of the nervous system during metamorphosis, it remains to be seen which input pathways to NSC are conserved across animal development. Importantly, there are stark differences in physiology and behavior of larval and adult *Drosophila.* Therefore, the larval neuroendocrine system, which is mainly concerned with growth and development, is not entirely suitable to understand adult physiology and behavior.

**Figure 1:**
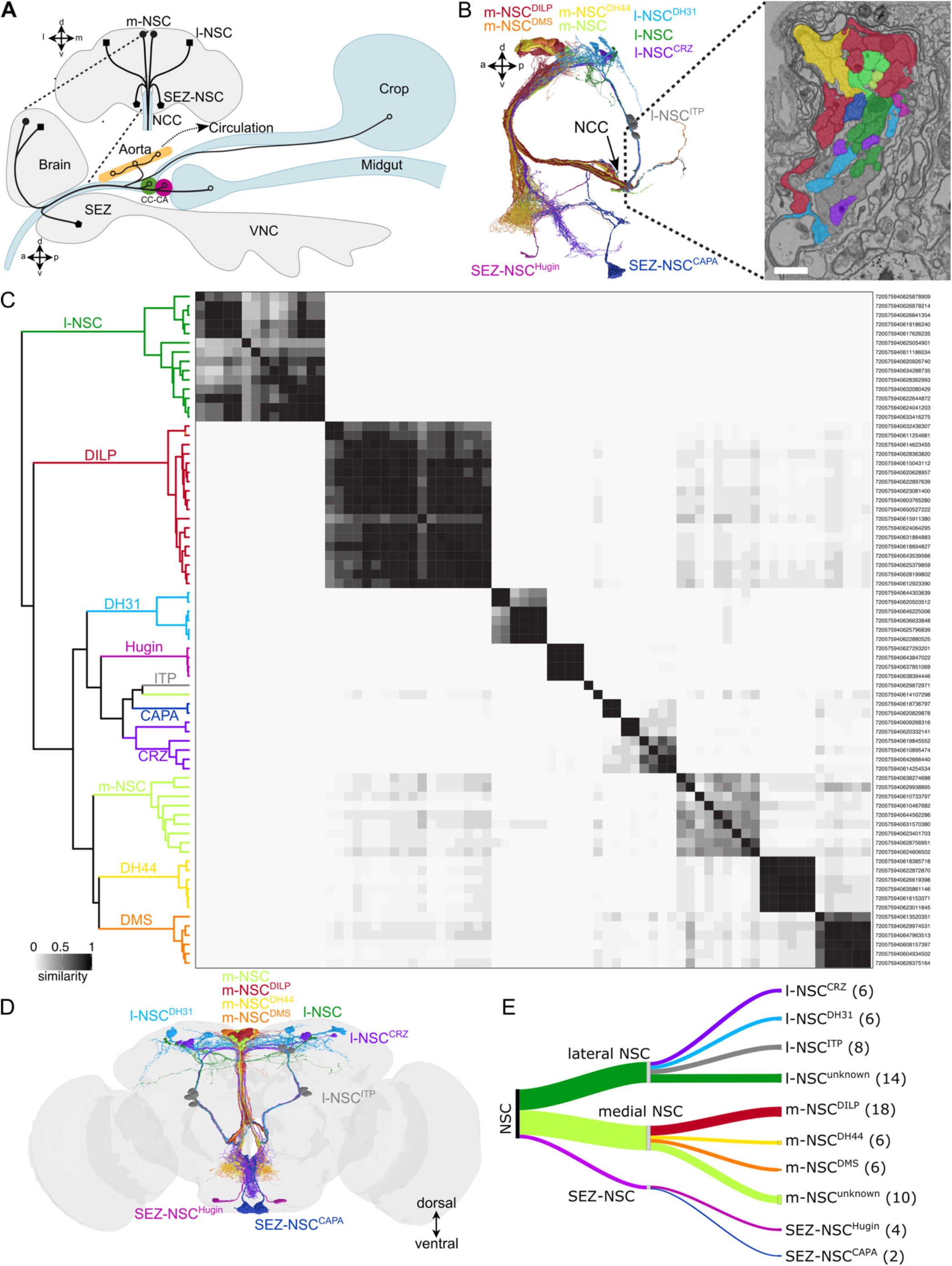
Identification of NSC in the *Drosophila* brain. **(A)** Schematic drawing of the different types of NSC and their projections to different release sites within the fly. Based on (Nässel *et al*., 2013). **(B)** All NSC projections exit the brain via the nervi corpora cardiaca (NCC). Electron micrograph showcasing a cross section of the NCC. Scale bar = 750nm. **(C)** Cosine similarity matrix of all NSC in the FlyWire connectome based on their total inputs. The darker the color, the higher the similarity between neurons. Neurons within the clades are colored based on the schematic in (D). **(D)** Reconstructions of the 80 NSC within the adult brain connectome. **(E)** Classification of NSC based on their location and neuropeptide expression. Refer to Table 1 for further details. Abbreviations: SEZ, subesophageal zone; CC, corpus cardiacum; CA, corpus allatum.

**Table 1:** Classification of *Drosophila* neurosecretory cells (NSC) based on their cell body position in the central brain.

| NSC Super Class | Neuropeptide | Other names | NSC Class | # in larvae | Expected # in adults | # in FAFB / FlyWire | # in BANC | # in maleCNS |
| --- | --- | --- | --- | --- | --- | --- | --- | --- |
| Medial | Myosuppressin (DMS) | SP3 | m-NSC<sup>DMS</sup> | 4 <sup>1</sup> | 4 <sup>2</sup> | 6 | 6 | 6 |
| Medial | Diuretic Hormone 44 (DH44) | DH44-PI | m-NSC<sup>DH44</sup> | 6 <sup>1</sup> | 6 <sup>3</sup> | 6 | 6 | 6 |
| Medial | *Drosophila* Insulin-Like Peptides (DILP) 2, 3 and 5 | IPC | m-NSC<sup>DILP</sup> | 14 <sup>1</sup> | 14 <sup>4</sup> | 18 <sup>12</sup> | 16 | 16 |
| Medial | Eclosion Hormone (EH) | V<sub>m</sub> | NSC<sup>EH</sup> | 2 <sup>1</sup> | 0 <sup>5</sup> | 0 | 0 | 0 |
| Lateral | Corazonin (CRZ) | DLP, CC-LP2 | I-NSC<sup>CRZ</sup> | 6 <sup>1</sup> | 14 <sup>6</sup> | 6 | 6 | 6 |
| Lateral | Ion-Transport Peptide (ITP) | ALK, ipc-1 | I-NSC<sup>ITP</sup> | 8 <sup>1</sup> | 8 <sup>7</sup> | 8 | 8 | 7 |
| Lateral | Diuretic Hormone 31 (DH31) | CA-LP | I-NSC<sup>DH31</sup> | 6 <sup>1</sup> | 6 <sup>8</sup> | 6 | 6 | 6 |
| Lateral | ProThoracicoTropic Hormone (PTTH) | PG-LP | I-NSC<sup>PTTH</sup> | 4 <sup>1</sup> | 0 <sup>9</sup> | 0 | 0 | 0 |
| Subesophageal | Capability (CAPA) | Large SEG neurons | SEZ-NSC<sup>CAPA</sup> | 2 <sup>1</sup> | 2 <sup>10</sup> | 2 | 2 | 2 |
| Subesophageal | Hugin | HugRG | SEZ-NSC<sup>Hugin</sup> | 4 <sup>1</sup> | 4 <sup>11</sup> | 4 | 4 | 4 |
| Medial | Unknown |  | m-NSC<sup>unknown</sup> |  |  | 10 | 9 | 8 |
| Lateral | Unknown |  | I-NSC<sup>unknown</sup> |  |  | 14 | 9 | 10 |
|  |  | **Total** |  | 56 | 58 | 80 | 72 | 71 |

**Notes:**
<sup>1</sup> (Huckesfeld *et al.*, 2021)
<sup>2</sup> This number was estimated based on expression in larvae.
<sup>3</sup> (Cabrero *et al.*, 2002; Zandawala *et al.*, 2018a)
<sup>4</sup> (Ohhara *et al.*, 2018)
<sup>5</sup> (Scott *et al.*, 2020)
<sup>6</sup> (Kapan *et al.*, 2012)
<sup>7</sup> (Kahsai *et al.*, 2010)
<sup>8</sup> (Kurogi *et al.*, 2023)
<sup>9</sup> (McBrayer *et al.*, 2007)
<sup>10</sup> (Koyama *et al.*, 2021; Zandawala *et al.*, 2021)
<sup>11</sup> (Melcher and Pankratz, 2005). This study did not quantify SEZ-NSC<sup>Hugin</sup> numbers in adults but the expression in the larvae and adults appeared similar.
<sup>12</sup> It is unclear at this point if all 18 m-NSC<sup>DILP</sup> express all three or different combinations of DILP2, 3 and 5.

To address this gap, we leveraged connectomics to characterize the first synaptic connectome of an adult neurosecretory network in an invertebrate. We deciphered all the major neuronal inputs to *Drosophila* NSC and focused on direct and indirect sensory input pathways to NSC. We also utilized single-cell transcriptomic analyses to explore putative paracrine interconnectivity between NSC classes, as well as endocrine inter-organ pathways. Our analyses shed light on the broader principles governing hormonal regulation and their impact on organismal physiology and behavior.

## Results

### Identification of 10 cell classes comprising the neuroendocrine network in the Drosophila brain

To characterize the synaptic connectome of the adult *Drosophila* neuroendocrine network, we first identified all endocrine or NSC in the brain which are a major source of circulating hormones. These endocrine cells can be broadly classified into lateral, medial, and subesophageal zone NSC (l-NSC, m-NSC, and SEZ-NSC, respectively) based on their location in the brain. Their axons exit the brain via a pair of nerves (nervii corpora cardiaca, NCC), and depending on the NSC cell type, innervate the CC, CA, hypocerebral ganglion, crop, aorta, or the anterior midgut **(Figure 1A)** (Siegmund and Korge, 2001; Hartenstein, 2006). Their axon terminals form neurohemal sites through which hormones are released into the circulation or locally on peripheral targets such as the crop. Collectively, the NSC in the brain form a major, yet distributed, neuroendocrine network that is functionally analogous to the hypothalamus (Nässel and Zandawala, 2020). We identified all brain NSC which project via the NCC in the FlyWire connectome by isolating the nerve bundle containing their axons **(Figure 1B)**. In total, we independently identified 80 brain NSC, in agreement with our companion studies (Dorkenwald *et al*., 2024; Schlegel *et al*., 2024). We propose and utilize a systematic nomenclature for all brain NSC based on their location and neuropeptide identity **(Table 1)**.

In addition to the FlyWire connectome, two additional comprehensive *Drosophila* connectomes, the brain and nerve cord (BANC) and male central nervous system (maleCNS) connectomes, are now available (Berg *et al*., 2025; Bates *et al*., 2026). Since NSC have also been identified in these datasets, we have classified them according to our scheme **(Table 1 and Figure 1 Supplement 1)** to facilitate comparison.

### NSC classes can be classified based on their morphology, neuropeptide identity, and synaptic connectivity

Interestingly, the number of NSC identified in the FlyWire connectome is larger than the number of NSC characterized in the neural connectome of the first instar larvae (Huckesfeld *et al*., 2021) **(Table 1)**. In larvae, two groups of m-NSC express myosuppressin (m-NSC^DMS^) and diuretic hormone 44 (m-NSC^DH44^). A third group of m-NSC express insulin-like peptides 2, 3, and 5 (m-NSC^DILP^), and are commonly referred to as insulin-producing cells. In addition, there are five groups of l-NSC which express ion-transport peptide (l-NSC^ITP^), corazonin (l-NSC^CRZ^), diuretic hormone 31 (l-NSC^DH31^), prothoracicotropic hormone and eclosion hormone. The latter two populations undergo apoptosis soon after adult eclosion and are thus not found in mature adults (Nässel and Zandawala, 2019; Scott *et al*., 2020). Lastly, the SEZ-NSC include two groups which express CAPA (SEZ-NSC^CAPA^) and Hugin (SEZ-NSC^Hugin^) neuropeptides. While 14 CRZ neurons are present in adults, it is not clear if all of these are bona fide NSC and release CRZ into the circulation. Since the number of neurons comprising the remaining NSC classes is thought to remain constant across development, the identity of additional NSC found in adults needed to be clarified.

With this aim in mind, we sought to classify the adult NSC into different classes based on their neuropeptide identity. All SEZ-NSC and some l-NSC classes can easily be identified based on their morphology and location **(Figure 1A)**. However, this approach is not feasible for m-NSC since they are clustered together in the superior medial protocerebrum and appear similar based on gross morphology.

Therefore, we asked whether clustering NSC based on cosine similarity between their synaptic inputs (Schlegel *et al*., 2021) can help distinguish and identify different NSC populations in the FlyWire connectome. We have recently utilized a similar approach to successfully classify neurons of the circadian clock (Reinhard *et al*., 2024). As expected, SEZ-NSC^Hugin^, SEZ-NSC^CAPA^, and l-NSC^DH31^ form three separate clusters **(Figure 1C)**. Most l-NSC^ITP^ do not have any input synapses in this dataset and were thus excluded from the analysis. However, the 8 l-NSC^ITP^ are easily recognizable based on their unique morphology **(Figure 1D-E)**. Notably, our clustering analysis resulted in two clades of m-NSC each comprising 6 neurons **(Figure 1C)**. These clusters likely represent m-NSC^DMS^ (addressed below) and m-NSC^DH44^. We also obtained two additional clusters of m-NSC comprised of 18 and 10 neurons, with the latter having low similarity between the neurons forming that cluster. We suspected that the cluster comprised of 18 m-NSC represents m-NSC^DILP^ as we expected at least 14 m-NSC^DILP^ in the connectome. To clarify the number of m-NSC^DILP^ in adults, we quantified the number of cells using various combinations of m-NSC^DILP^-specific drivers and antibodies against DILP2 or DILP3 **(Figure 2A and Figure 2 Supplement 1)**. On average, we detected 15-16 m-NSC^DILP^ with antibodies against DILPs, with several preparations labelling 18 neurons and all of them labelling greater than 10 neurons. Hence, the largest m-NSC cluster represents m-NSC^DILP^. While there are 18 m-NSC^DILP^ in the FlyWire connectome, there are 16 such cells in BANC and maleCNS connectomes, indicating inter-individual differences. Additionally, we also quantified the number of m-NSC^DMS^ since we did not retrieve any clusters with 4 neurons as was anticipated for m-NSC^DMS^. *DMS-T2A-Gal4* drives GFP expression in 6 pars intercerebralis neurons on average which project via the NCC **(Figure 2B-C)**. Surprisingly, we also detected weak GFP expression in SEZ-NSC^CAPA^ **(Figure 2B and 2D)**. Thus, there are additional m-NSC^DILP^ and m-NSC^DMS^ neurons in adults compared to larvae, and circulating/hormonal DMS can originate from two NSC classes.

**Figure 2:**
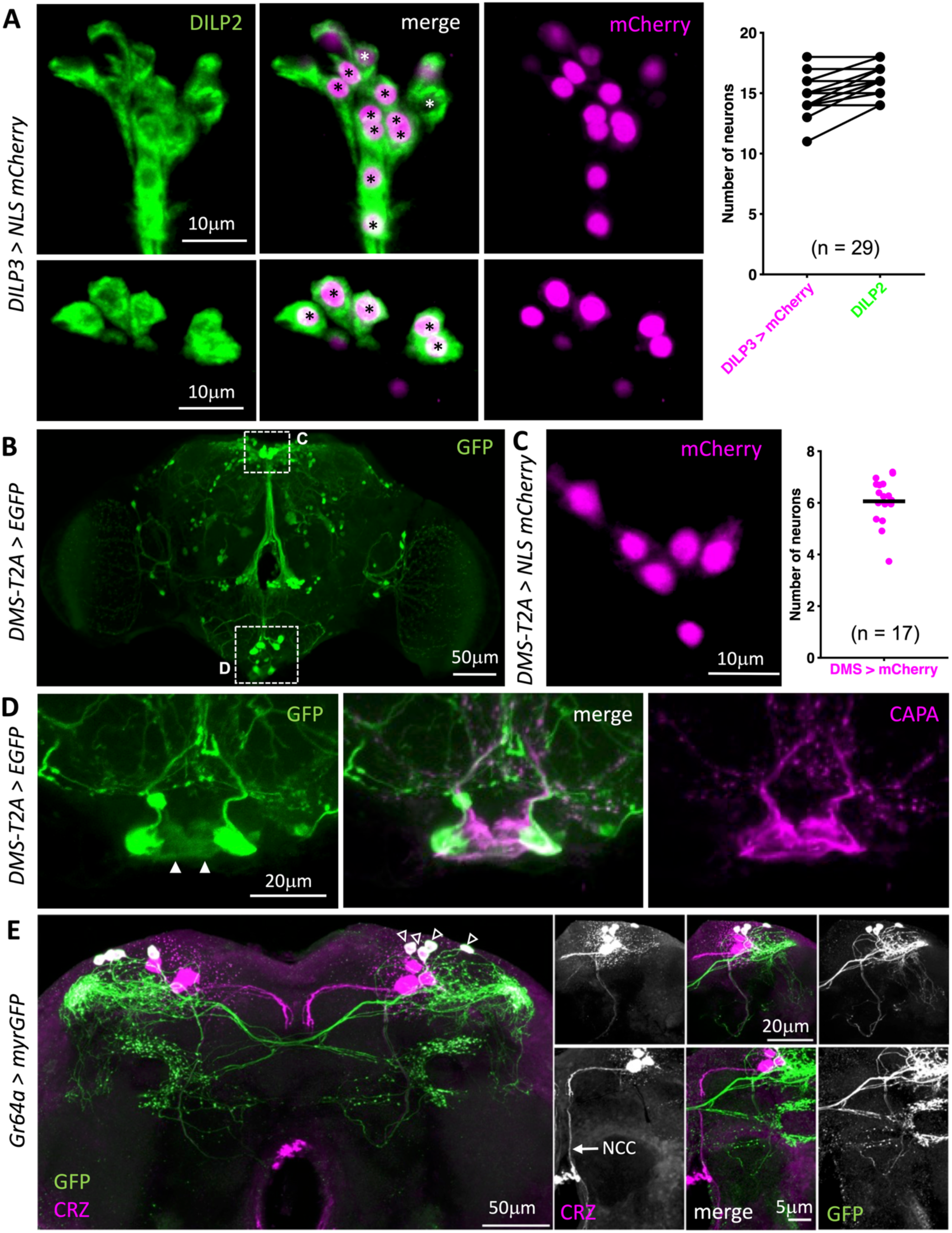
Quantification of NSC. **(A)** On average, there are 16 m-NSC^DILP^ as labelled by *DILP3-Gal4* and DILP2 antibody (marked with asterisks). Note that some preparations contain 18 m-NSC^DILP^, in agreement with the number determined based on the FlyWire connectome. **(B)** *DMS-T2A-Gal4* drives GFP expression in several neuronal populations across the brain. Regions imaged in independent samples in panels (C) and (D) are indicated using a dashed box. **(C)** *DMS-T2A-Gal4* drives nuclear mCherry expression in about six m-NSC in the pars intercerebralis. **(D)** *DMS-T2A-Gal4* drives weak GFP expression in the pair of SEZ-NSC^CAPA^ (indicated with filled arrowheads). **(E)** CRZ is expressed in 7 pairs of neurons in adult flies, 4 of which co-express *Gr64a* (empty arrowheads). These smaller *Gr64a*-expressing CRZ neurons form dense arborizations in the lateral horn. They project contralaterally but do not send projections via the nervii corpora cardiaca (NCC) and are thus not considered neurosecretory.

Intriguingly, one of the m-NSC clusters comprising 6 neurons represents m-NSC^DMS^ while the other represents m-NSC^DH44^. Since our clustering was based on input synapses, we reasoned that the differences in their pre-synaptic partners could enable us to identify these clusters. To explore this possibility, we performed retrograde trans-synaptic labelling of m-NSC^DH44^ neurons with retro-Tango (Sorkac *et al*., 2023) using a highly-specific *DH44-Gal4* for m-NSC^DH44^ **(Figure 2 Supplement 2A)**. The majority of the input to m-NSC^DH44^ originates from neurons in the SEZ which are consistently labelled across samples **(Figure 2 Supplement 3)**. Since no specific Gal4 drivers for m-NSC^DMS^ are currently available, we could not clearly map their presynaptic partners using retro-Tango. We next compared the retro-Tango output with the *in silico* retrograde tracing of 6 neurons belonging to each of the two m-NSC clusters **(Figure 2 Supplement 2B-C)**. Surprisingly, both sets of m-NSC receive a majority of their inputs from neurons in the SEZ which have similar location and morphology. However, one set of m-NSC receives inputs from a group of central neurons **(Figure 2 Supplement 2C)** that are not visible with *DH44 > retro-Tango*. We therefore refer to this cluster as m-NSC^DMS^ **(Figure 2 Supplement 2C)** and the other cluster as m-NSC^DH44^ **(Figure 2 Supplement 2B)**. To obtain additional support for our classification, we next focused on their ultrastructural features, specifically their dense core vesicles (DCVs). A recent study has shown that vesicles for different neurotransmitters can exhibit slight but significant differences that are visible in electron micrographs (Eckstein *et al*., 2024). We thus asked whether visual or morphological differences in DCVs are also observed for cells expressing DH44 and DMS. We addressed this by first identifying a pair of descending neurons (DNp32 cell type) in the connectome **(Figure 2 Supplement 2D)** which have previously been shown to express DMS (Carlsson *et al*., 2010). DCVs within these neurons are clearly visible in the soma **(Figure 2 Supplement 2G)**. These DCVs appear to be heterogenous as they are of different sizes and contrast, suggesting presence of another neuromodulator in addition to DMS. Nonetheless, the higher contrast of some DCVs in the DMS descending neuron is more similar to that of DCVs in m-NSC^DMS^ **(Figure 2 Supplement 2F)** compared to m-NSC^DH44^ **(Figure 2 Supplement 2E)**. This agrees with our hypothesis that DCVs containing the same neuromodulator appear similar visually across different cell types. To quantify these differences, we calculated the mean gray value based on at least 50 DCVs per cell for m-NSC^DMS^, m-NSC^DH44^ and DMS-expressing DNp32 in FlyWire, BANC and maleCNS connectomes **(Figure 2 Supplement 2H-J)**. For consistency, we examined the cross section of each cell where the diameter of the nucleus was largest. Our analysis shows that mean gray values of putative m-NSC^DMS^ and DMS descending neurons in FlyWire and maleCNS connectomes are not significantly different, whereas the mean gray values of m-NSC^DH44^ are significantly higher. Taken together, our synaptic connectivity and DCV analyses lend strong support to our classification of m-NSC^DH44^ and m-NSC^DMS^.

Lastly, we could only reliably identify 6 out of the expected 14 l-NSC^CRZ^ **(Figure 1C and Table 1)** (Kapan *et al*., 2012). Our inability to identify the remaining 8 CRZ neurons prompted us to examine if these adult-specific CRZ neurons are indeed neurosecretory. Previous work has shown that some of the CRZ neurons express *Gr64a* and *Gr43a* gustatory receptors (Miyamoto and Amrein, 2014; Fujii *et al*., 2015). Using *Gr64a-Gal4* to label the adult-specific CRZ neurons (Fujii *et al*., 2015), we showed that there are only 6 adult l-NSC^CRZ^ which project via the NCC **(Figure 2E).** These neurons correspond to the ones present in larvae (Huckesfeld *et al*., 2021). The 8 adult-specific CRZ neurons, labelled by *Gr64a-Gal4*, do not project via the NCC and are thus not neurosecretory. Hence, our clustering analysis accounts for all the NSC that persist into adulthood. Further, we uncovered 8-10 additional putative m-NSC and 9-14 additional putative l-NSC across the three adult brain connectomes **(Table 1**, **Figure 1C-E and Figure 1 Supplement 1)**. These neurons, especially l-NSC^unknown^, have relatively fewer DCVs than neurons such as l-NSC^ITP^ (data not shown). Hence, the type (neuropeptide, biogenic amine, or fast-acting neurotransmitter) and the identity of the signaling molecules within these neurons remain unknown. Taken together, some NSC classes have expanded in number in adults, along with an additional population of l-NSC and m-NSC **(Table 1)**.

### NSC classes comprise heterogenous cell types

Having classified NSC into 10 classes, we next compared their morphological characteristics including their cable length, surface area, cell size, and nuclei volume **(Figure 2 Supplement 4)**. SEZ-NSC^CAPA^ are about twice as large compared to other NSC types **(Figure 2 Supplement 4C)**; however, the function of pyrokinin neuropeptide produced by these cells is still unknown (Wegener *et al*., 2006). Additionally, we performed principal component analysis of the four morphological features, which revealed that the NSC of a given class generally cluster together **(Figure 2 Supplement 4E)**. However, we also observed high variability within l-NSC^CRZ^, l-NSC^DH31^, m-NSC^DH44^, and m-NSC^DILP^ populations, indicating that they are comprised of morphologically heterogenous subpopulations. Clustering based on synaptic connectivity also supports this heterogeneity as we observed multiple subclades for these NSC classes **(Figure 1C)**. For instance, the 6 l-NSC^CRZ^ cluster into two separate subclades as they represent a heterogeneous population both anatomically and functionally (Oh *et al*., 2019; Zandawala *et al*., 2021). Thus, some of the 10 NSC classes identified here are heterogenous.

### Descending neurons provide input to NSC

NSC represent a conduit through which information processed by the nervous system is relayed to peripheral tissues via different peptide hormones. As such, several neural pathways are expected to converge onto NSC. To comprehensively elucidate the inputs to NSC, we first mapped the location of input synapses for each NSC type **(Figure 3A and Figure 3 Supplement 1)**. Dendritic regions for the majority of NSC are found in the protocerebrum and SEZ. Interestingly, none of the 8 l-NSC^ITP^ had more than five synapses in the FlyWire connectome, which was the threshold used for identifying significant connections. Accordingly, their input synapses may be located outside the brain and/or the major inputs to these neurons are likely to be paracrine or hormonal in nature. We next examined the major neuronal classes providing inputs to NSC. Surprisingly, only 20 sensory neurons, all of which project to the SEZ, provide inputs despite most NSC having dendrites in that region **(Figure 3B)**. This sensory input is directed to SEZ-NSC^CAPA^, l-NSC^CRZ^, and m-NSC^DMS^ **(Figure 3C and Figure 3 Supplement 2A)**. Instead, a majority of the inputs to NSC arise from neurons in the central brain and ascending neurons from the ventral nerve cord **(Figure 3D and Figure 3 Supplement 2A)**. l-NSC^DH31^, m-NSC^DH44^, and m-NSC^unknown^ almost exclusively receive inputs from central neurons **(Figure 3 Supplement 2A)**. Interestingly, several classes of NSC receive inputs from descending neurons which are generally associated with the sensory-motor pathways that control locomotion and other behaviors (Rossignol *et al*., 2006; Namiki *et al*., 2018). This suggests that descending neurons regulate both hormonal and motor output to synchronize physiological changes with appropriate behaviors, which could explain the inhibition of m-NSC^DILP^ during locomotion (Liessem *et al*., 2023). Lastly, l-NSC^DH31^ (not shown) receive small but significant direct input from ITP-expressing visual projection neurons that are part of the circadian clock network (Kurogi *et al*., 2023; Reinhard *et al*., 2024). Overall, m-NSC^DH44^ and m-NSC^DILP^ receive the largest number of inputs, and l-NSC^ITP^ and SEZ-NSC^Hugin^ receive the least synaptic inputs **(Figure 3 Supplement 2B)**. The types of inputs to NSC, as well as their relative proportions, are consistent across the three datasets **(Figure 3 Supplement 3)**, suggesting that the broad trends observed here are not specific to an individual or sex.

**Figure 3:**
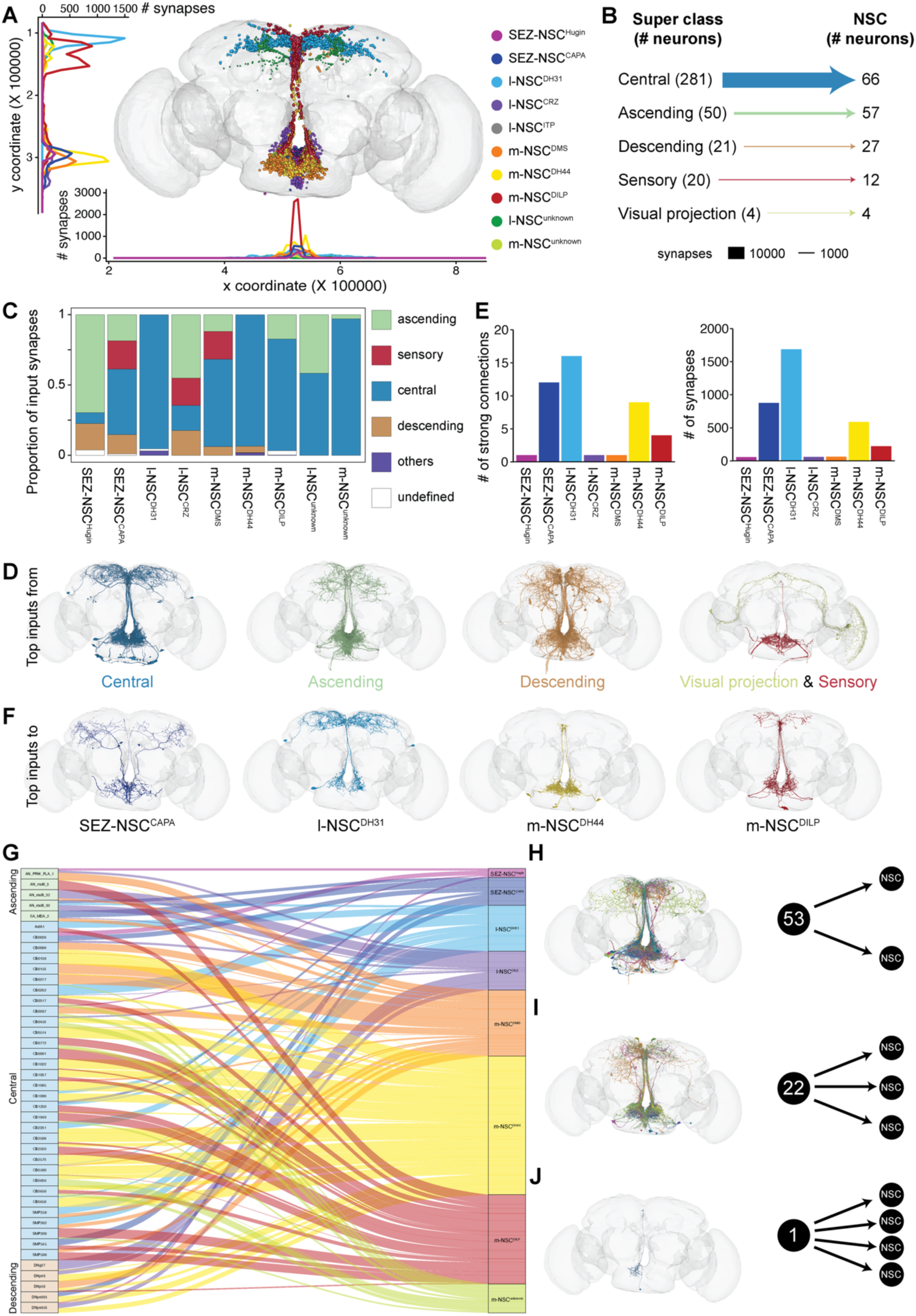
Synaptic inputs to NSC in the FlyWire connectome. **(A)** Postsynaptic sites of different NSC classes. The majority of the dendrites are found in the protocerebrum and SEZ. **(B)** Input to NSC grouped by the neuronal super classes annotated in the FlyWire connectome. Central neurons are the largest group providing inputs to NSC. **(C)** Proportion of inputs from various neuronal super classes to different NSC classes. **(D)** Reconstructions of neurons from different super classes which provide major inputs to NSC. Only the top 10 cell types per super class are shown. **(E)** Number of strong input connections (greater than 50 synapses) to each NSC class and the total number of synapses constituting these connections. **(F)** Reconstructions of neurons that provide major inputs (more than 50 synapses per connection) to SEZ-NSC^CAPA^, l-NSC^DH31^, m-NSC^DH44^ and m-NSC^DILP^. **(G)** 42 unique cell types (76 neurons) provide inputs to multiple NSC classes. The line width is scaled according to the proportion of synaptic input each of these cell types provides to different NSC classes. m-NSC^DH44^ and m-NSC^DILP^ receive the majority of these inputs. Out of these 76 neurons, **(H)** 53 neurons provide inputs to two NSC classes, **(I)** 22 neurons provide inputs to three classes of NSC and **(J)** 1 neuron provides input to four NSC classes. Reconstructions of corresponding neurons next to each schematic. For panel (E), bars have been color coded according to the legend in the panel (A).

Next, we examined the neurotransmitters expressed in the neurons presynaptic to NSC **(Figure 3 Supplement 4)**. For this, we used electron microscopy-based neurotransmitter predictions determined previously (Eckstein *et al*., 2024). We only focused on fast-acting neurotransmitters (i.e. acetylcholine, glutamate, and GABA) since those predictions were generally more reliable compared to other neurotransmitters such as serotonin (Eckstein *et al*., 2024). Both l-NSC^DH31^ and m-NSC^DH44^ receive strong glutamatergic inputs **(Figure 3 Supplement 4A)**. Out of the three fast-acting neurotransmitters, GABA provides the least inputs **(Figure 3 Supplement 4B)** consistent with the proportional usage of the three neurotransmitters across the brain (Eckstein *et al*., 2024).

Since NSC receive inputs from several different cell types whose functions are yet unknown, we focused on cells that provide strong inputs to NSC. First, we examined the number of strong input connections (≥ 50 synapses) to each NSC class **(Figure 3E)**. l-NSC^DH31^ receive the largest number of inputs via 16 strong connections, and these are comprised of over 1500 synapses in total **(Figure 3E)**. These inputs originate from neurons in the SEZ and the lateral horn **(Figure 3F)**. SEZ-NSC^CAPA^, m-NSC^DH44^, and m-NSC^DILP^ also receive substantial inputs via strong connections **(Figure 3E-F)**. In the case of SEZ-NSC^CAPA^, the strong presynaptic connections include sensory neurons and GABAergic olfactory projection neurons **(Figure 3 Supplement 2A and Figure 3 Supplement 4A)**. For both m-NSC^DH44^ and m-NSC^DILP^, the strong inputs are exclusively from the SEZ. In summary, these results suggest that sensory pathways could have a major effect on NSC activity and will be explored in detail later.

### Synaptic architecture supports orchestration of physiology by multiple hormones

Several studies have previously characterized neuroendocrine pathways which regulate different aspects of *Drosophila* physiology including metabolism, reproduction, and osmotic homeostasis (Lee *et al*., 2015; Kubrak *et al*., 2016; Oh *et al*., 2019; Hadjieconomou *et al*., 2020; Koyama *et al*., 2021; Zandawala *et al*., 2021; Kurogi *et al*., 2023; Lee *et al*., 2023; Gera *et al*., 2025). Based on these and other studies, it is becoming increasingly evident that multiple hormonal systems interact to orchestrate specific physiological processes rather than individual hormones operating in isolation. For instance, DH31, DH44, CAPA, tachykinin, and ITP, which can all be released from brain NSC, influence osmotic homeostasis via direct actions on kidney-like Malpighian tubules (Kahsai *et al*., 2010; Halberg *et al*., 2015; Zandawala *et al*., 2018a; Agard *et al*., 2024; Gera *et al*., 2025). While these hormones could be released individually under specific contexts, we anticipate some of them to be co-released to elicit an additive or synergistic response (Paluzzi *et al*., 2012; Zandawala *et al*., 2018a). As such, NSC producing these hormones could be regulated by common pre-synaptic partners. In total, we identified 76 neurons, belonging to 42 unique cell types, which provide inputs to more than one type of NSC, with m-NSC^DH44^ receiving inputs from most of these neurons **(Figure 3G and Figure 3 Supplement 5)**. This is in line with the role of DH44 in multiple processes including feeding, reproduction, metabolism, and osmotic homeostasis (Dus *et al*., 2015; Lee *et al*., 2015; Lee *et al*., 2023). Out of the 76 neurons, 53 neurons provide inputs to two types of NSC **(Figure 3H)** and 22 neurons provide inputs to three types of NSC **(Figure 3I)**. In addition, one neuron (CB3500 cell type) influences four types of NSC **(Figure 3J).** Three other neurons of the CB3500 cell type are pre-synaptic to 2-3 NSC types. Therefore, CB3500 and other neurons that provide inputs to multiple NSC could potentially integrate information from various pathways to orchestrate the release of hormones in different combinations. Taken together, this analysis provides the basis to investigate the neural control of hormonal networks regulating various physiological processes.

### NSC receive direct and indirect synaptic inputs from various sensory modalities

Sensory to endocrine pathways enable animals to maintain homeostasis by adjusting physiological processes in response to changing external environments. Since only 20 sensory neurons lie directly upstream of NSC **(Figure 3B)**, we delineated both monosynaptic and disynaptic sensory-endocrine pathways in further detail. For simplicity, we refer to interneurons mediating these connections as sensory interneurons. By extension, neurons that provide inputs to NSC but don’t receive direct sensory input are called non-sensory interneurons **(Figure 4A)**. Focusing first on the direct sensory input to NSC, only 2 gustatory receptor neurons (GRN) and 2 mechanosensory neurons are presynaptic to m-NSC^DH44^, SEZ-NSC^CAPA^, and l-NSC^CRZ^ respectively **(Figure 4A-B)**. The remaining 16 sensory neurons that provide input to m-NSC^DMS^ are enteric neurons **(Figure 4A-B)** (Giakoumas *et al*., 2025). These enteric neurons can be further classified into at least five subtypes based on morphology and synaptic connectivity **(Figure 4 Supplement 1)**. Enteric neurons project via the pharyngeal nerve to the SEZ and some of these neurons have recently been implicated in regulating sucrose, sodium or general food intake (Cui *et al*., 2024; Gao *et al*., 2024; Kim *et al*., 2024a). Thus, nutrient and mineral content of an ingested meal could influence the activity of m-NSC^DMS^.

**Figure 4:**
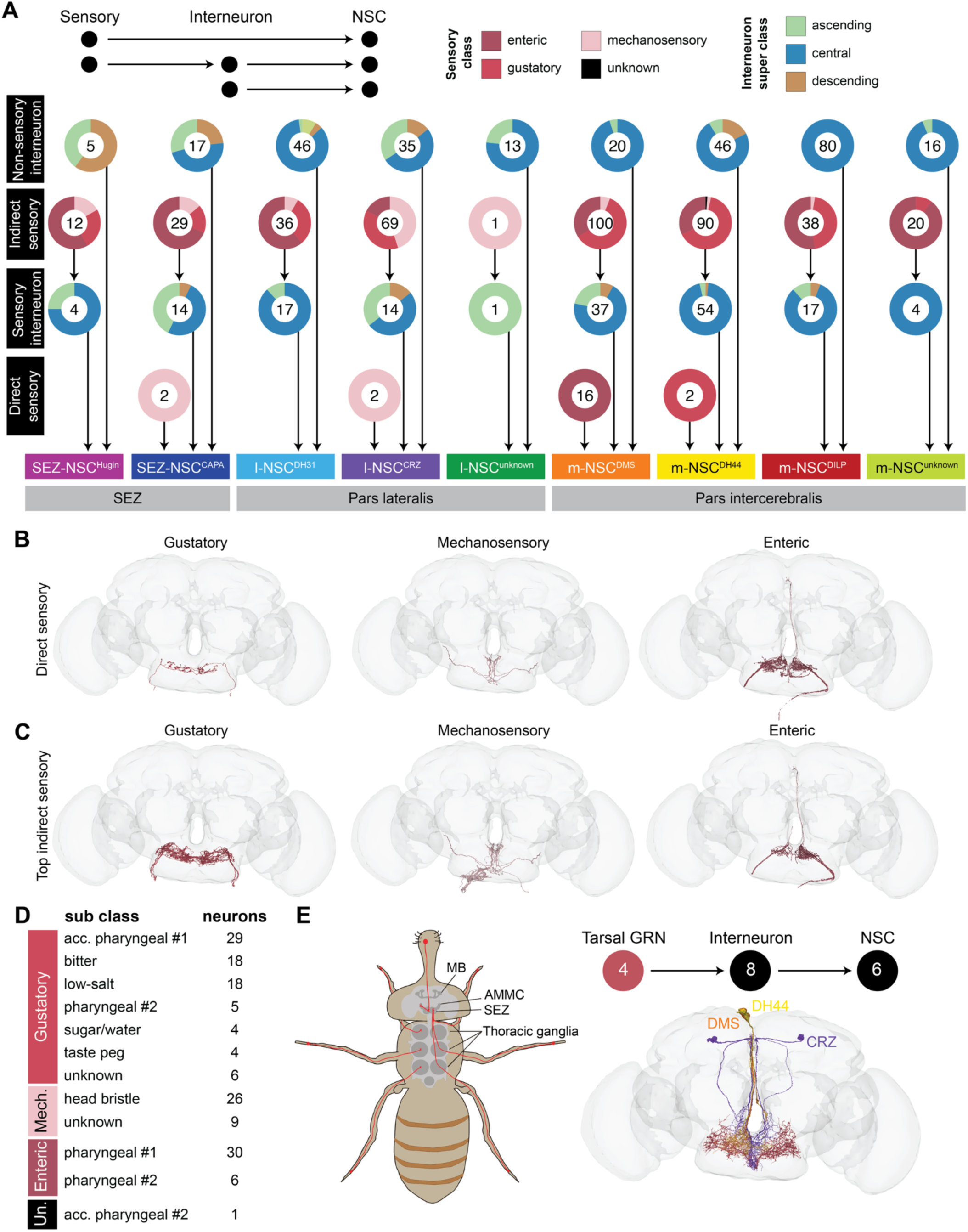
Sensory inputs to NSC in the FlyWire connectome. **(A)** Direct and indirect (disynaptic) sensory inputs to NSC. Interneurons mediating connectivity between sensory neurons and NSC are referred to as sensory interneurons. The donuts represent the proportion of cells and the number in the donut reflects the total number of neurons in that group. NSC receive very minimal direct sensory inputs. Gustatory, mechanosensory and enteric neurons provide the majority of the monosynaptic and disynaptic inputs to NSC. Note that l-NSC^ITP^ do not receive any significant synaptic inputs and are thus not represented here. **(B)** Reconstructions of sensory neurons (separated by class) providing direct inputs to NSC. **(C)** Reconstructions of sensory neurons providing indirect inputs to NSC. **(D)** Number of sensory neurons (grouped by sub class) that provide indirect inputs to NSC. **(E)** Schematic showing the projections of labellar and tarsal gustatory receptor neurons (GRN) from the periphery to the SEZ (adapted from (Freeman and Dahanukar, 2015)). Reconstructions of four tarsal GRN (colored red; classified as ascending neurons on Codex) that provide indirect inputs to six NSC (also shown). Abbreviations: acc. pharyngeal, accessory pharyngeal; mech, mechanosensory; un, unknown sensory.

Next, we examined indirect sensory inputs to each NSC type. Almost all types of NSC receive disynaptic inputs from GRN, enteric and mechanosensory neurons, with m-NSC^DMS^ and m-NSC^DH44^ receiving inputs from the largest number of neurons **(Figure 4A and 4C).** Although l-NSC^unknown^ is the second largest cluster after m-NSC^DILP^, they hardly receive any input from sensory neurons **(Figure 4A)**. Looking further at the categories of sensory neurons, NSC receive inputs from all the major sub-classes of GRN as well as mechanosensory neurons in the head bristle **(Figure 4D)**. The latter neurons likely regulate grooming behavior (Zhang *et al*., 2020); however, the link between grooming and endocrine signaling remains to be explored. Further, only 4 labellar sugar/water taste neurons are disynaptically connected to NSC. We expected sweet taste to be a major regulator of insulin, DH44, and CRZ signaling since these hormones have known roles in feeding and glucose homeostasis (Dus *et al*., 2015; Kubrak *et al*., 2016; Nässel and Zandawala, 2019). This prompted us to explore pathways from external GRN in other structures such as the legs **(Figure 4E)** (Freeman and Dahanukar, 2015). We identified 4 tarsal GRN in the connectome based on anatomical similarity (Thoma *et al*., 2016). Although tarsal GRN are not directly connected to NSC, they do provide indirect inputs to l-NSC^CRZ^, m-NSC^DH44^, and m-NSC^DMS^ **(Figure 4E)**. Hence, tarsal and labellar GRN could regulate NSC activity to some extent. Intriguingly, the majority of the taste-related input to NSC stems from enteric neurons (i.e., neurons in the hypocerebral ganglion which are associated with the anterior gut (Min *et al*., 2021)) as well as neurons projecting via the accessory pharyngeal nerve **(Figure 4D)**. This suggests that internal taste neurons in the pharynx and those associated with the gut, which are activated upon feeding rather than by external taste, are more important for neurosecretion. Interestingly, gustatory, mechanosensory, and enteric neurons are the only sensory neurons disynaptically upstream of NSC. Therefore, other sensory modalities such as vision and olfaction require additional layers of connectivity.

We next focused on olfactory pathways to NSC because pheromones and odors can have a profound impact on hormonal activity and resultant physiology (Lushchak *et al*., 2015; He *et al*., 2022; Zhang *et al*., 2022). For instance, acute exposure to food odors alone can trigger an anticipatory endocrine response (Lushchak *et al*., 2015). To obtain novel insights into olfactory modulation of NSC activity, we explored the shortest path from olfactory receptor neurons (ORN) to NSC. The canonical olfactory pathway in *Drosophila* and other insects begins with the ORN in the antennae and maxillary palps **(Figure 5A)**. ORN of a given type all project to a single glomerulus in the antennal lobe (Vosshall *et al*., 2000). From here, uni-glomerular and multi-glomerular projection neurons (PN) transmit olfactory information to higher-order brain centers such as the mushroom body for learning and memory and the lateral horn which controls innate behaviors (Schlegel *et al*., 2021). Since NSC are not located within these brain regions, additional interneurons likely transmit the information from PN to NSC. In addition, local interneurons (LN) in the antennal lobes innervate multiple glomeruli and modulate olfactory pathways. Our analysis identified a total of 321 ORN which provide inputs to 8 NSC via this canonical pathway **(Figure 5B)**. Moreover, we grouped the different ORN based on their behavioral significance (Zheng *et al*., 2022). ORN such as V, DL4 and DL5 that detect aversive odors comprise the largest group which provides input to NSC **(Figure 5B-E)**. Food odor-related ORN such as DP1l, VA6, DL2d, VL2a, and VM7d make up the second largest group **(Figure 5B-E)**. Olfactory information from the antennal lobe is relayed via 10 uni-glomerular PN and 3 multi-glomerular PN **(Figure 5F and 5G)**. Interestingly, while all SEZ-NSC^CAPA^ are indirectly downstream of ORN, only 2 out of the 18 m-NSC^DILP^ and 2 out of the 6 l-NSC^DH31^ receive olfactory inputs **(Figure 5H)**, further emphasizing the heterogeneity within these clusters. Intriguingly, 11 pheromonal and 2 egg-laying related ORN provide input to m-NSC^DILP^ via this 3-hop pathway **(Figure 5I)**. This pathway is distinct from a male-specific pathway identified earlier whereby pheromonal inputs from the leg ppk23 neurons activate m-NSC^DILP^ to inhibit courtship drive (Zhang *et al*., 2022). In addition, the strongest olfactory inputs are directed to l-NSC^DH31^ **(Figure 5J)**. These inputs stem from all types of ORN, with aversive and food-related ORN providing the majority of inputs. Since DP1l and M_lvPNm35 express the excitatory neurotransmitter acetylcholine, food odors likely promote the release of DH31 and ITP from l-NSC^DH31^ (Gera *et al*., 2025). Since both hormones, albeit from different sources, have previously been implicated in feeding (Lin *et al*., 2022; Gera *et al*., 2025), l-NSC^DH31^ could also have a complementary role in feeding-related behaviors and physiological processes. Lastly, SEZ-NSC^CAPA^ also receive strong inputs from aversive ORN via cholinergic PN and GABAergic LHPV10c1 interneurons **(Figure 5K)**. Although our analysis was based on the shortest 3-hop pathway, if we consider an additional layer of neurons and account for 4 hops, more than 91% of all ORN provide input to m-NSC^DILP^, l-NSC^DH31^ and SEZ-NSC^CAPA^ (not shown). At this level of connectivity, where most ORN are connected to NSC, it is difficult to decipher specific pathways. This is not surprising since the average shortest path length between any two neurons in the entire connectome is about 4 hops (Lin *et al*., 2024). In summary, olfactory inputs to NSC appear to be relatively sparse, and pheromonal and aversive odors could modulate hormonal signaling.

**Figure 5:**
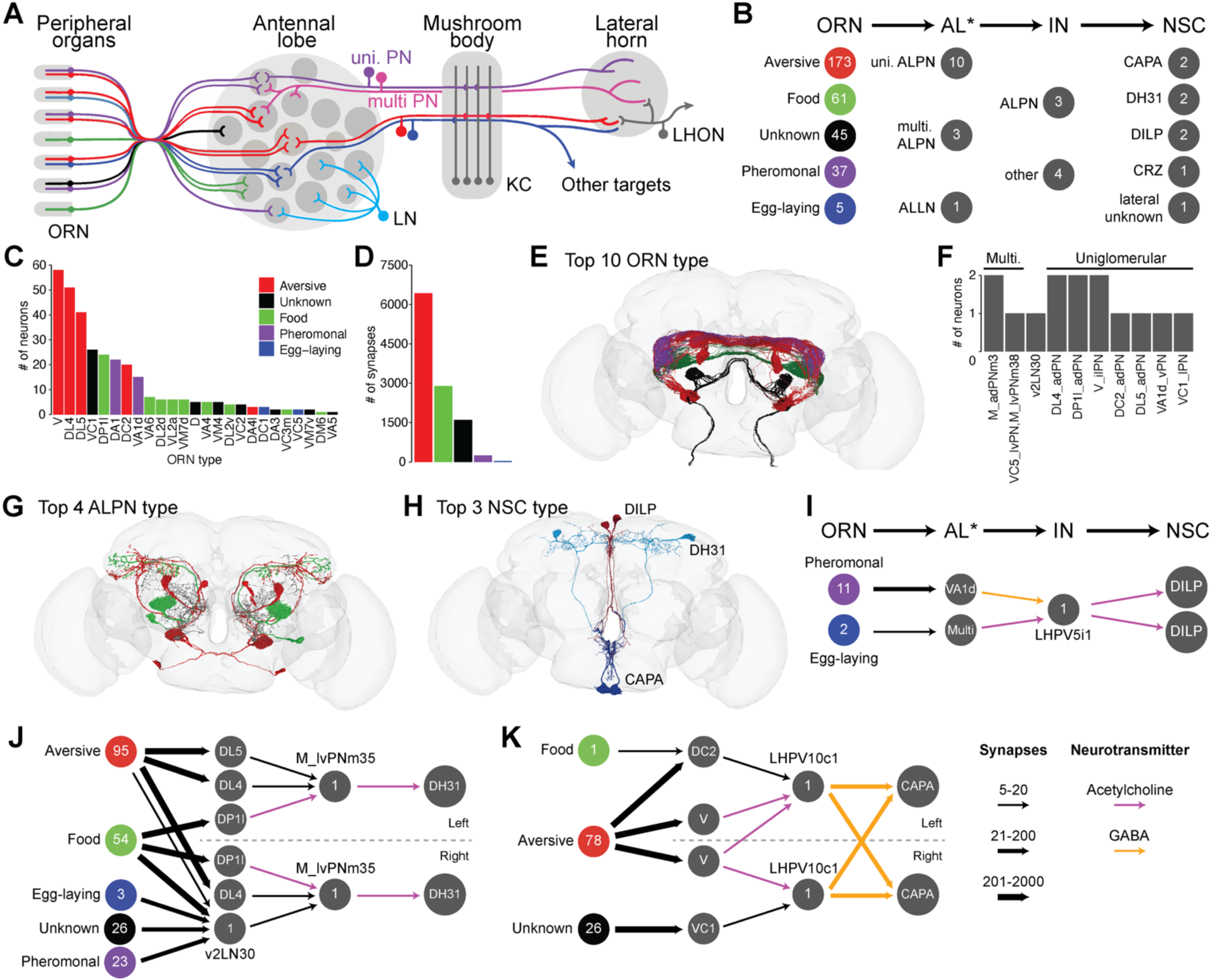
Olfactory inputs to NSC. **(A)** Schematic showcasing the flow of olfactory information from olfactory receptor neurons (ORN) in the antenna to the higher-order brain centers (e.g. mushroom bodies and lateral horn) via the antennal lobe (adapted from (Zhao and McBride, 2020)). **(B)** Number of neurons (grouped by different categories) that comprise the shortest pathway from ORN to NSC. ORN have been grouped based on their behavioral significance (based on (Zheng *et al*., 2022)). Antennal lobe associated neurons (AL*) include projection neurons (ALPN) and local interneurons (ALLN). IN represent interneurons that link AL* neurons and NSC. **(C)** Numbers of each ORN type that provide indirect inputs to NSC. **(D)** Number of synapses formed by these ORN. Note that the ORN which detect aversive odors followed by those that detect food odors provide the strongest indirect inputs to NSC. **(E)** Reconstructions of top ten ORN types. **(F)** Number of AL* in the pathway. v2LN30 is the only ALLN whereas the rest are ALPN. **(G)** Reconstructions of top four ALPN types and **(H)** top three NSC classes that are part of this pathway. Bars in (C) and (D) and neurons in (E) and (G) have been colored based on their behavioral significance. **(I)** Pheromonal and egg-laying associated olfactory information is relayed to m-NSC^DILP^. **(J)** ORN belonging to all five behavioral categories provide inputs to l-NSC^DH31^. **(K)** SEZ-NSC^CAPA^ primarily receive aversive olfactory inputs. For I-K, the numbers within the circles indicate the number of neurons or the name of that neuron. Arrows have been weighted based on the number of synapses and colored based on the neurotransmitter mediating those connections (see legend). Abbreviations: LN, local interneuron; uni. PN, uniglomerular projection neuron; multi. PN, multiglomerular projection neuron; KC, Kenyon cell; LHON, lateral horn output neuron.

### Enteric neurons have a stronger influence on NSC compared to other sensory neurons

While a circuit diagram depicts which neurons are connected and how strongly (i.e. the number of synapses), they cannot assess the impact of one set of neurons on another set. This is especially true for neuron sets that are connected indirectly via multiple synaptic pathways. Moreover, specific synaptic inputs to a given neuron need to be weighted in relation to the receiving neuron’s total synaptic input to obtain a more accurate representation of the impact on the receiving neuron. Thus, to obtain further insight into sensory pathways regulating NSC activity, we used a recently established linear dynamical modeling-based approach (Bates *et al*., 2026) to quantify the influence of various sensory source neurons on different NSC cell classes **(Figure 6A)**. First, we stimulated different types of sensory neurons including gustatory, olfactory, mechanosensory, enteric, hygrosensory and thermosensory neurons in the FlyWire dataset. Consistent with our olfactory pathway analysis **(Figure 5J-K)**, SEZ-NSC^CAPA^ and l-NSC^DH31^ are influenced by various types of olfactory inputs, and more strongly than other types of NSC **(Figure 6B)**. However, olfaction has a much weaker influence on NSC in relation to peripheral gustatory neurons and enteric neurons. Enteric neurons, specifically ENS4, have the strongest influence on m-NSC^DMS^, consistent with their direct connectivity **(Figure 4A)**. Hygro-and thermo-sensation predominantly influence SEZ-NSC^CAPA^, consistent with the role of CAPA in modulating desiccation and cold tolerance (Terhzaz *et al*., 2015). Since influence scores are relative rather than absolute values, we next assessed the influence of sensory inputs on NSC in relation to other cell types **(Figure 6C-D)**. In relation to other cell types in the olfactory pathway (e.g. antennal lobe projection neurons and Kenyon cells) **(Figure 5A)**, the influence of olfaction on NSC is quite low **(Figure 6C)**. However, enteric neurons, especially ENS4, do have a strong influence on m-NSC, and this is ranked top in relation to their influence on other cell types **(Figure 6D)**. We also observed similar trends when examining sensory to NSC pathways in the BANC connectome **(Figure 6 Supplement 1)**. Taken together, this analysis provides a framework to generate testable hypotheses regarding novel functional connections between sensory neurons and NSC.

**Figure 6:**
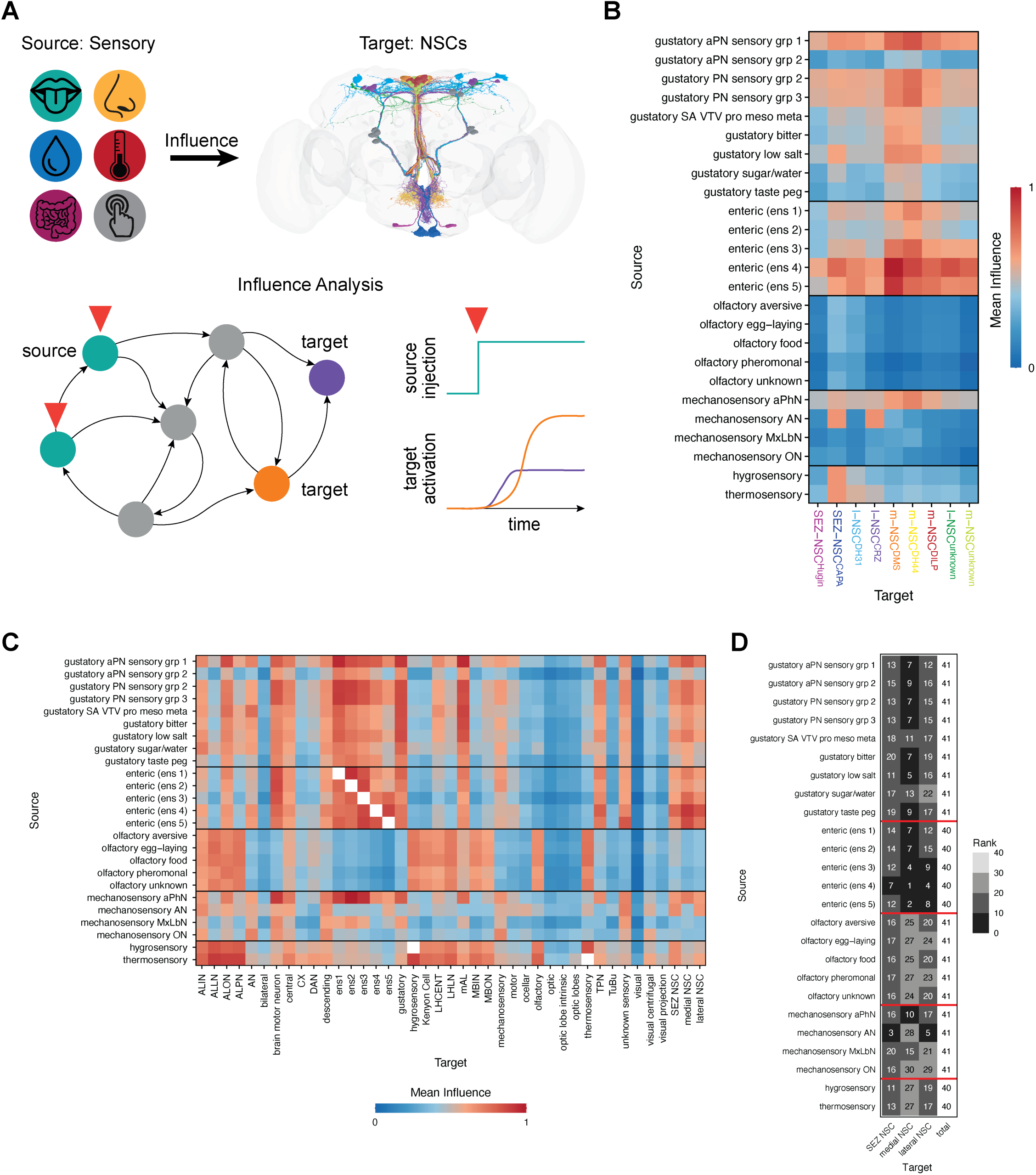
Influence of *in silico* sensory neuron stimulation on NSC in the FlyWire connectome. **(A)** A schematic of the linear dynamical modeling-based approach (Bates *et al*., 2026) used to calculate the ‘influence’ of sensory source cells on target NSC. **(B)** Mean influence of different sensory neurons (rows) on NSC classes (columns). Note the strong influence of enteric neurons on NSC, especially m-NSC^DMS^. **(C)** Mean influence of different sensory neurons (rows) on major cell types in the brain, including NSC. **(D)** The mean influence in panel (C) was used to determine the rank of influence on NSC in relation to other cell types. Enteric neurons, especially ENS4, have the strongest influence on medial NSC.

### NSC have sparse synaptic output

While NSC predominantly release hormones into the circulation following their activation, some NSC types can also signal to other neurons within the brain (King *et al*., 2017). With this in mind, we examined synaptic output from NSC. Output synapses have been predicted for most NSC in the FlyWire connectome, with l-NSC^CRZ^ and l-NSC^unknown^ having the largest number **(Figure 7A and Figure 7 Supplement 1)**. Similar to the location of their dendrites, these output synapses are situated in the protocerebrum and SEZ. Although all types of NSC form output synapses, most of these comprise a connection which does not meet our threshold of 5 synapses. Hence, l-NSC^CRZ^ and l-NSC^unknown^ are the only NSC types which provide significant output to other neurons within the brain volume **(Figure 7B)**. The output from l-NSC^unknown^ is primarily directed to cholinergic and glutamatergic central neurons **(Figure 7B-D and Figure 7G)** whereas two pairs of l-NSC^CRZ^ provide strong output to DNg27 descending neurons **(Figure 7B-C and Figure 7E-F)**. DNg27 neurons, whose function is yet unknown, contain DCVs and innervate the SEZ, intermediate tectulum, lower tectulum and wing tectulum (Court *et al*., 2020). This suggests that they could regulate various functions including feeding and flight **(Figure 7E-F)** through synaptic and paracrine signaling. We also explored the output from NSC after lowering the threshold of significant connections to 2 synapses **(Figure 7 Supplement 2)**. Although the output from NSC increases substantially at this threshold, most of this output is directed to undefined cells which include partial fragments as well as non-neuronal cells **(Figure 7 Supplement 2A-C)**. Nonetheless, additional output to central, endocrine, and descending neurons is also observed, some of which could be biologically significant even though only a few synapses mediate these connections. To determine if any of these connections are biologically relevant, we also explored synaptic output from NSC in BANC **(Figure 7 Supplement 3A-B)** and maleCNS **(Figure 7 Supplement 3C-D)** datasets. In BANC, the only connection observed is that between l-NSC^CRZ^ and DNg27 **(Figure 7 Supplement 3A-B)**. The l-NSC^CRZ^ to DNg27 connection is also present in maleCNS, but additional connectivity between NSC and central brain neurons, as well as NSC interconnectivity, is observed **(Figure 7 Supplement 3C-D)**. It remains to be seen if this additional connectivity observed in the maleCNS connectome compared to the FlyWire and BANC connectomes (both from a female fly) is due to sexual differences and/or differences in synapse prediction accuracy across datasets. In conclusion, sparse synaptic output from NSC agrees with the expectation that they mainly signal in a paracrine and endocrine manner. Since l-NSC^CRZ^ to DNg27 is the only connection common across the datasets, we validated the functions of these neurons.

**Figure 7:**
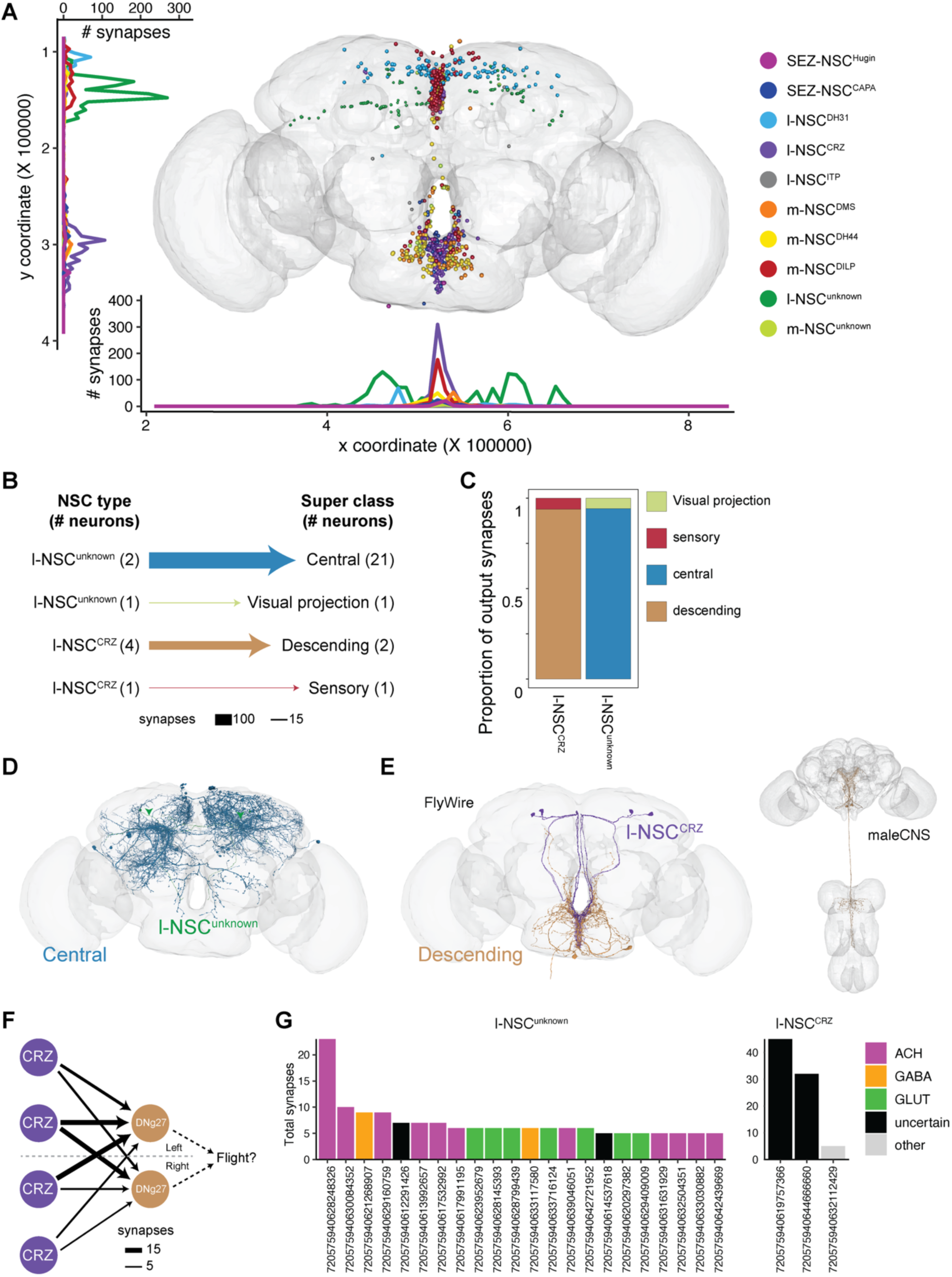
Synaptic output from NSC in the FlyWire connectome. **(A)** Presynaptic sites of different NSC classes. **(B)** Output from NSC grouped by the neuronal super classes annotated in the FlyWire connectome. Central neurons receive inputs from l-NSC^unknown^ and descending neurons receive inputs from l-NSC^CRZ^. **(C)** Proportion of outputs from different NSC classes to various neuronal super classes. **(D)** Reconstructions of l-NSC^unknown^ and all their postsynaptic partners. **(E)** Reconstructions of l-NSC^CRZ^ and all their postsynaptic partners (DNg27 descending neurons) in the FlyWire connectome. DNg27 reconstructions in the maleCNS connectome show that these neurons innervate the wing tectulum. **(F)** Weighted connections between l-NSC^CRZ^ and DNg27 descending neurons which innervate wing power motor neurons in the wing tectulum and could thus regulate flight. **(G)** Individual postsynaptic partners of l-NSC^unknown^ and l-NSC^CRZ^ sorted based on the number of synapses and colored based on their neurotransmitter identity.

### l-NSC^CRZ^ and DNg27 regulate egg-laying

Previous work has shown that l-NSC^CRZ^ co-express CRZ and short neuropeptide F (sNPF) neuropeptides (Kapan *et al*., 2012), and these neurons regulate feeding, metabolism and osmotic homeostasis (Kubrak *et al*., 2016; Oh *et al*., 2019; Zandawala *et al*., 2021). However, these effects are primarily mediated via modulation of other hormonal pathways or peripheral tissues. Our connectome analysis here reveals that l-NSC^CRZ^ also exhibit synaptic output in addition to their known roles via volume transmission. Since the synaptic output from l-NSC^CRZ^ is targeted to DNg27 descending neurons with unknown function, we tested the function of both cell types in various behaviors and physiology. We examined behaviors that are primarily regulated by the brain (i.e. feeding) as well as those that are governed by the VNC (i.e. flight and egg-laying). Genetically silencing all CRZ neurons including l-NSC^CRZ^ with Kir2.1, an inwardly rectifying potassium channel, resulted in reduced food intake **(Figure 8A)** and increased starvation survival **(Figure 8B)**. Hence, our genetic manipulation of l-NSC^CRZ^ can recapitulate the phenotypes observed following reduced CRZ signaling (Kubrak *et al*., 2016). In contrast, silencing DNg27 neurons had no effect on food intake **(Figure 8D)** or starvation survival **(Figure 8E)**. Thus, the effect of l-NSC^CRZ^ on feeding and metabolic physiology appears to be independent of DNg27.

**Figure 8:**
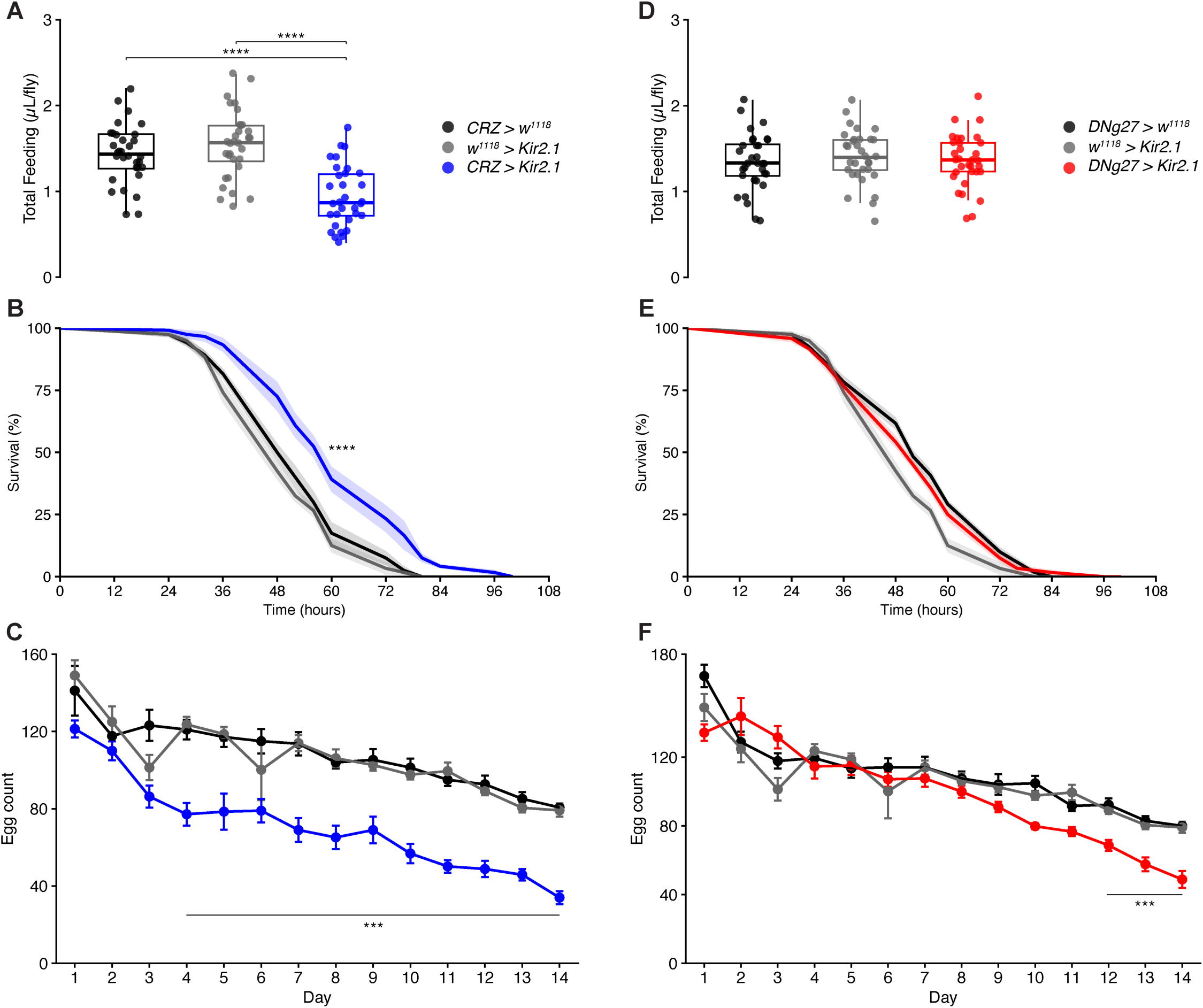
Functional characterization of Corazonin (CRZ) and DNg27 descending neurons. Chemogenetic silencing of CRZ neurons using *Kir2.1* results in **(A)** reduced food intake, **(B)** increased starvation survival and **(C)** a reduction in the number of eggs laid. In contrast, silencing DNg27 neurons has no effect on **(D)** food intake and **(E)** starvation survival but also results in **(F)** a reduction in the number of eggs laid. In panels (B) and (E), lines represent mean percent survival curve, with shaded bands representing ± 1 standard error of the mean (SEM). Error bars in panels (C) and (F) indicate SEM. For panels (A) and (D), **** p < 0.0001, as assessed by one-way ANOVA followed by Tukey’s HSD for multiple comparisons. For panels (B) and (E), **** p < 0.0001, as assessed by log-rank (Mantel-Cox) test, followed by Bonferroni correction for multiple comparisons. For panels (C) and (F), *** p < 0.001, as assessed using a linear mixed effects model, followed by Tukey-adjusted pairwise comparisons.

Research in *Drosophila* and other insects has also implicated CRZ or CRZ-expression neurons in various reproductive behaviors (Bergland *et al*., 2012; Xi *et al*., 2025; Zhang *et al*., 2026). In addition, male-specific CRZ neurons in the ventral nerve cord affect fertility by regulating sperm transfer to females during mating (Tayler *et al*., 2012). Therefore, we asked whether female CRZ neurons also regulate fertility, and if this is dependent on DNg27. To address this, we mated experimental or control females flies with wild-type males and monitored the number of eggs laid over two weeks. Females with genetically-silenced CRZ neurons lay less eggs compared to their genetic controls starting from day 4 **(Figure 8C).** Similarly, females with silenced DNg27 neurons also lay less eggs; however, this effect is only visible from day 12 onwards **(Figure 8F)**. This suggests that CRZ from l-NSC^CRZ^ regulates egg-laying, and this effect may partially be mediated via DNg27 neurons.

Finally, our connectome analysis suggests that DNg27 neurons could regulate flight, since they innervate wing power motor neurons. We utilized an optogenetic strategy established previously (Stupski and van Breugel, 2024) to test the role of both CRZ and DNg27 neurons in a free-flight wind tunnel **(Figure 8 Supplement 1A)**. During free-flight, activation of neither CRZ nor DNg27 neurons in female flies demonstrated significant alterations in flight trajectory parameters, including course direction, horizontal velocity, and flight altitude during or after optogenetic illumination, compared with no-LED flash sham controls **(Figure 8 Supplement 1D-F)**. Trajectory parameters were overall similar to those of the corresponding *empty-Gal4* and *empty split-Gal4* genetic controls. *CRZ-Gal4* females did demonstrate a slightly increased startle response during red LED stimulation, with a small increase in trajectories oriented towards π rad/180° and a larger peak in horizontal velocity 100-200 milliseconds after illumination onset **(Figure 8 Supplement 1D-E,** top rows). CRZ neurons express other neuropeptides and acetylcholine (Oh *et al*., 2019) and *CRZ-GAL4* also drives strong expression in the optic lobe neurons **(Figure 8 Supplement 2)**. Thus, release of multiple neuronal signaling molecules following stimulation with *CRZ-GAL4* may increase a visual startle effect often observed during optogenetic activation in the flight chamber.

We independently confirmed that activation of CRZ neurons in the abdominal ganglion of males can recapitulate copulation-like behaviors **(Figure 8 Supplement 1B-C)** reported previously (Inagaki *et al*., 2014). Optogenetic activation of CRZ neurons in males elicited significantly more copulatory-related behaviors (genital eversion with or without abdominal curling and targeted genital grooming) than genetic controls **(Figure 8 Supplement 1C**). Hence, the lack of an observable phenotype following CRZ neuron activation during free flight is likely not due to insufficient activation. However, we cannot rule this possibility out for DNg27 since no functions are currently known for these neurons, and the driver targeting DNg27 drove CsChrimson-mVenus expression very weakly **(Figure 8 Supplement 2)**.

Together, our functional characterization of CRZ and DNg27 neurons identifies a novel pathway via which l-NSC^CRZ^ could regulate egg-laying in females. Since l-NSC^CRZ^ to DNg27 connections also exist in males, these neurons could regulate other behaviors besides egg-laying.

### Peptide hormone receptors are broadly expressed in NSC and peripheral tissues

The availability of large-scale single-cell transcriptome datasets now enables us to identify and explore transcriptomes of rare cell types such as NSC. We have recently used this strategy to determine the modulatory inputs to m-NSC^DILP^ (Held *et al*., 2025), l-NSC^ITP^ (Gera *et al*., 2025), and some other NSC types (Reinhard *et al*., 2024). Here, we expand this approach to first identify single-cell transcriptomes of all NSC classes based on previously established markers **(Figure 9A)**. Consistent with our anatomical mapping **(Figure 2D)**, *Capa* and *Ms* are co-expressed in SEZ-NSC^CAPA^. Given the proximity of all NSC axon terminations, it is extremely likely that a hormone released from a given NSC will influence the activity of other NSC types if its receptor is expressed in those cells. In fact, we have previously shown CRZ to inhibit CAPA release from a different set of neurosecretory cells much further away in the ventral nerve cord (Zandawala *et al*., 2021). Therefore, we examined the expression of hormone receptors in all the NSC transcriptomes to determine the molecular substrates for paracrine interaction between different NSC types **(Figure 9B)**. Consistent with previous studies, *CrzR* is expressed in SEZ-NSC^CAPA^ (Zandawala *et al*., 2021), *sNPF-R* and *Lkr* in m-NSC^DILP^ (Kapan *et al*., 2012; Zandawala *et al*., 2018c) and *Dh44-R2* in SEZ-NSC^Hugin^ (Mizuno *et al*., 2021). While receptors for CAPA and ITP were not detected in any transcriptomes, receptors for other hormones were expressed in varying amounts. l-NSC^DH31^ express receptors for several hormones and thus appear to be heavily modulated **(Figure 9B)**. Further, *InR* (insulin receptor) is expressed in all NSC classes consistent with the role of insulin in promoting cellular glucose uptake (O’Neill, 2013). Having mapped the expression of peptide hormones and their receptors in different types of NSC, we sought to explore the strength of putative paracrine connections based on their expression levels. Thus, higher expression of both the hormone and receptor implies a stronger connection. Using this approach, we show the extent of putative paracrine connectivity between different hormonal systems **(Figure 9C and Figure 9 Supplement 1)**. We used a conservative approach (see methods for details) to reduce false prediction by setting a stringent expression threshold for the hormone levels **(Figure 9 Supplement 1I)**. Our analysis reveals that paracrine signaling can greatly enhance the interactions between different hormonal pathways.

**Figure 9:**
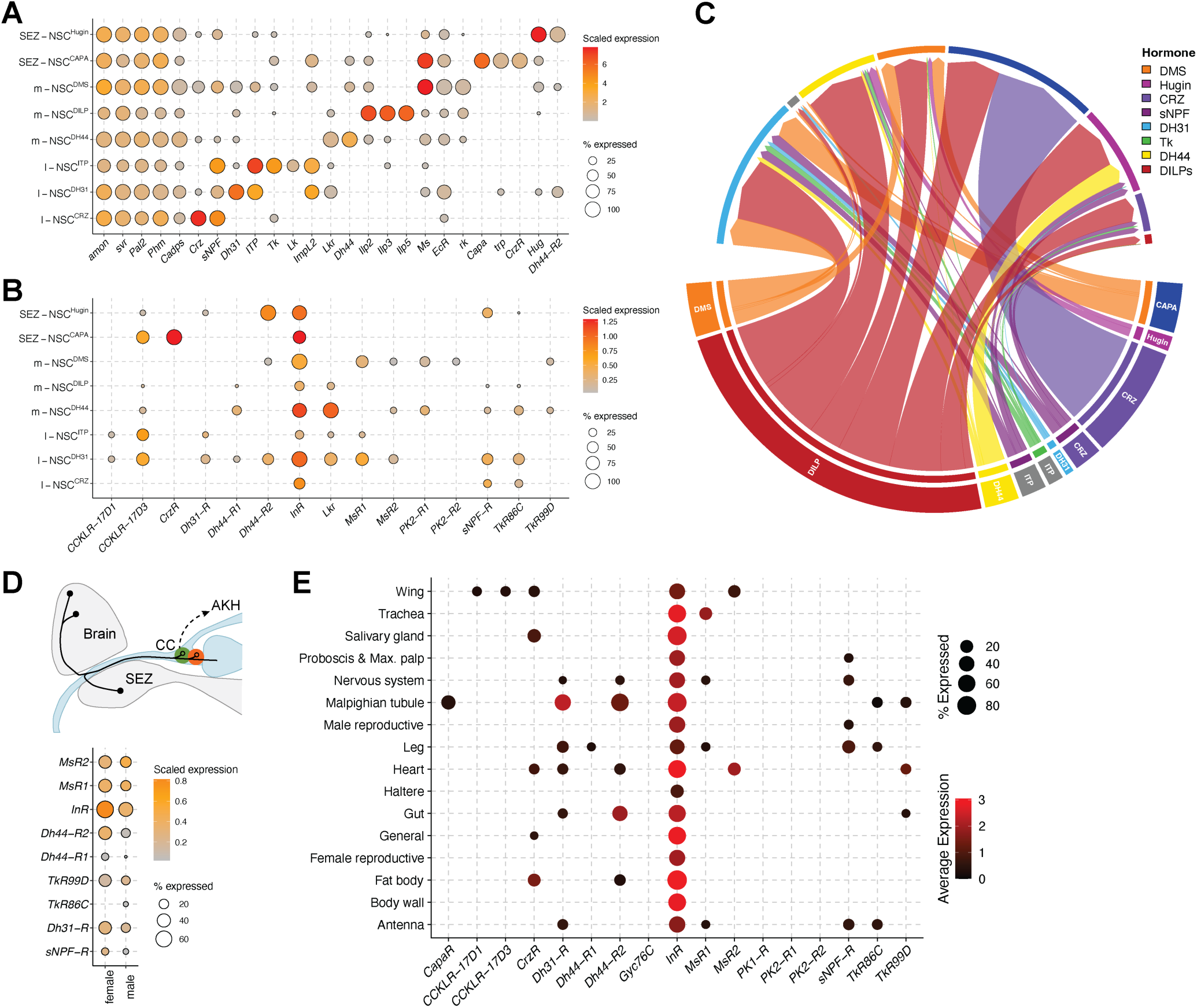
Putative NSC interconnectivity and endocrine output. **(A)** Identification of single-cell transcriptomes representing different NSC subsets in the adult brain (Davie *et al*., 2018). All NSC express genes required for neuropeptide processing and release (*amon*, *svr*, *Pal2*, *Phm* and *Cadps*) and were identified primarily based on the neuropeptides that they express. **(B)** Dot plot showing expression of receptors in NSC. Expression of only those receptors whose corresponding neuropeptides are expressed in NSC are shown. **(C)** Connectivity diagram (weighted based on neuropeptide and receptor expression) showing putative paracrine connectivity between different classes of NSC. Note that short neuropeptide F (sNPF) and myosuppressin (DMS) are expressed in two different NSC classes. Ion transport peptide and CAPA pathways are not included because their receptors were not detected in these transcriptomes. Leucokinin was excluded because its expression levels were below the threshold used here. Dot plot showing the expression of neuropeptide receptors in **(D)** adipokinetic hormone cells of the corpus cardiacum and **(E)** all the tissues in adults. “General” in panel (E) includes cell types that are found across multiple tissues including sensory neuron, visceral muscle and hemocytes amongst others. See Figure 9 Supplement 3 for all the different cell types that are part of this cluster.

Since all NSC have their release sites on or near adipokinetic hormone (AKH) producing cells of the CC, various hormones can influence the release of AKH (Oh *et al*., 2019; Koyama *et al*., 2021). Glucagon-like AKH, along with DILPs, is a major regulator of metabolic homeostasis and associated behaviors (Isabel *et al*., 2005; Yu *et al*., 2016; Nässel and Zandawala, 2019). Consequently, modulation of AKH release is one way of regulating metabolic physiology. We thus examined single-cell transcriptomes of AKH cells for expression of hormone receptors **(Figure 9D)**. While sNPF-R expression in AKH cells was demonstrated previously (Oh *et al*., 2019), we additionally show the presence of receptors for DMS, DH44, DH31 and tachykinin. Thus, these neuropeptides could regulate metabolic homeostasis via AKH-signaling in addition to their known roles in feeding-related processes (Song *et al*., 2014; Dus *et al*., 2015; Hadjieconomou *et al*., 2020).

Previously, Nässel and Zandawala (2019) catalogued the expression of neuropeptide receptors across all *Drosophila* tissues. However, that analysis was based on a microarray-based dataset (Chintapalli *et al*., 2007) and therefore lacks the sensitivity and resolution offered by current sequencing technologies. We sought to fill this gap by cataloguing the expression of peptide hormone receptors using Fly Cell Atlas, a single-cell transcriptomic resource of all cells of the adult fly. Expression of hormone receptors in salivary glands, nervous system, Malpighian tubules, heart, gut and fat body **(Figure 9E and Figure 9 Supplement 2-4)** is largely consistent with the previous analysis. For instance, expression of *CapaR*, *Dh31-R* and *Dh44-R2* in Malpighian tubules and *CrzR*, *Dh31-R*, *Dh44-R2* and *TkR99D* in the heart was reported previously (Nässel and Zandawala, 2019). Interestingly, we now additionally detect *TkR86C* and *TkR99D* expression in the Malpighian tubules and *MsR2* expression in the heart. Examining expression of these receptors at cellular resolution reveals that *TkR99D* is highly expressed in stellate cells of the Malpighian tubules and *MsR2* is strongly expressed in the alary muscles of the heart **(Figure 9 Supplement 3)**. Insights from a similar analysis were recently used to characterize the effects of tachykinin on stellate cells (Agard *et al*., 2024). Our analysis here suggests that DMS can modulate heart contractility via activation of *MsR2* on the alary muscles. Hence, this approach can uncover novel cellular targets of various hormones. Importantly, Fly Cell Atlas also includes tissues such as trachea, leg, wing, haltere, proboscis and maxillary palp where expression of peptide hormone receptors has not been explored comprehensively. We reveal expression of several receptors including *CrzR*, *MsR1*, *MsR2*, *Dh44-R1* and *Dh44-R2* in sensory neurons of the antenna and other tissues **(Figure 9 Supplement 3)**. Thus, m-NSC^DMS^, m-NSC^DH44^, and l-NSC^CRZ^ not only receive direct sensory inputs **(Figure 4A)** but they could also modulate other types of sensory neurons, potentially forming sensory-endocrine feedback loops. Taken together, our analysis presents an important resource to functionally characterize novel hormonal targets.

## Discussion

Here, we describe the first connectome of a neurosecretory network in the brain of an adult animal. This connectome is based on 72-80 NSC (depending on the connectome) which can be subclassified into 10 categories based on neuropeptide expression, morphological similarity and/or synaptic connectivity **(Figure 10A)**. Moreover, our integration of connectomics with anatomical analyses provides a comprehensive view of the NSC landscape and their connectivity in the adult *Drosophila* brain. Our analyses reveal a functionally diverse, yet highly interconnected neuroendocrine system which provides the basis to perform comparisons with other neuroendocrine connectomes from larval *Platynereis* (Williams *et al*., 2017) and *Drosophila* (Huckesfeld et al., 2021) established previously.

**Figure 10:**
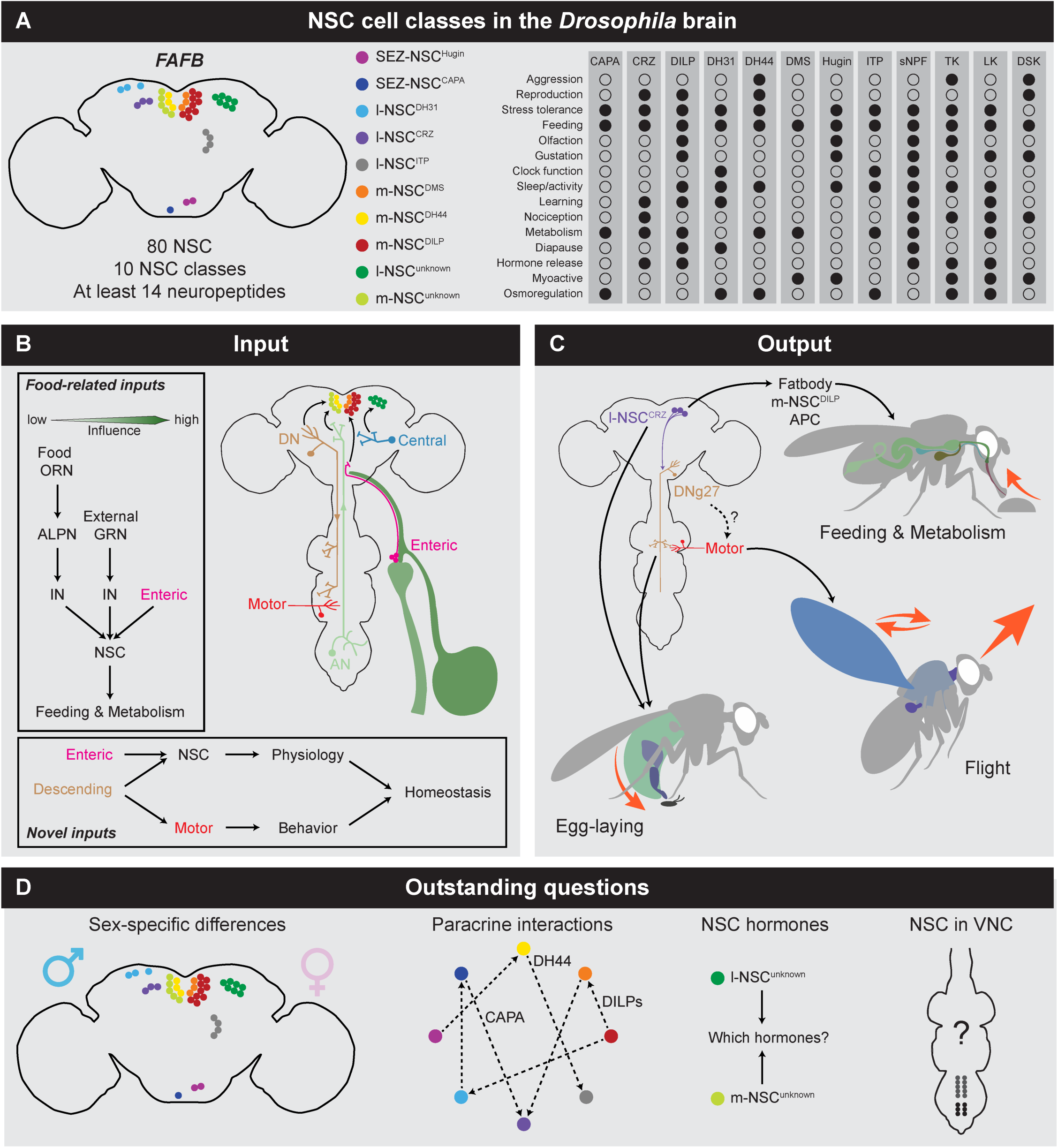
Summary of main findings presented in the study in relation to broader context. **(A)** Schematic showing the different classes of NSC in the FlyWire (or FAFB) connectome as well as the functions regulated by hormones released from these cells. Filled circle indicates that a hormone regulates that function. The right panel is adapted from (Nässel, 2025). **(B)** Major synaptic inputs regulating NSC. With regards to food-related inputs, enteric neurons have a stronger influence on NSC, followed by peripheral GRN and then ORN. Other novel inputs to NSC include descending neurons which could help maintain homeostasis by coordinating physiology (through NSC) with behavior (through motor neurons). **(C)** Major synaptic output from NSC, focusing on l-NSC^CRZ^. l-NSC^CRZ^ target DNg27 and both of these cell types regulate egg-laying. The role of these cells in flight is still unclear. Previous work has shown that l-NSC^CRZ^ regulate feeding and metabolism via multiple targets (Kubrak *et al*., 2016; Oh *et al*., 2019). **(D)** Outstanding questions or future directions include sex-specific differences in connectivity of NSC, functional validation of paracrine interactions between NSC, identification of neuronal messengers in l-NSC^unknown^ and m-NSC^unknown^, and comprehensive identification of NSC in the ventral nerve cord (VNC). Abbreviations: IN, inter neuron; ALPN, antennal lobe projection neuron; GRN, gustatory receptor neuron; ORN, olfactory receptor neuron; AN, ascending neuron; DN, descending neuron; APC, adipokinetic hormone-producing cells.

### Comparison of neuroendocrine connectomes across development, sex and species

The composition of the adult NSC is different compared to larvae. We observed an expansion of m-NSC^DILP^ and m-NSC^DMS^ clusters along with the presence of two additional populations of putative NSC whose identity and function remain to be explored. The expansion in the number of identified NSC classes from larvae to adults underscores developmental changes that might reflect the life-history adaptation of these cells to new physiological demands or environmental challenges. For instance, adults are considerably larger than larvae and comprise more cells in general. Glucose uptake by these cells is modulated by DILPs via InR. Hence, additional m-NSC^DILP^ may be needed for increased DILP production and release to compensate for this size increase. The increase in NSC number is not merely a quantitative change but suggests functional diversification, as evident by the presence of previously unidentified m-NSC and l-NSC populations in adults. The connectome datasets examined here do not include the complete axonal projections of these neurons outside the brain volume. Hence, it is unclear whether these neurons innervate the CC and CA for regulating release of AKH and juvenile hormone, respectively, or if they release hormones into the circulation via neurohemal sites. However, given their smaller size compared to other NSC populations, they might influence the release of other hormones via local actions rather than hormonal regulation of peripheral tissues.

Similar to the *Drosophila* and *Platynereis* larval neuroendocrine connectomes, NSC in adult *Drosophila* have limited synaptic output in the brain (Williams *et al*., 2017; Huckesfeld *et al*., 2021). Eclosion hormone-expressing NSC are the only cells with synaptic output in larvae (Huckesfeld *et al*., 2021). However, these cells undergo apoptosis soon after eclosion and are thus not found in mature adults (Scott *et al*., 2020). In contrast, l-NSC^CRZ^ and l-NSC^unknown^ provide the majority of synaptic output in the FlyWire connectome. l-NSC^CRZ^ are especially of interest here because only 4 out of the 6 neurons in this cluster lie upstream of DNg27 descending neurons that primarily innervate the wing neuropil. The remaining two l-NSC^CRZ^ are internal glucose sensors which signal via sNPF to influence DILP and AKH release, which in turn regulate glucose homeostasis (Oh *et al*., 2019) **(Figure 10C)**. Furthermore, DILPs suppress starvation-induced food search whereas AKH promotes this behavior (Yu *et al*., 2016). Therefore, it is plausible that one subset of l-NSC^CRZ^ affect starvation-dependent locomotor activity via AKH and DILPs, while the other subset modulate flight via DNg27 in response to starvation. However, activation of CRZ neurons or DNg27 did not elicit any obvious phenotypes during free-flight. The function of both neuron types during flight is still unknown. Interestingly, silencing either of these neuron types in females did cause a significant reduction in the number of eggs laid. Since DNg27 do not have direct synaptic output to any known oviposition-regulating neurons in the BANC dataset, they could modulate egg-laying indirectly or via volume transmission. Lastly, the l-NSC^CRZ^ to DNg27 pathway could also regulate other behaviors or physiology besides egg-laying since this circuit motif is also found in males.

In addition to l-NSC^CRZ^, other NSC types including m-NSC^DILP^, m-NSC^DH44^ and l-NSC^DH31^ also exhibit heterogeneity in their morphology, synaptic inputs, and gene expression. This heterogeneity may reflect an adaptive mechanism allowing for fine-tuned responses to environmental and physiological cues. In support of this, only a subset of m-NSC^DILP^ and m-NSC^DH44^ express the mechanosensitive channel Piezo (Wang *et al*., 2020; Oh *et al*., 2021). Similar functional differences between l-NSC^DH31^ subtypes remain to be explored. Regarding non-synaptic output, neuropeptides expressed in the *Platynereis* neuroendocrine center are largely distinct from those found in the *Drosophila* brain NSC, with insulin-like peptides, tachykinin and sulfakinin being the only hormones that are common across both species (Williams *et al*., 2017; Huckesfeld *et al*., 2021). The latter was previously shown to be expressed in a subset of m-NSC^DILP^ (Soderberg *et al*., 2012); however, we were unable to detect it in our single-cell transcriptomic analysis, likely due to low expression. Interestingly, cholecystokinin, the vertebrate ortholog of sulfakinin, is also expressed in the hypothalamus (Williams *et al*., 2017; Nässel and Wu, 2022). Thus, the expression of cholecystokinin/sulfakinin in neuroendocrine centers and their function in regulating satiety are conserved across evolution. Examination of NSC in other species can shed light on other conserved hormonal systems.

### Elucidation of sensory inputs to NSC suggests novel roles for conserved peptide hormones

Our analysis of sensory input pathways revealed that only 20 sensory neurons, primarily enteric neurons, provide direct inputs to NSC in the FlyWire connectome. While this number is much smaller than the corresponding number in larvae (Huckesfeld *et al*., 2021), that study utilized a synaptic threshold of only one. Perhaps one synapse may be sufficient to modulate the activity of NSC on slower timescales. However, this number also includes some transient connections which may not persist due to context-dependent synaptic plasticity. Therefore, we used a higher threshold in line with other studies using the same dataset (Dorkenwald *et al*., 2024; Reinhard *et al*., 2024). To account for multiple indirect pathways between sensory neurons and NSC, we used a linear dynamical modelling approach to calculate the ‘influence’ of sensory neuron stimulation on NSC activity. Our analysis indicates that SEZ-NSC^CAPA^, which express the mammalian neuromedin U homolog CAPA, receive and integrate inputs from multiple sensory modalities. For instance, they receive thermosensory and hygrosensory inputs which could be important for CAPA neuropeptide to regulate desiccation and cold stress tolerance (Terhzaz *et al*., 2015). In addition, they also receive inputs from mechanosensory and olfactory pathways. This cross-sensory integration could allow an animal to comprehensively assess the environment such as during odor-guided navigation where both wind and odor inputs are important. These sensory modalities could also be important during feeding to assess food quality and texture, where gustatory and enteric inputs would also be relevant. Our olfactory circuit tracing analysis also revealed olfactory inputs to l-NSC^DH31^. Interestingly, l-NSC^DH31^ respond to both pheromonal and food-odor inputs. Integration of these odors can attract flies of both sexes to food sources that are suitable for mating (Kohl *et al*., 2015). Based on this circuit motif, we would expect these cells to play a role in feeding, mating and/or courtship. Consistent with this prediction, calcitonin-like DH31 from these neurons targets the CA to suppress juvenile hormone signaling (Kurogi *et al*., 2023), which in turn influences egg maturation, courtship and sex pheromone production (Bilen *et al*., 2013). l-NSC^DH31^ also express ITP which is important for feeding and metabolic homeostasis (Gera *et al*., 2025). Moreover, l-NSC^DH31^ could thus be part of a feedback loop where they receive pheromonal and food-related odor inputs to regulate pheromone production and food intake. Since olfactory inputs have a much weaker influence on all NSC compared to other sensory modalities, we predict that olfactory inputs alone are likely not sufficient to induce or inhibit hormone release **(Figure 10B)**. In comparison to olfaction, internal enteric neurons and peripheral gustatory neurons are predicted to have a greater influence on NSC activity. This is not surprising given that multiple hormones produced by NSC regulate feeding-associated behaviors and metabolic physiology (Kahsai *et al*., 2010; Kapan *et al*., 2012; Dus *et al*., 2015; Yu *et al*., 2016; Oh *et al*., 2019; Gera *et al*., 2025; Qin *et al*., 2026). Importantly, contents of the ingested meal sensed by enteric neurons (Cui *et al*., 2024; Kim *et al*., 2024a) have a stronger influence on NSC compared to inputs from peripheral gustatory receptors. This circuit architecture could ensure that systemic physiological changes orchestrated by hormones are only activated once the food has been consumed. Together, our findings suggest that different NSC classes could integrate information from multiple sensory modalities to regulate crucial interdependent behaviors to regulate systemic homeostasis.

### Interactions between hormonal pathways

Our analyses highlight extensive putative interactions between hormonal systems via both their synaptic inputs and hormonal output. Multiple NSC classes receive inputs from the same set of pre-synaptic neurons, indicating that different hormonal pathways do not operate in isolation but rather interact within a complex network. m-NSC^DH44^ are of particular interest here since they receive the most extensive input from neurons which also provide inputs to other types of NSC. Consequently, circuits which influence other hormones could also affect DH44 release. In addition, m-NSC^DH44^ can function as cell-autonomous glucose and amino acid sensors (Dus *et al*., 2015; Yang *et al*., 2018). m-NSC^DH44^ in females can also integrate inputs regarding the quality of their male partner’s ejaculate by directly sensing a phospho-galactoside present in the male ejaculate (Kim *et al*., 2024b). Further, these cells are also intrinsically mechanosensitive and can monitor the feeding state of the animal based on crop distension (Oh *et al*., 2021). Thus, various inputs regulate the activity of m-NSC^DH44^ and DH44 could act as a major co-coordinator of physiology and behavior. For instance, DH44 could be co-released with DMS and/or DILPs to orchestrate feeding and reproductive processes (Nässel and Zandawala, 2019; Hadjieconomou *et al*., 2020). In addition to the regulation of hormonal pathways by common synaptic inputs, the various hormones can also modulate each other’s release via paracrine signaling. Hence, sulfakinin and DILPs from m-NSC^DILP^ can interact with other signaling pathways as their receptors are expressed in most NSC types. Moreover, l-NSC^DH31^ and AKH-producing cells are also heavily modulated by different hormones. Therefore, in agreement with their major roles in metabolic physiology, several endocrine pathways converge on m-NSC^DILP^ (Held *et al*., 2025) and AKH-producing cells.

### Limitations of our approach

Our connectomics and transcriptomics-based approach to decipher synaptic and paracrine connectivity of the NSC has some general limitations. Firstly, despite providing several lines of anatomical evidence, our assignment of m-NSC^DMS^ and m-NSC^DH44^ cell classes should be treated with caution since the connectome datasets examined here lack molecular markers to identify these cell classes unambiguously. Similarly, the novel synaptic and paracrine pathways identified here are only putative until experimentally verified using functional connectivity or behavioral analyses. Moreover, our analyses underestimate the connectivity for several reasons: 1) the algorithm used for synapse prediction was not 100% effective, 2) we used a fairly stringent threshold (≥ 5 synapses) for assessing connectivity, 3) some neurons have not yet been proofread and 4) NSC could form synapses with each other near their release sites and outside the brain volume examined here. Moreover, NSC can also couple electrically via gap junctions which are not accounted for here (Orchard and Shivers, 1986; Alvarado Alvarez *et al*., 1993). Further, the connectome depicts a static snapshot of connectivity which we anticipate changing with age as well as with the mating and feeding status of the animal. Hence, weaker connections should be treated with caution. However, since we observed similar NSC connectivity across three different datasets, we believe that the inter-individual variability is not drastic when using a threshold of 5 synapses for significant connections. Lastly, we only focus on the brain NSC that project via the NCC. Other brain NSC, namely serotonin-producing DNg28 (Yao and Scott, 2022) and CB2748 cell types, which project via other nerve bundles, are not addressed here.

### Conclusion and future directions

Future research should focus on functional validation of the observed synaptic and paracrine connectivity patterns, particularly how specific sensory inputs to NSC are translated into physiological responses **(Figure 10D)**. Moreover, a more detailed analysis of the neuroendocrine connectome in the male brain can provide insights into sexual dimorphism within the neuroendocrine pathways. Identification and functional characterization of neuronal messengers in novel NSC (l-NSC^unknown^ and m-NSC^unknown^) and comprehensive identification of NSC in the ventral nerve cord can provide a more complete outlook on how the neuroendocrine system integrates different inputs to regulate systemic physiology. Taken together, our characterization of the adult *Drosophila* brain neuroendocrine network connectome provides a foundation to experimentally disrupt endocrine pathways and establish causal relationships with disorders such as diabetes, hypertension, and infertility. In addition, it provides a blueprint for understanding complex hormonal networks and how they orchestrate animal behaviors and physiology.

## Materials and methods

### Fly strains

*Drosophila melanogaster* strains used in this study are listed in **Supplementary Table 1**. Fly lines were mostly obtained from the Bloomington *Drosophila* Stock Center (BDSC). Flies were reared at 25°C under LD12:12 on a standard *Drosophila* medium containing 8.0% malt extract, 8.0% corn flour, 2.2% sugar beet molasses, 1.8% yeast, 1.0% soy flour, 0.8% agar and 0.3% hydroxybenzoic acid. We used 13-15 day old adults for *retro-Tango* analyses.

For CRZ and DNg27 neuron inactivation experiments with Kir2.1, flies were reared on a diet containing 5% w/v cornmeal, 2% w/v yeast, 0.5% w/v agar, 1.35% w/v dextrose, 3% v/v saccharose syrup, 0.75% v/v propionic acid, hydroxybenzoate and 1.125% ethanol.

For the optogenetic testing and validation of *CRZ-Gal4* F1 genotype experiments, parental flies were reared in a humidity-controlled incubator (25°C; 60% relative humidity; LD16:8) in plastic *Drosophila* colony vials (Genesee Scientific Flystuff, #32-117), filled with standard molasses-containing *Drosophila* medium (Genesee Scientific Nutri-Fly MF, #66-44). Crosses between male (*empty-Gal4*, *CRZ-Gal4, empty split-Gal4, DNg27-split-Gal4*), and virgin female (*20xUAS*-*CsChrimson::mVenus*) flies were performed in vials containing the same *Drosophila* medium which also incorporated 0.35 mM all-trans retinal (Sigma; R2500). The *20xUAS-CsChrimson::mVenus* flies were created by injecting the transgene into the attp2 site of wild-type Berlin flies, and have a X chromosome, including a wild-type white gene. Newly eclosed F1 adult flies were transferred to fresh vials containing molasses *Drosophila* medium supplemented by 200 µl of 0.8 mM all-trans retinal deposited on the food surface.

### Immunohistochemistry and confocal imaging

Immunostainings were performed as described previously (Gera *et al*., 2025). Briefly, whole female flies were fixed in 4% paraformaldehyde in phosphate-buffered saline (PBS) with 0.5% Triton X-100 (PBS-T) for 2.5 h on a nutator at room temperature (RT). Fixed flies were washed four times with PBS-T before dissecting their brains. The samples were blocked in PBS-T containing 5% normal goat serum for 1 hour at RT and subsequently incubated in primary antibodies at 4°C for 48 h. Following four washes with PBS-T, the brains were incubated in secondary antibodies at 4°C for 48 h. Lastly, the samples were washed four times in PBS-T and mounted using either Fluoromount-G^TM^ (Invitrogen, Thermo Fisher) or Vectashield mounting medium (Vector Laboratories, Burlingame, CA, USA). Images were acquired using Leica SPE and SP8 confocal microscopes (Leica Microsystems) using 20x or 40x objectives. All images presented in this study are representative images based on at least 5 independent samples.

All primary and secondary antibodies are listed in **Supplementary Table 2**.

### Connectome datasets and NSC identification

We used the v783 snapshot of the FlyWire whole brain connectome and its annotations (annotations last updated: 12 June 2024) for all the analyses (Dorkenwald *et al*., 2024; Schlegel *et al*., 2024). Enteric neurons (ENS1 to ENS5), previously classified as unknown sensory neurons, were recently annotated in the FlyWire connectome. We therefore updated the classification of the relevant unknown sensory neurons to ENS1-5. To identify NSC in the FlyWire connectome, we first identified NSC subsets that have a characteristic morphology and location, e.g. l-NSC^ITP^. We then identified the nerve bundle (nervii corpora cardiaca, NCC) through which l-NSC^ITP^ axons exit the brain. This nerve bundle contains all the axons for NSC in the *Drosophila* brain. We manually assessed other axons in NCC to identify the remaining NSC in the connectome. Independently, we also examined the cross-section of the soma of all putative NSC for the presence of dense core vesicles, which suggests that they contain neuropeptides/neuromodulators.

We used the v888 snapshot of the BANC dataset and its annotations (downloaded from Harvard dataverse or bancr package: 28 May 2026). For maleCNS, we used the v0.9 snapshot of the dataset and its annotations (downloaded from malecns package: 2 April 2026).

Cell IDs of NSC identified in FlyWire, BANC and maleCNS are provided in **Supplementary Table 3**.

### Neurotransmitter predictions

Neurotransmitter predictions for the FlyWire connectome are based on Eckstein *et al*. (2024). We only considered neurotransmitter prediction scores for fast-acting neurotransmitters (i.e., acetylcholine, glutamate, and GABA) that were greater than 62%.

### DCV brightness quantification

The electron micrographs for all m-NSC^DH44^ (6 neurons), m-NSC^DMS^ (6 neurons) and DNp32 descending neurons (2 neurons) from Flywire, BANC and maleCNS connectomes were obtained using neuroglancer (Dorkenwald *et al*., 2022). For consistency, we examined the cross section of each cell where the diameter of nuclei was the largest. We quantified the mean gray value of at least 50 DCVs per cell using the Fiji software. The individual who performed these analyses was blind to the neuron identity.

### Data visualization

Data was visualized using ggplot2 (v 3.5.1, Wickham, 2016) and circlize (v 0.4.16) for R (v 4.4.1) in RStudio (2024.04.2+764) (Gu et al., 2014). FlyWire NSC reconstructions were downloaded using the navis library (v 1.0.4, https://github.com/navis-org) and cloud-volume library (v 8.10.0, https://github.com/seung-lab/cloud-volume) for Python (v 3.8.5), and visualized using blender (v 3.01, Community, B. O. 2018). BANC and maleCNS reconstructions for NSCs were created in R using natverse and rgl libraries. All other neuron reconstructions were visualized using FlyWire neuroglancer (Dorkenwald *et al*., 2022). Links to each visualization in figures are included on Zenodo.

### Connectivity analyses

Natverse libraries (v 0.2.4) for R in RStudio (Bates *et al*., 2020) were used to analyze the connectivity data. NSC were clustered based on all their synaptic inputs with coconatfly (v 0.1.0.9000) for R (Schlegel *et al*., 2024). Enteric neurons in the FlyWire connectome (previously classified as “unknown sensory”) were clustered based on all their synaptic inputs and outputs. Filtered synapses for the FlyWire connectome were retrieved in Python using the navis and pandas (v 1.1.3) libraries. Connectivity data for BANC and maleCNS was retrieved using bancr (v 0.3.2) and malecns (v 0.4.1) packages, respectively. Unless stated otherwise, a threshold of 5 synapses was used to determine significant connections. Our analyses were based on a custom code generated previously (Reinhard *et al*., 2024).

### Influence analysis

Influence analysis was conducted in Python (v 3.13.1) using the connectome influence calculator (v 0.4: https://github.com/DrugowitschLab/ConnectomeInfluenceCalculator).

For analysis using FlyWire, we downloaded the meta data tables (cell types, classification) and unfiltered connections based on Princeton synapse predictions (v783: updated 08 July 2025) from Codex (14 Nov 2025) and used these to run the analysis in Python. Output and results were then processed and visualized in R. We appended any updates from the cell types or classifications from Codex (v783: updated 24 Feb 2026). For analysis using BANC, we downloaded the meta and connectivity (simple_v2) data from the Harvard dataverse (https://dataverse.harvard.edu/dataset.xhtml?persistentId=doi:10.7910/DVN/7WTH1N), using v888 associated with the published manuscript. To match cell types across datasets, we filtered the cell types in BANC according to the sensory cell types we used in FlyWire for the influence analysis.

For both datasets, we followed the analysis pipeline used by (Bates *et al*., 2026). We used a lenient constant c of 24 for adjusted influence that corresponds to the minimum accepted influence of 3.78e-11 and normalized the influence scores by both the number of source and target neurons. We scaled the influence values to be between 0 and 1 for ease of interpretation.

### Starvation survival

To monitor starvation survival, female flies in groups of 20 were placed in glass vials containing 2% agar and maintained at 25°C with 60% relative humidity under a 12:12 LD cycle. Dead flies were quantified visually at regular intervals of 4 hours during daytime until no flies remained alive. Survival curves were generated based on at least 120 flies per genotypes.

### Feeding assay

Capillary feeding (CAFE) assay was performed to measure total food intake as well as to monitor the preference between nutritive and non-nutritive sugars. For total food intake, ad libitum-fed female flies were anesthetized with CO_2_ and individually transferred into 2 ml tubes. A 5 µl capillary tube, filled with 10% sucrose, 10% yeast and 0.1% propionic acid was inserted through a hole into the lid of the 2 ml tubes. Similar tubes without flies were used as controls for evaporation. All the prepared tubes were then kept inside a moist chamber within a standard fly incubator at 25°C on a 12:12 LD cycle. Flies were allowed to feed for 24 hours and the drop in the volume of the food remaining within the capillary was recorded over time. This was then used to calculate the amount of food consumed.

### Egg-laying assay

Virgin females of different genotypes were allowed to mate with wild-type males for 2 days to eliminate any potential effects of genetic manipulation on male fertility. Following mating, females were isolated and transferred to fresh food vials (20 females per vial). Females were flipped into fresh food vials every 24 hours, and eggs were quantified daily for 14 consecutive days. Egg-laying curves were generated based on at least 120 flies per genotype.

### Optogenetic activation in free-flight

The behavioral paradigm was mostly as described previously (Stupski and van Breugel, 2024) with some modifications. Flies were tested in free-flight within a wind-tunnel (1.0 x 0.5 x 0.5 m^3^), housed in a room maintained at 22° C and 60% relative humidity. Ambient light was from blue (STN-BBLU-B6A-08C1M-24V, Super Bright LEDs Inc. St. Louis, MO, USA), white, and infrared (PN: 7031, Waveform Lighting, Vancouver, WA, USA) LEDs. A constant-speed laminar wind was generated continuously with a measured wind speed of 0.4 m/s, calibrated using a hot-wire anemometer (Alnor Velometer AVM410; TSI Inc.). When flies entered a defined trigger volume, one of two events was triggered at random: either a flash event, during which red LEDs (Triple Output High Power RGB LEDs, COM-15200, Sparkfun.com; λ = 620 nm) were activated for 2000 milliseconds, or a control sham event without any additional LED activation. Regardless of event type, the trajectories of each fly were recorded. The trigger volume dimensions differed somewhat from (Stupski and van Breugel, 2024), having larger dimensions along the tunnel length, width, and height, resulting in a greater number of recorded trajectories. Comparing sham controls with responses to flashes was used to determine the effects of the activation of CRZ and DNg27 neurons on free-flight characteristics.

The total illumination of the tunnel chamber was measured along the midline (y-axis) at multiple points across the length of the tunnel (x-axis) and at multiple altitude levels (z-axis). Illumination was measured using a light meter (Dr. meter LX 1330B) positioned horizontally to measure light coming from above the sensor, which includes all of the red LEDs. As lux values across the x-axis positions did not vary significantly, we averaged along the tunnel length, resulting in mean lux values for red, white, and blue LEDs at the 3 z-axis positions: 13, 23, and 33 centimeters above the tunnel floor. For each LED type, we converted the lux values to mean irradiance *E_irr_* (μW/mm^2^) using the equation:

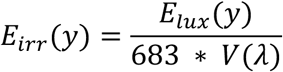

*E_lux_*(*y*) is the mean lux values at z-position *y*, and 683 ∗ *V*(*λ*) represents the luminous efficacy, which is the efficiency of conversion of a light source’s power to visible light, consistent with human cone physiology (CVRL: http://www.cvrl.org/lumindex.htm). Adding the irradiance for individual LED types, the ambient irradiance (white and blue LEDs) ranged from 4.8 - 10.7 μW/mm^2^ from the lowest to highest altitudes, respectively, whereas ambient light plus optogenetic stimulation from red LEDs varied from 12-52 μW/mm^2^.

The morning of an experiment, female F1 flies (2-5 days old) were anesthetized at 4°C on a cold plate, and 20-26 females were placed in an empty tube with a moist Kimwipe to prevent desiccation. We then starved flies in their preparation tubes for 6-8 hours in darkness prior to any experimentation to encourage flight and food seeking. In the evening (16-19:00) we placed experimental flies into the wind tunnel behavioral system without any further anesthesia. Flies were permitted to fly about the tunnel volume overnight until the following morning at which point the data collection was terminated. Flies from each F1 genotype were tested on three independent, overnight flight sessions, and the data from each experiment pooled by genotype. We analyzed the data using a custom Python software pipeline, similar to (Stupski and van Breugel, 2024). Trajectories were filtered for length, minimum distance travelled along the x-axis, and for those which stayed entirely within the wind tunnel volume. We also eliminated any trajectories that re-triggered sham or flash events. From the filtered trajectories, we calculated histograms of ground-speed (x-y) and angular velocities.

### Optogenetic activation of CRZ neurons in males to trigger copulation behaviors

We validated our optogenetic setup by phenocopying a behavior known to be controlled by CRZ neurons (Inagaki *et al*., 2014). Male flies (2-4 days old) of both genetic control and experimental genotypes were positioned on their backs and glued to glass microscope slides using UV-sensitive glue (Bondic) by the wings and pronotum. Each slide had equal numbers of both genotypes mounted. The slides were held almost vertically in a custom-made holder with the flies at a position of approximately 63 x 26 x 37 cm in the wind tunnel (as measured from bottom, rear, downwind corner of the flight space).

Similar to the free-flight experiments described above, light levels were measured by holding a light meter at both horizontal and vertical positions at the approximate location of the glued files, to capture light from the LEDs mounted above the wind tunnel (which include the red LEDs for optogenetic stimulation), as well as LEDs from both the top and bottom of the flight space illuminating the flies from the downwind tunnel portion. Under ambient, non-optogenetic stimulation (illumination with blue and white LEDs only), light intensity from above measured 666-668 lux from the top, and 303-305 lux from the side. Under bright optogenetic and ambient stimulation (blue, white, and red LED illumination), top-down luminance was 10,800-10,820 lux and 1823-1825 lux from the side. As the flies were 4-5 cm higher than the highest light measurement position for the free-flight experiments, we can place a minimum bound of 10.7 and 52 μW/mm^2^ from top-down illumination for ambient and ambient plus optogenetic stimulation irradiance, respectively. From the side, we can place minimum mean irradiance at 4.8 and 11 μW/mm^2^ for ambient alone, and ambient with optogenetic light stimulation, respectively.

Slides were recorded under constant ambient light from blue, white, and infrared LEDs described previously, and optogenetic stimulation was elicited by red LEDs using the following protocol: 30 s baseline with ambient LEDs only, followed by 15 s of red LED stimulation in addition to ambient, then ended by a second 30 s period of ambient-only LED exposure. For each slide, the protocol was repeated 5 times. During each stimulation trial, the flies were recorded using a USB digital camera (acA2040-120um, Basler AG, Ahrensburg, Germany) connected to a laptop, running the manufacturer’s video software (pylon Viewer). Flies were viewed at 200% magnification, using a manual exposure setting allowing good contrast and detail during high-intensity red LED illumination.

We developed an abdominal ethogram of observed behaviors, including those involved in copulation. The following behaviors were chosen for analysis: (1) abdominal curling (without eversion of male genitalia), (2) abdominal curling with genital eversion, (3) genital eversion without curling, (4) targeted grooming of genitalia of ≥ 1 s in duration, (5) ejaculation with visible sperm emission. The responses to optogenetic stimulation were quantified in both control and experimental F1 flies (*n =* 20 for each genotype).

### Statistical analyses

<u>Starvation survival</u>: survival over time was visualized by converting cumulative death counts to percent survival. Plots show mean percent survival ± one standard error of the mean (SEM) at each time point using the ggplot2 package (v 4.0.3) in R-Studio (v 4.5.3). To conduct statistical analysis of survival between genotypes, cumulative death counts were used to assign each fly a time of death corresponding to the hour at which it was first observed dead. Flies still alive at the final check were censored at that timepoint. Kaplan-Meier survival curves were estimated using the survfit function in the survival package (v 3.8-6). Overall differences in survival across genotypes were assessed using the log-rank test, and pairwise comparisons between genotypes were performed using the log-rank test with Bonferroni correction for multiple comparisons (pairwise_survdiff, survminer package v 0.5.2).

<u>Feeding assay</u>: total feeding volume per fly (μL/fly) was compared across genotypes using one-way analysis of variance (ANOVA) from the stats package in R-Studio (v 4.5.3). Pairwise post hoc comparisons were performed using Tukey’s honestly significant difference (HSD) test as implemented in the rstatix package (v 0.7.3). Significance brackets were added to plots using the ggpubr package (v 0.6.3).

<u>Egg-laying assay</u>: to assess the effects of genotype and time on egg-laying, we fit a linear mixed-effects model using the lmer function in the lme4 package (v 2.0-1) in R-Studio (v 4.5.3). Egg count was modeled as a function of genotype, day, and their interaction (fixed effects), with a random intercept for replicate to account for repeated measurements taken from the same flies across days. Day was treated as a categorical factor rather than a continuous predictor, so that the model estimated genotype-specific means at each timepoint. Degrees of freedom and p-values for fixed effects were estimated using the Satterthwaite approximation as implemented in the lmerTest package (v 3.2-1). The genotype × day interaction tested whether the pattern of differences between genotypes varied across days. Post hoc pairwise comparisons between genotypes within each day were performed on the estimated marginal means using the emmeans package (v 2.0.3), with p-values adjusted using Tukey’s method.

### Prediction of paracrine and endocrine networks

Single-cell transcriptomes of AKH-producing cells and all tissues (stringent version) were obtained from the Fly Cell Atlas (Li *et al*., 2022). We manually reclassified the cell clusters for different tissues since some tissues included artefacts or cells that are not unique to a particular tissue (e.g., hemocytes). Cell types present in multiple tissues were classified as “general”. Unannotated and artefact clusters were excluded. Head and body clusters were also excluded since they included cell types that were present in individual tissues.

NSC transcriptomes were identified from the brain single-cell transcriptomes generated previously (Davie *et al*., 2018). The parameters used to identify the different NSC types were based on previous studies and provided below (Kahsai *et al*., 2010; Kapan *et al*., 2012; Miyamoto and Amrein, 2014; Cannell *et al*., 2016; Yang *et al*., 2018; Nässel and Zandawala, 2019; Oh *et al*., 2019; Hadjieconomou *et al*., 2020; Mizuno *et al*., 2021; Zandawala *et al*., 2021; Gera *et al*., 2025).

l-NSC^DH31^ (6 cells): ITP > 2 & Dh31 > 4 & amon > 0 & Phm > 0

l-NSC^CRZ^ (4 cells): Crz > 3 & sNPF > 3 & Dh44 == 0 & ITP == 0 & ChAT == 0 & Gr64a == 0 & Phm > 0

l-NSC^ITP^ (7 cells): Tk > 1 & sNPF > 1 & ITP > 1 & ImpL2 > 1 & Crz == 0

m-NSC^DH44^ (6 cells): Dh44 > 2 & CG13248 > 0 & CG13743 > 0 & Lkr > 0 & Phm > 0

m-NSC^DILP^ (14 cells): Ilp2 > 3 & Ilp3 > 3 & Ilp5 > 3 & ChAT == 0

m-NSC^DMS^ (5 cells): Ms > 2 & EcR > 0 & rk > 0 & amon > 0 & Phm > 0

SEZ-NSC^CAPA^ (3 cells): Capa > 0 & CrzR > 0 & trp > 0 & amon > 0 & Phm > 0

SEZ-NSC^Hugin^ (15 cells): Hug > 3 & Dh44-R2 > 0 & amon > 0 & Phm > 0

To determine the strength of paracrine connections between NSC classes, we multiplied the composite expression scores (see text below for calculation) of each neuropeptide with those of its corresponding receptor. Some neuropeptides mediate their effects via two receptors. If both receptors were expressed in a given cell-type, we only considered the one with higher expression for the sake of simplicity. To minimize false positives, neuropeptides were subjected to a two-step filtering process. First, only those expressed in at least 50% of the cells within a given cluster were retained. Second, a composite expression score was calculated for each neuropeptide by multiplying its average expression by its percent detection. These values were normalized to the maximum observed signal across the dataset, and only hormone-cluster pairs maintaining a relative score of 0.25 or higher were included in the final analysis. This stringent filtering approach was implemented to focus the analysis on dominant neuropeptides and to exclude contamination from ambient RNA. To account for lower abundance of receptor transcripts, we used a more permissive threshold for receptors by retaining those expressed in at least 5% of the cells within a cluster. Unlike the neuropeptides, no secondary relative-score filtering was applied to the receptors to ensure that biologically relevant signaling targets were not prematurely excluded due to low transcript density. We used this thresholding criteria to align previous anatomical studies with our single-cell expression analysis. Since m-NSC^DILP^ produce DILP2, DILP3, and DILP5, all of which target the same receptor, we used their average expression for all analyses.

Analyses were performed in R-Studio (v 2024.04.2+764) using the Seurat package (v 4.4.0 (Hao *et al*., 2021)), unless specified otherwise. Analyses presented in Figure 9E and Figure 9 Supplement 2-4 were performed using Seurat package (v 5.4.0) as the dataset was large.

## Acknowledgements

We are thankful to Emilia Derksen and Zeyu Chang for providing technical assistance, Dick Nässel and Anthony Crown for helpful feedback during preparation of this manuscript and Gilad Barnea, Dan Chuntao and Vivek Jayaraman for fly lines. We thank the Princeton FlyWire team and members of the Murthy and Seung labs, as well as members of the Allen Institute for Brain Science, for development and maintenance of FlyWire (supported by BRAIN Initiative grants MH117815 and NS126935 to Murthy and Seung). Development of the natverse including the coconatfly and fafbseg packages has been supported by the NIH BRAIN Initiative (grant 1RF1MH120679-01), NSF/MRC Neuronex2 (NSF 2014862/MC_EX_MR/T046279/1) and core funding from the Medical Research Council (MC_U105188491). We acknowledge members of the Princeton FlyWire team and the FlyWire consortium for neuron proofreading and annotation. J.G. was supported by funding from the University of Würzburg. M.Z. was supported by funding from the University of Würzburg, German Research Foundation (DFG; ZA1296/1-1), FWO Odysseus Type I grant (WIRED2FEED; G0ASQ25N) and NV INBRE grant from the National Institute of General Medical Sciences (GM103440). Research reported in this publication used the Cellular and Molecular Imaging core facility supported by the National Institute of General Medical Sciences (P30 GM145646). G.M. and N.R. were supported by German Research Foundation grants (FO 207/21 and FO 207/16-1) to C.H.F. We also acknowledge funding from the DFG for the Leica TCS SP8 microscope (251610680, INST 93/809-1 FUGG).

## Author contributions

M.Z. conceived the study. M.Z., F.V.B., P.C. and C.H.F. supervised the project. J.G., G.M., K.C., J.S. and S.H. performed the experimental work. T.H.M., N.R., A.J.G. and M.Z. performed computational analyses. All authors contributed to data visualization. M.Z. wrote the manuscript with input from T.H.M. All authors read, provided feedback and approved the final manuscript.

## Data Availability

Connectivity analysis can be performed using the cell IDs provided at https://codex.flywire.ai/. Custom code and output files are available at: https://doi.org/10.5281/zenodo.20795211

## Code Availability

Custom code used to analyze and visualize the data is available at: https://github.com/Zandawala-lab/McKim-et-al-2026-Synaptic-connectome-of-a-neurosecretory-network-in-the-Drosophila-brain.

---

**Figure 1 Supplement 1:**
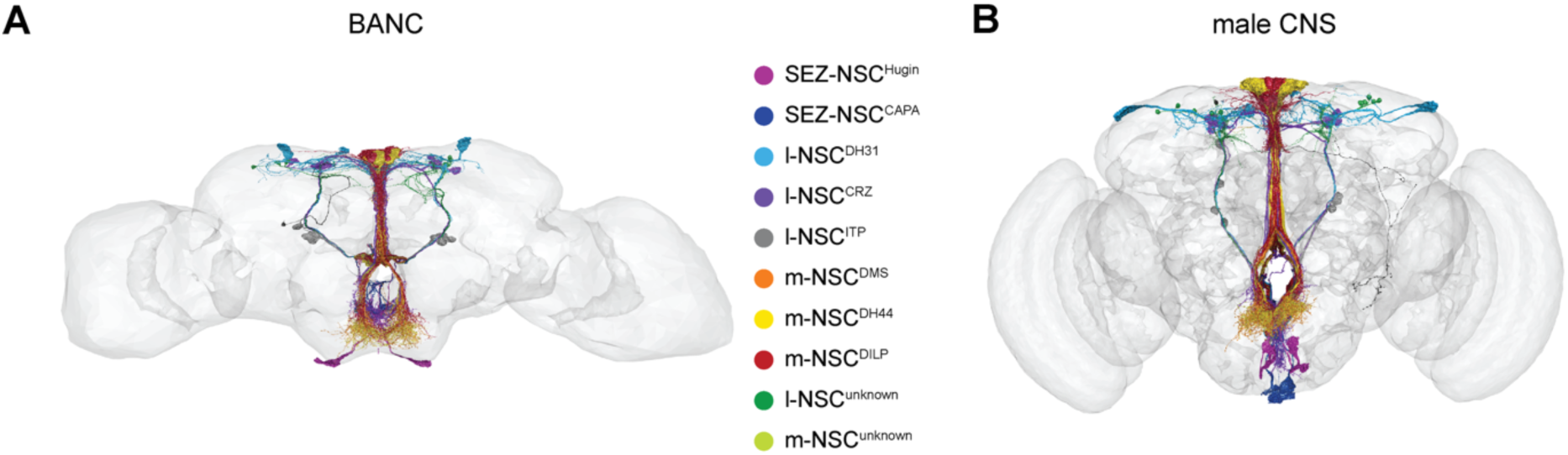
NSC in the brain and nerve cord (BANC) and male central nervous system (maleCNS) connectomes. **(A)** Reconstructions of the 72 NSC within the BANC connectome. **(B)** Reconstructions of the 71 NSC within the maleCNS connectome.

**Figure 2 Supplement 1:**
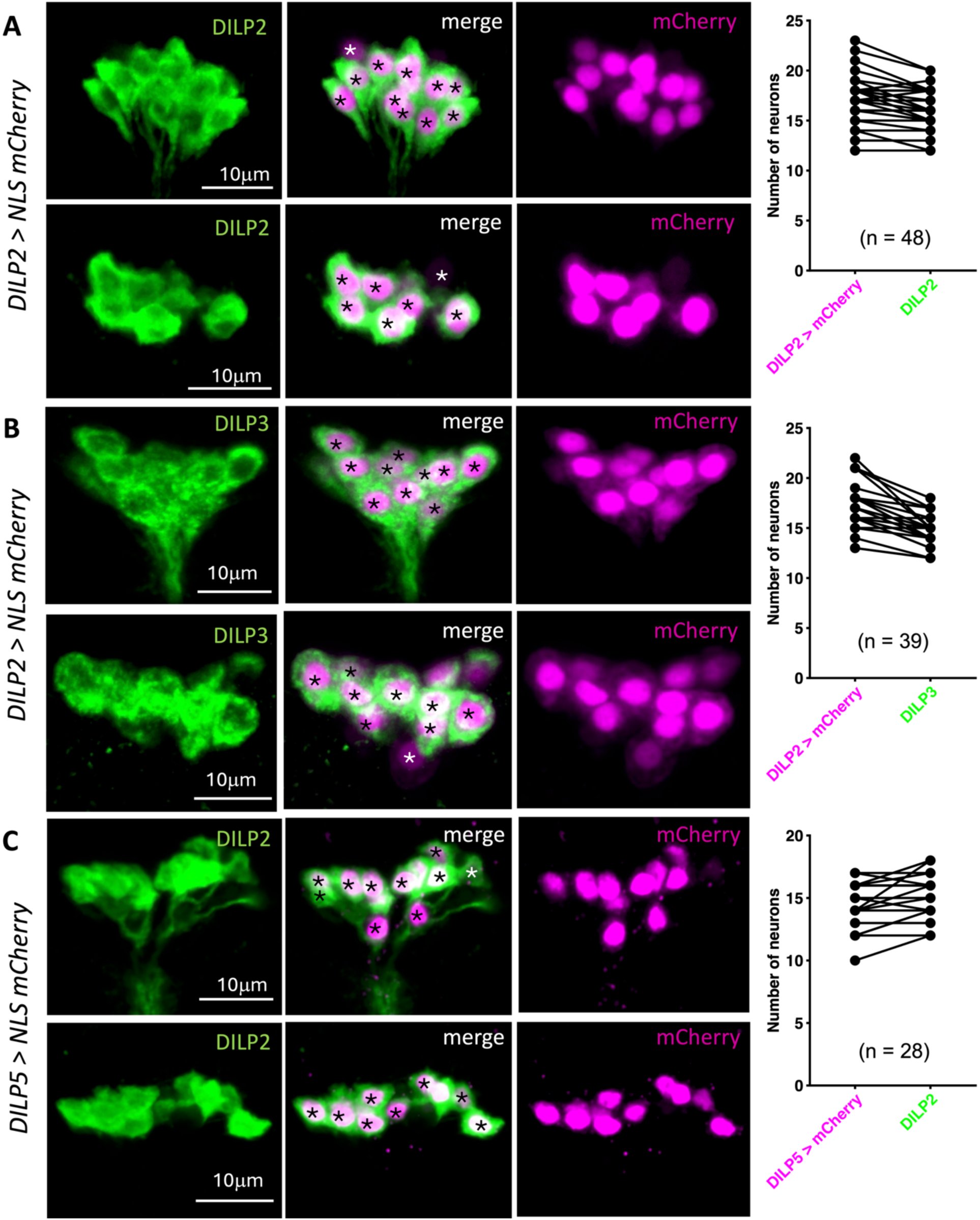
Quantification of m-NSC^DILP^. Quantification of m-NSC^DILP^ (marked with asterisks) as labelled by **(A)** *DILP2-Gal4* driven nuclear mCherry and DILP2 antibody, **(B)** *DILP2-Gal4* driven nuclear mCherry and DILP3 antibody, and **(C)** *DILP5-Gal4* driven nuclear mCherry and DILP2 antibody.

**Figure 2 Supplement 2:**
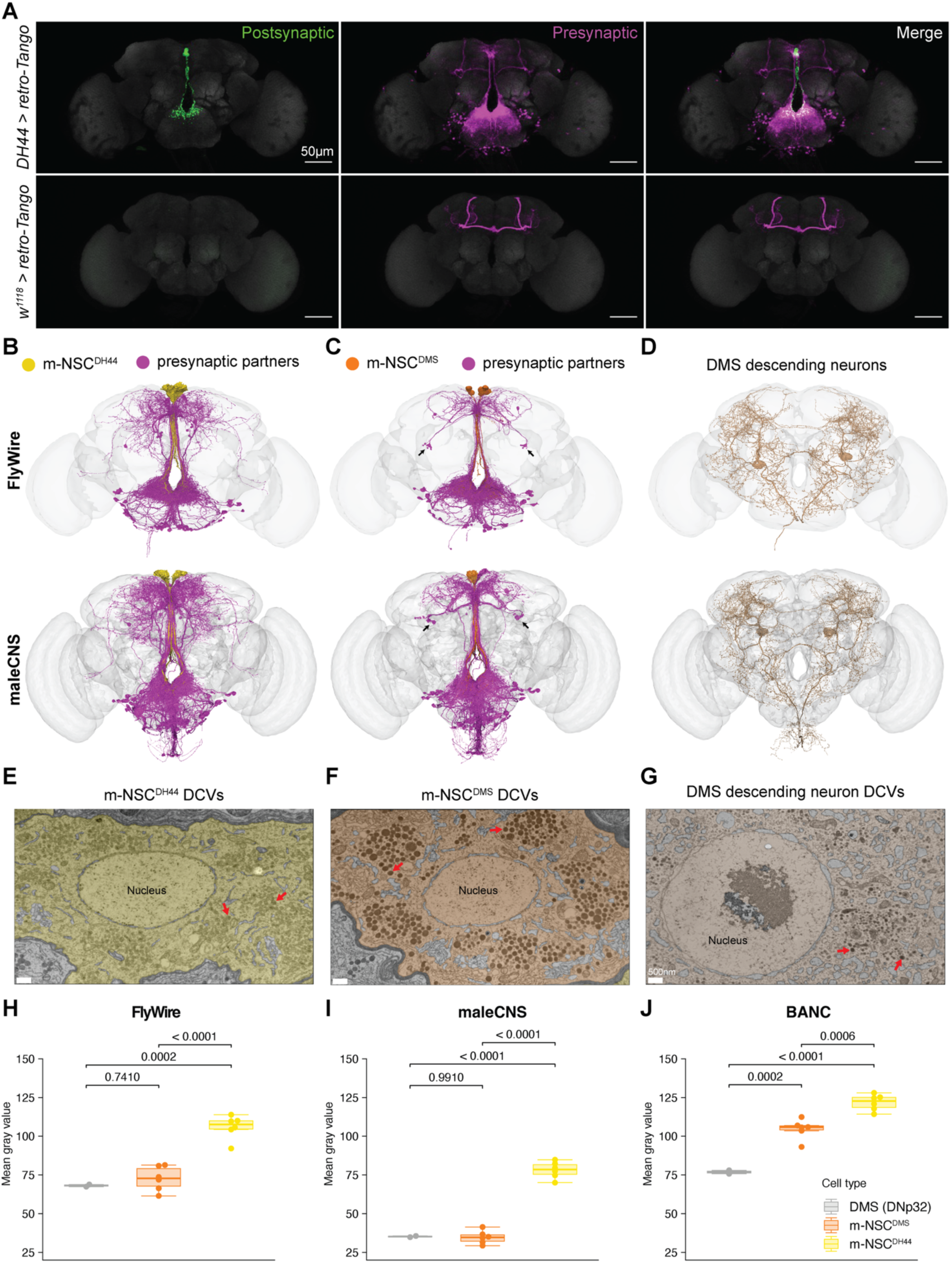
Differences between m-NSC^DH44^ and m-NSC^DMS^. **(A)** Retrograde trans-synaptic labelling of m-NSC^DH44^. m-NSC^DH44^ are labelled in green and their presynaptic partners are labelled in magenta. Note the ectopic expression in the mushroom body which is also visible in the controls. These representative images are based on at least five independent samples. *In silico* retrograde tracing of **(B)** m-NSC^DH44^ and **(C)** m-NSC^DMS^ in the FlyWire and maleCNS connectomes. Both of these NSC classes receive the majority of their inputs from neurons in the SEZ which have similar location and morphology. However, m-NSC^DMS^ also receive inputs from a group of central neurons (marked with an arrow) that are not visible in (A) and (B). **(D)** Reconstruction of myosuppressin (DMS) descending neurons (DNp32 cell type) in the FAFB and maleCNS connectomes. Representative electron micrographs showing a cross section of **(E)** m-NSC^DH44^, **(F)** m-NSC^DMS^ and **(G)** DMS descending neuron soma. Both types of DMS-expressing cells have darker dense core vesicles (marked by red arrows) compared to those found in m-NSC^DH44^. Quantification of DCV brightness for the three cell types (m-NSC^DH44^, m-NSC^DMS^ and DNp32) across the **(H)** FlyWire, **(I)** maleCNS and **(J)** BANC connectomes. For panels (H-J), statistical significance was assessed by one-way ANOVA followed by Tukey’s HSD for multiple comparisons.

**Figure 2 Supplement 3:**
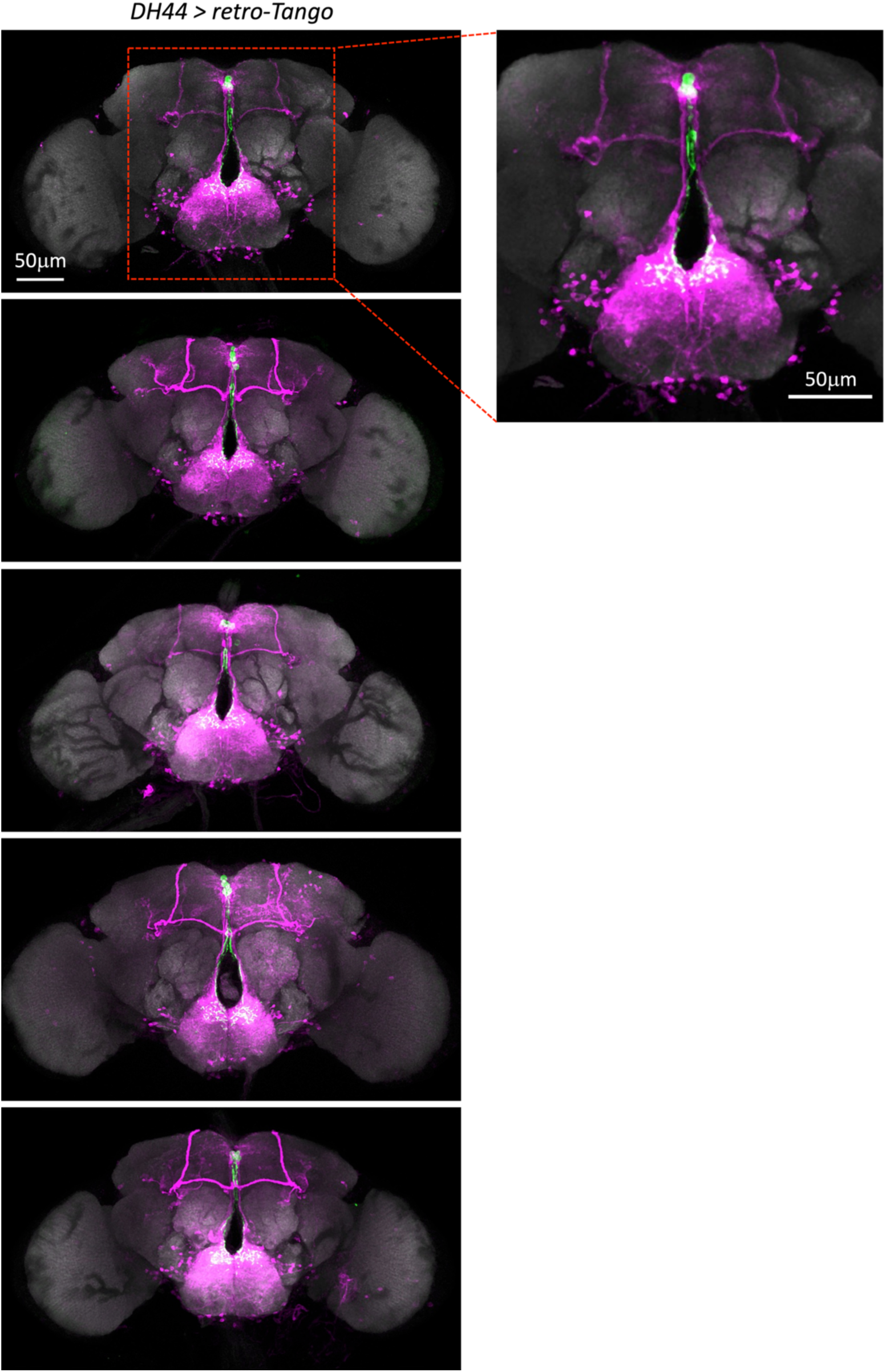
Retrograde trans-synaptic labelling of m-NSC^DH44^. m-NSC^DH44^ are labelled in green and their presynaptic partners are labelled in magenta. Note the ectopic expression in the mushroom body. Presynaptic neurons in the subesophageal zone are consistently labelled across five independent samples.

**Figure 2 Supplement 4:**
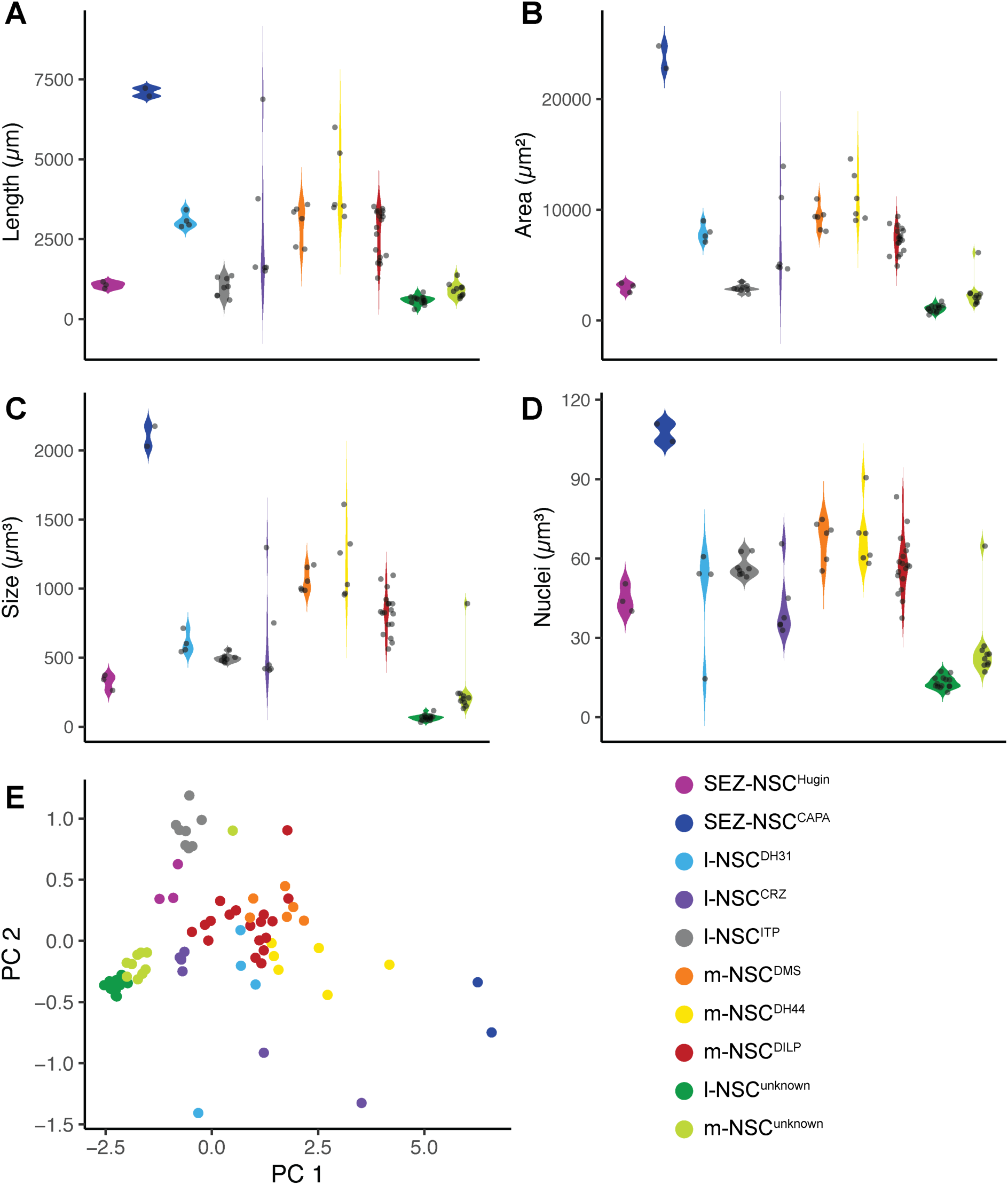
Morphological characteristics of NSC. **(A)** cable length, **(B)** surface area, **(C)** cell volume and **(D)** nuclei volume of different NSC classes. **(E)** Principal component analysis of these four features reveals that the NSC of a given class generally cluster together. Note the high variability for l-NSC^CRZ^, l-NSC^DH31^, m-NSC^DH44^ and m-NSC^DILP^ populations, suggesting that they comprise morphologically heterogenous subpopulations.

**Figure 3 Supplement 1:**
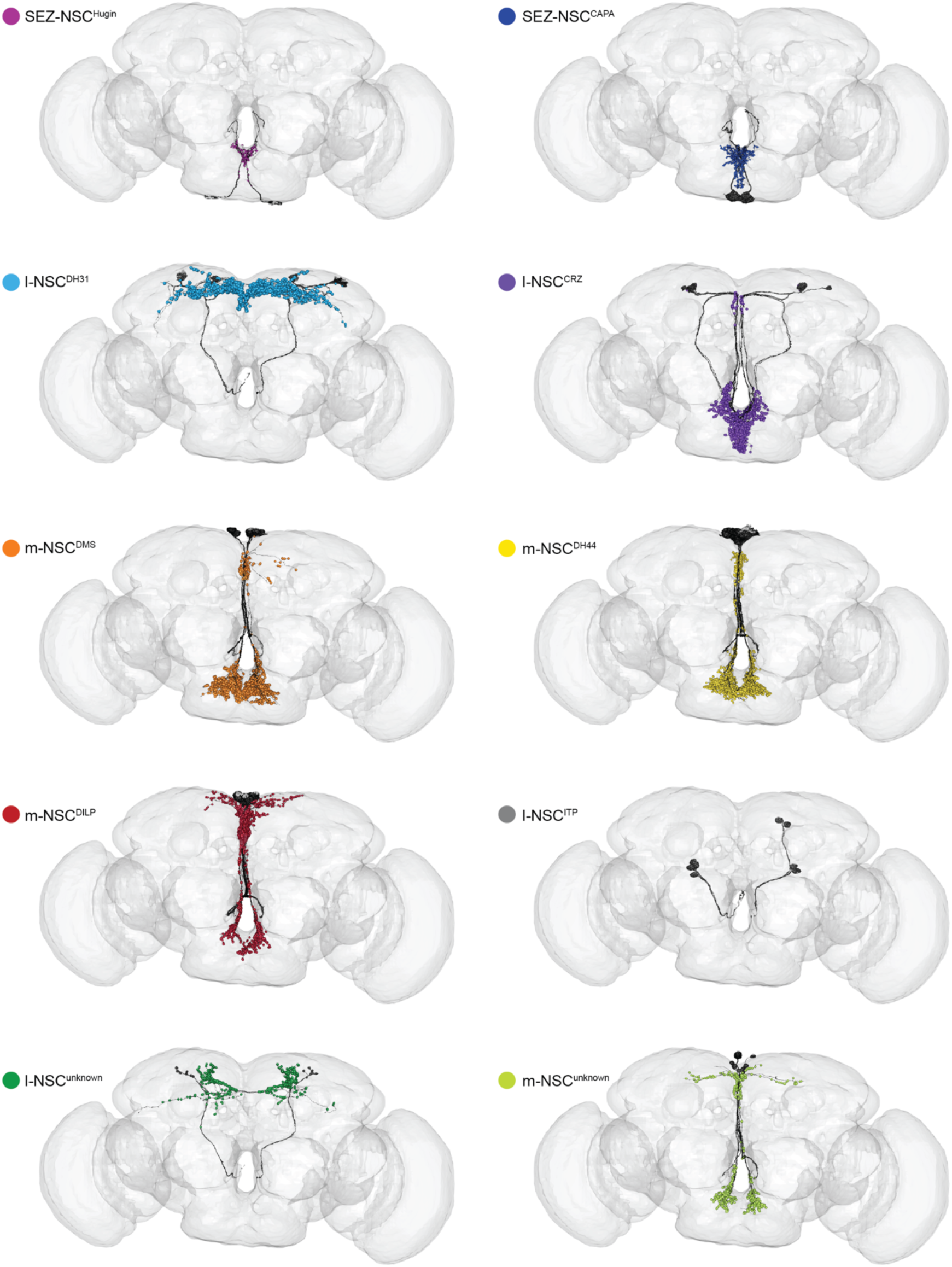
Postsynaptic sites of NSC in the FlyWire connectome. Reconstructions of different NSC classes along with their postsynaptic sites. l-NSC^ITP^ are an exception and have very few postsynaptic sites.

**Figure 3 Supplement 2:**
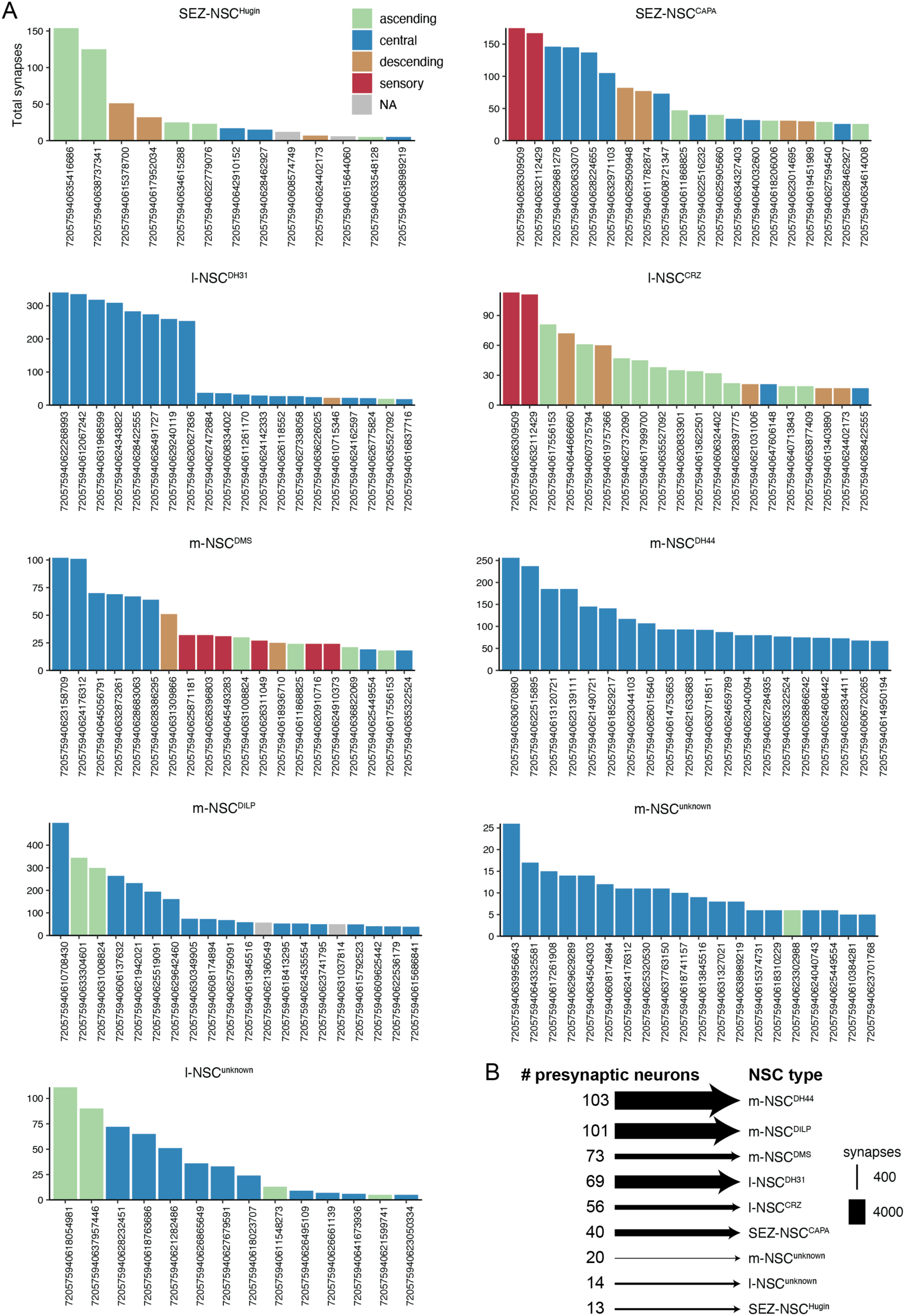
Inputs to NSC classes in the FlyWire connectome. **(A)** Individual presynaptic partners of different NSC sorted based on the number of synapses. Presynaptic neurons are colored based on the super class they belong to. Only the top 20 neurons are shown. SEZ-NSC^CAPA^ and l-NSC^CRZ^ receive strong sensory inputs whereas l-NSC^DH31^, m-NSC^DH44^ and m-NSC^unknown^ mostly receive inputs from central neurons. **(B)** Number of presynaptic neurons providing inputs to different NSC classes.

**Figure 3 Supplement 3:**
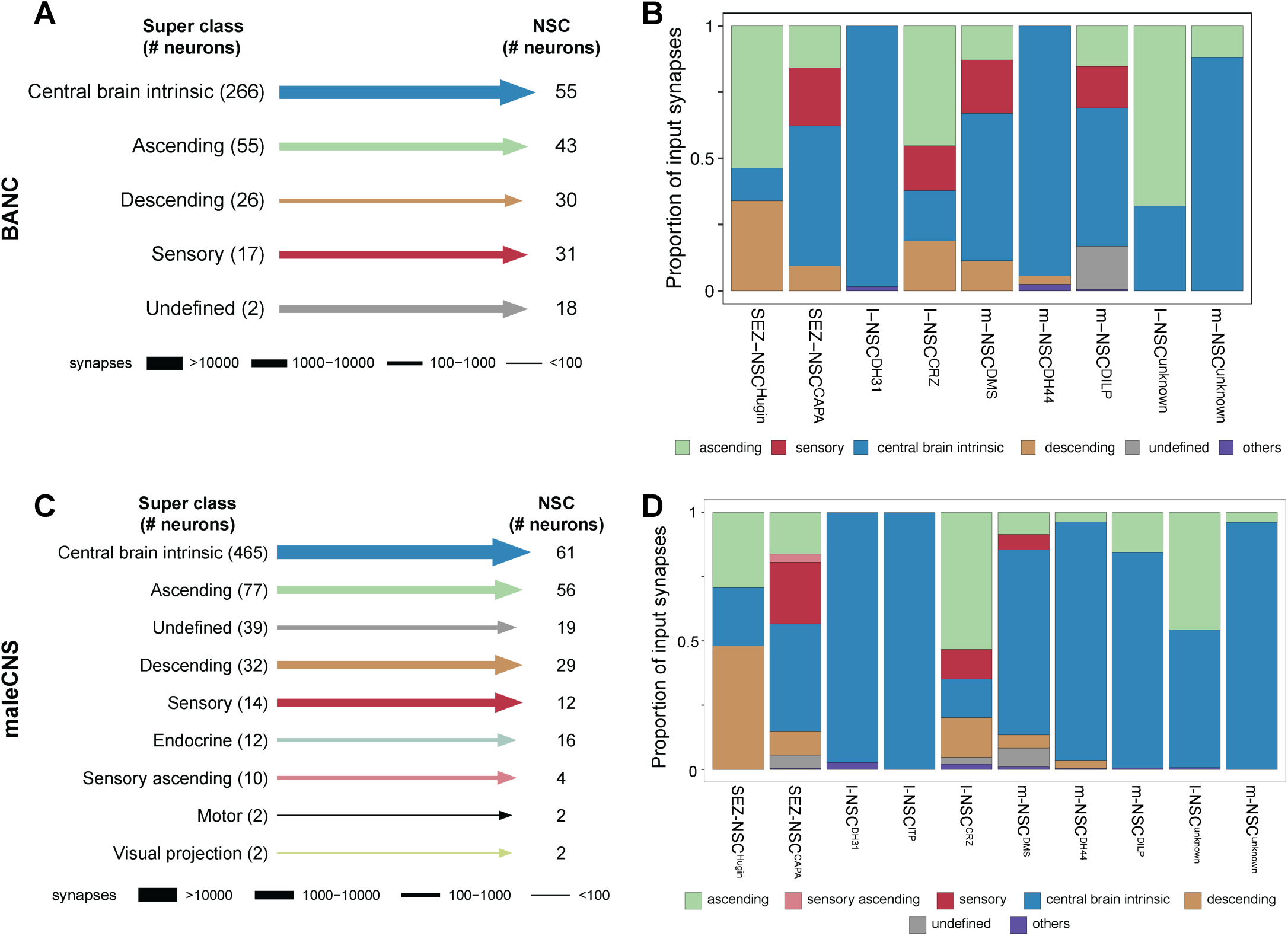
Synaptic inputs to NSC in the BANC and maleCNS connectomes. **(A)** Input to NSC grouped by the neuronal super classes annotated in the BANC connectome. **(B)** Proportion of inputs from various neuronal super classes to different NSC classes in the BANC connectome. **(C)** Input to NSC grouped by the neuronal super classes annotated in the maleCNS connectome. **(D)** Proportion of inputs from various neuronal super classes to different NSC classes in the maleCNS connectome. Note that central brain intrinsic neurons followed by ascending neurons are the largest groups providing inputs to NSC across both connectomes.

**Figure 3 Supplement 4:**
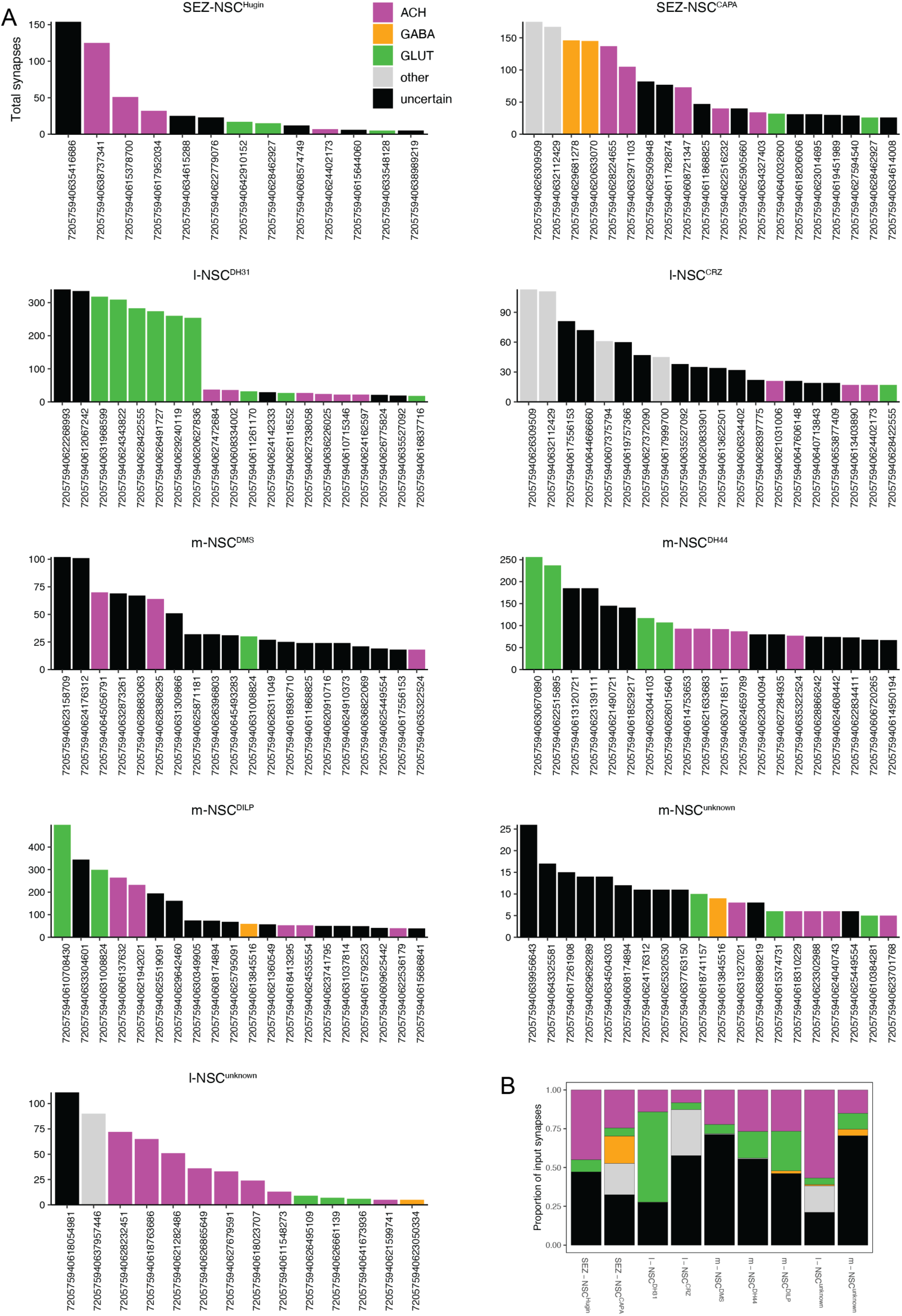
Neurotransmitters providing inputs to NSC classes in the FlyWire connectome. **(A)** Individual presynaptic partners of different NSC sorted based on the number of synapses and colored based on their neurotransmitter identity. l-NSC^DH31^ and m-NSC^DH44^ receive strong glutamatergic inputs. **(B)** Input to NSC grouped by the neurotransmitters. Out of the three fast-acting neurotransmitters, GABA provides the least inputs.

**Figure 3 Supplement 5:**
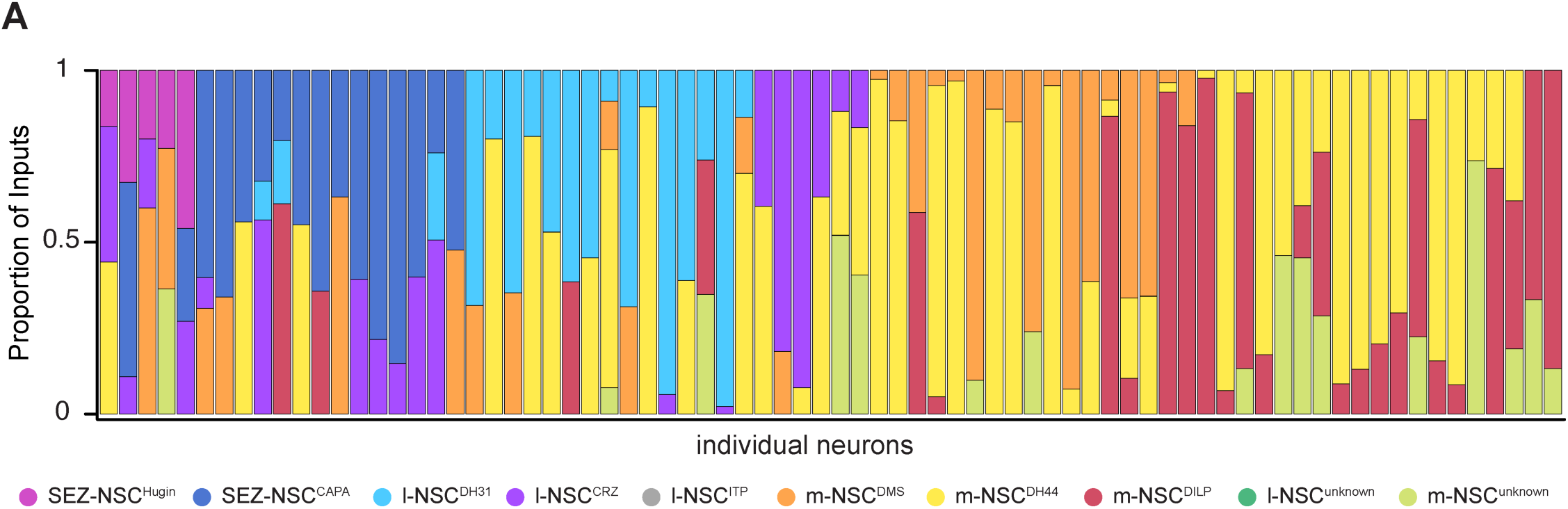
Neurons providing input to multiple NSC classes in the FlyWire connectome. Proportion of inputs (based on the number of synapses) from individual neurons to different NSC classes. Each bar represents an individual neuron and it is filled according to the NSC classes that it provides input to. In total, 76 neurons provide inputs to more than one NSC class, with m-NSC^DH44^ receiving inputs from most of these neurons.

**Figure 4 Supplement 1:**
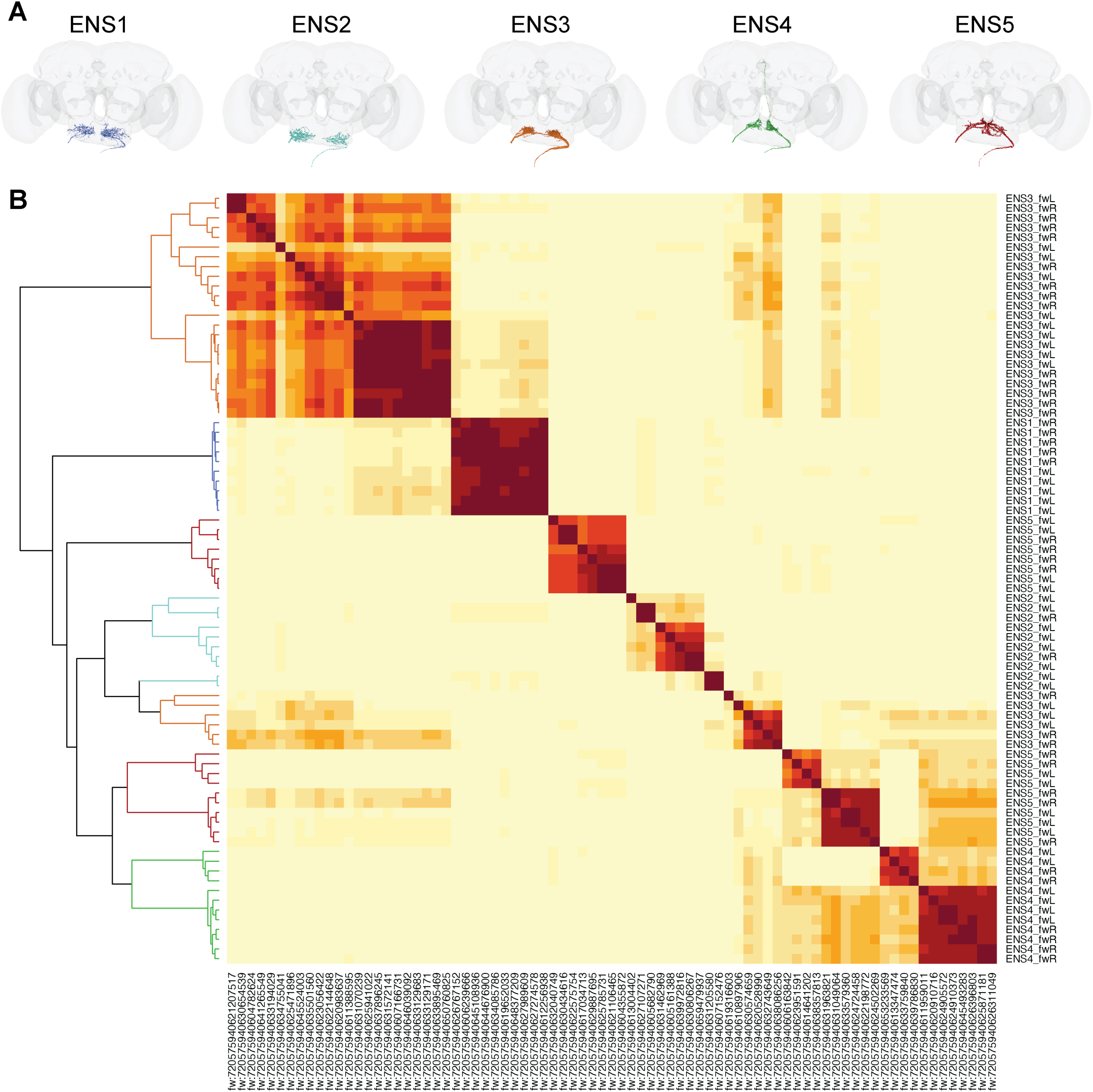
Classification of enteric neurons in the FlyWire connectome. **(A)** Enteric neurons have been classified into five cell types (ENS1 to ENS5) on Codex. Reconstructions of ENS1 to ENS5 show that these are morphologically distinct cell types. **(B)** Cosine similarity matrix of all enteric neurons in the FlyWire connectome based on their total inputs and outputs. Darker red colors indicate higher similarity between neurons. Neurons within the clades are colored based on the cell types in (A). Note that not all cells belonging to ENS3 and ENS5 cell types cluster together, suggesting that they could represent heterogeneous populations based on synaptic connectivity.

**Figure 6 Supplement 1:**
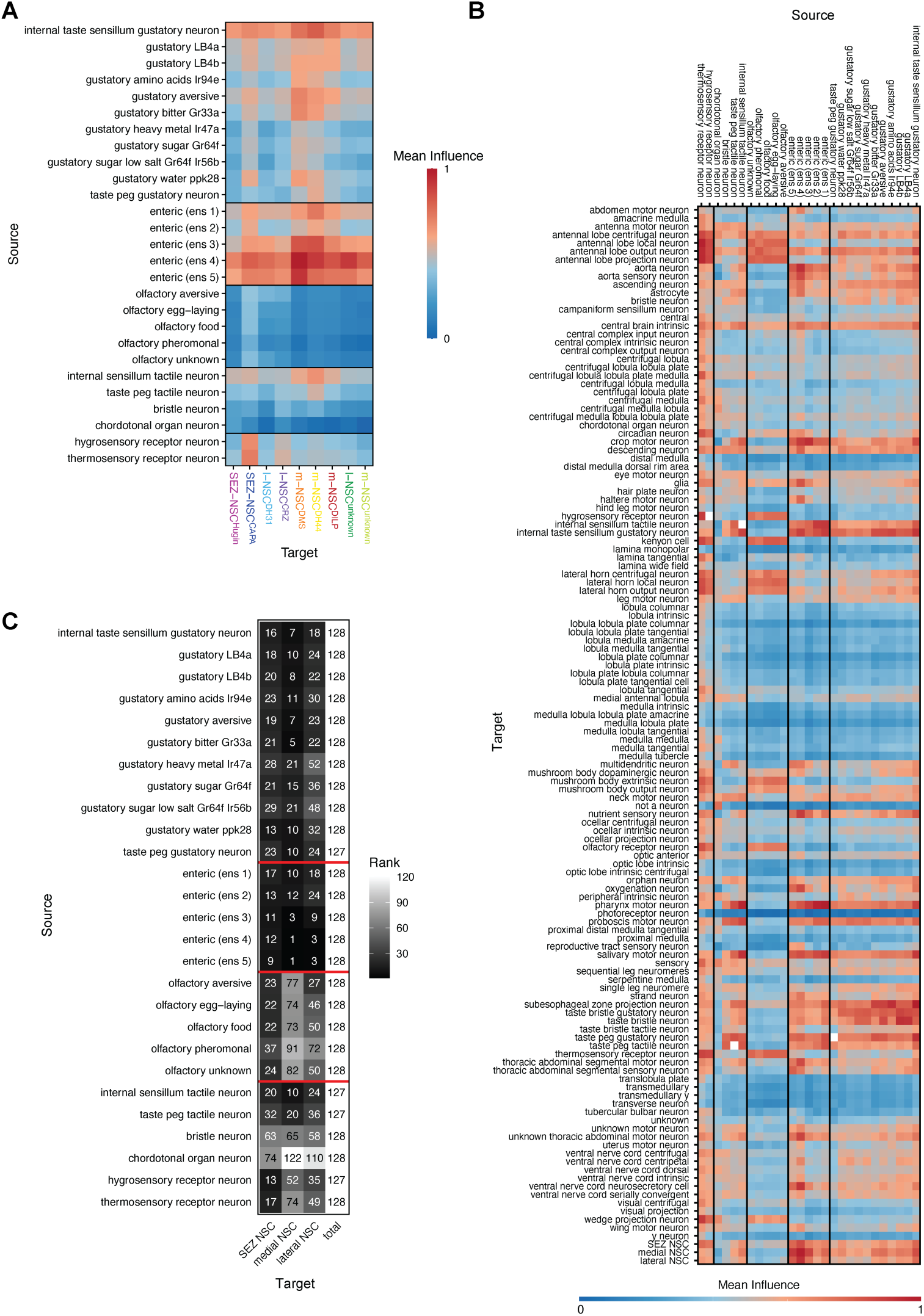
Influence of *in silico* sensory neuron stimulation on NSC in the BANC connectome. **(A)** Mean influence of different sensory neurons (rows) on NSC classes (columns). Note the strong influence of enteric neurons on NSC, especially m-NSC^DMS^. **(B)** Mean influence of different sensory neurons (rows) on major cell types in the brain, including NSC. **(C)** The mean influence in panel (B) was used to determine the rank of influence on NSC in relation to other cell types. Enteric neurons, especially ENS4 and ENS5, have the strongest influence on medial NSC.

**Figure 7 Supplement 1:**
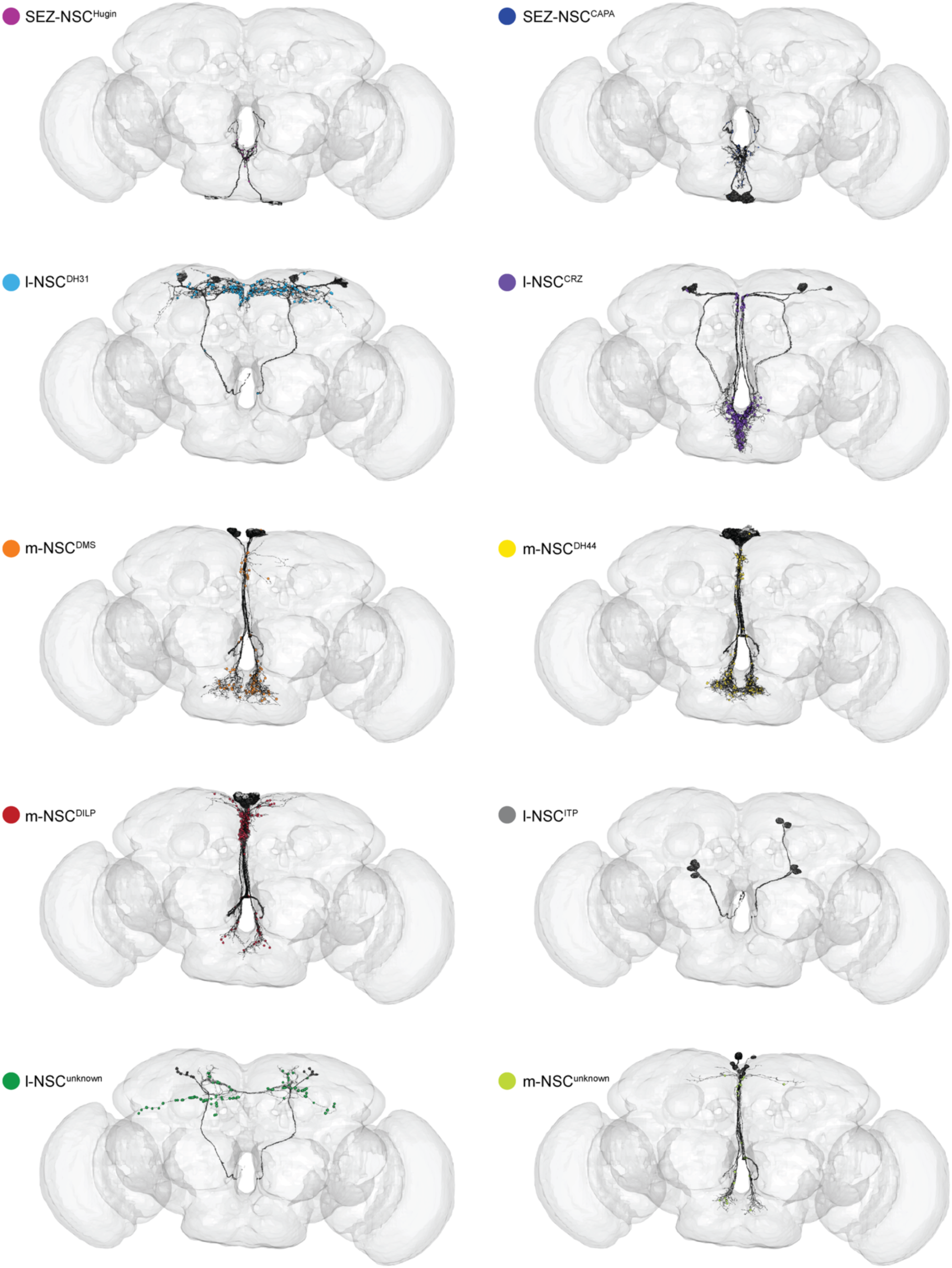
Presynaptic sites of NSC in the FlyWire connectome. Reconstructions of different NSC classes along with their presynaptic sites. l-NSC^CRZ^ have several presynaptic sites in the subesophageal zone.

**Figure 7 Supplement 2:**
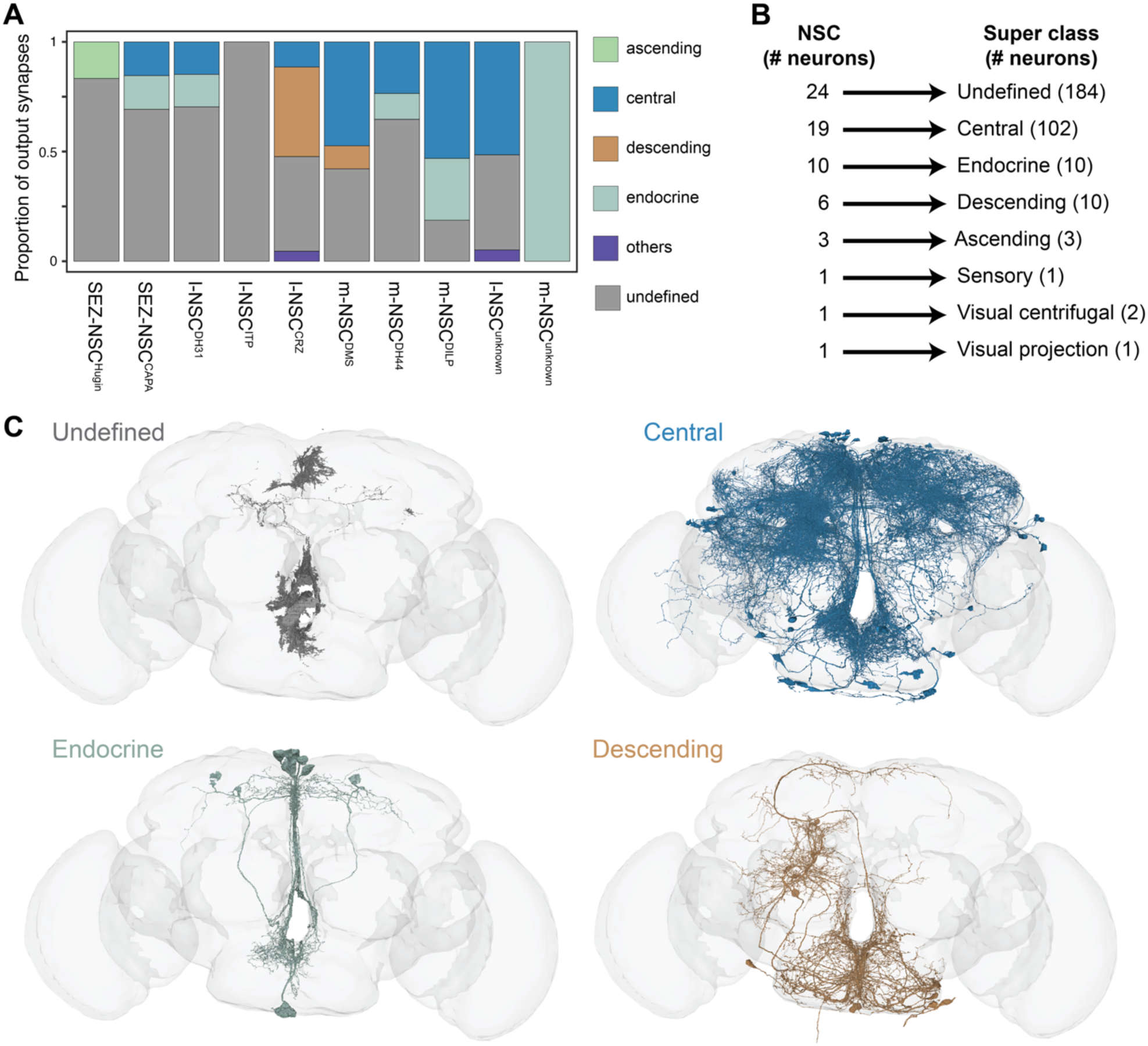
Synaptic output from NSC in the FlyWire connectome based on a low synaptic threshold. **(A)** Proportion of outputs from different NSC classes to various neuronal super classes when the threshold for a significant connection is lowered to 2 synapses. **(B)** Output from NSC grouped by the neuronal super classes annotated in the FlyWire connectome. **(C)** Reconstructions of neurons receiving inputs from NSC. Cells belonging to the top four super classes are shown. Note that most of the output from NSC is to partial fragments and non-neuronal cells (undefined), as well as central neurons.

**Figure 7 Supplement 3:**
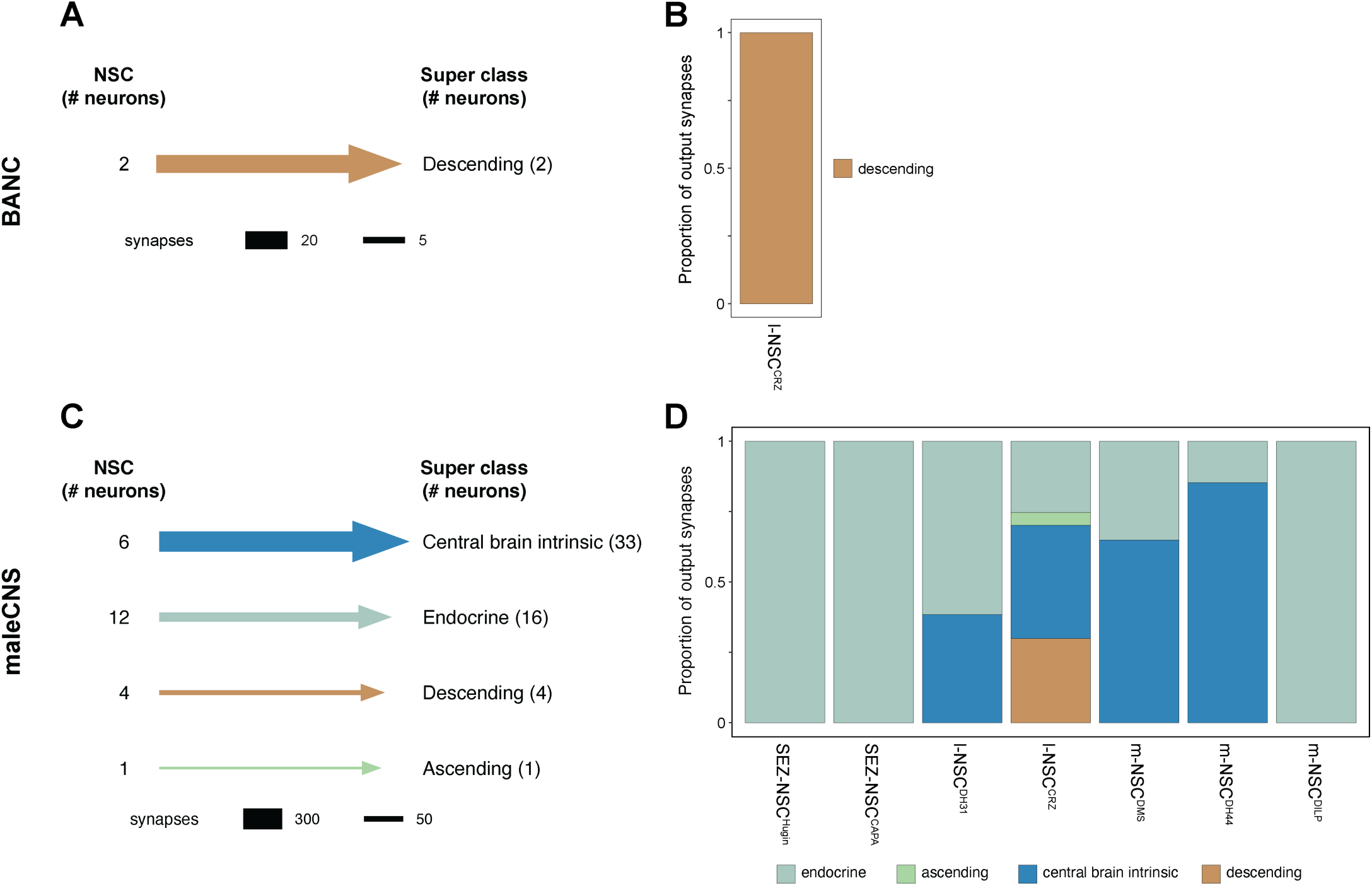
Synaptic output from NSC in the BANC and maleCNS connectomes. **(A)** Output from NSC grouped by the neuronal super classes annotated in the BANC connectome. **(B)** l-NSC^CRZ^ provides all of its output to descending neurons in the BANC connectome. **(C)** Output from NSC grouped by the neuronal super classes annotated in the maleCNS connectome. **(D)** Proportion of outputs from different NSC classes to various neuronal super classes in the maleCNS connectome.

**Figure 8 Supplement 1:**
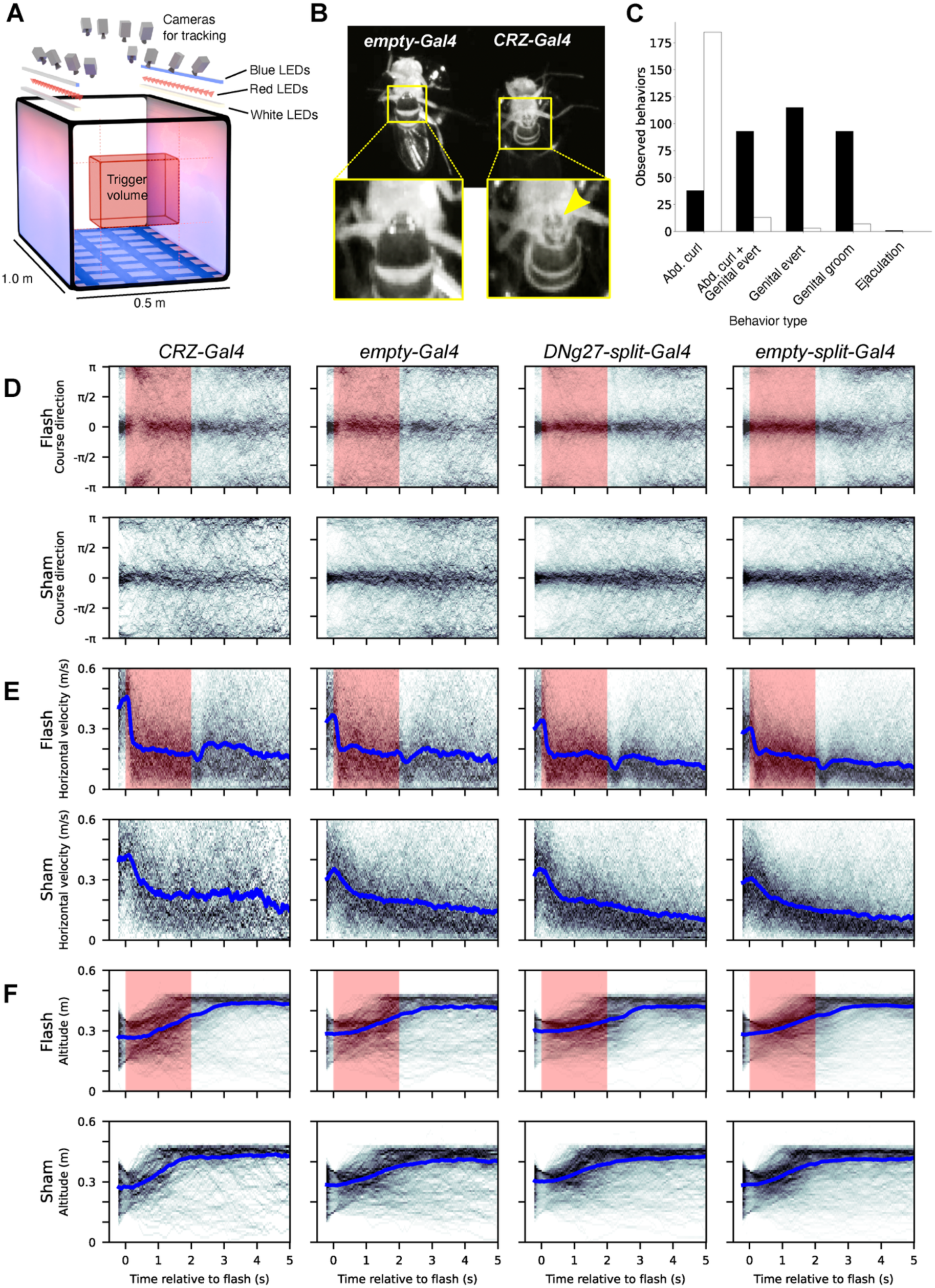
Optogenetic activation of CRZ or DNg27 neurons in female flies results in minimal large-scale changes in free-flight kinematics. **(A)** Schematic diagram of the wind-tunnel used for free-flight kinematic experiments, annotated with dimensions and the positions of cameras and LEDs. A schematic representation of the trigger volume, which activates the red LEDs upon entrance of a fly, is also indicated. **(B)** Still images of male *empty-Gal4* and *CRZ-Gal4* flies taken from videos during optogenetic stimulation. The images compare two related behaviors observed during optogenetic stimulation: (1) an abdominal curl demonstrated by the *empty-Gal4* control fly (left), and (2) a copulatory abdominal curl with genital eversion by the experimental *CRZ-Gal4* male (right). The extrusion of internal genital structures (yellow arrow) is evident in the zoomed-in portions. **(C)** Comparison of the number of observed abdominal behaviors in control (*empty-Gal4* in white bars; N = 20) and experimental (*CRZ-Gal4* in black bars; N = 20) male flies during optogenetic stimulation. Behaviors were counted over n = 5 x 15 s red LED flashes, with an inter-stimulus interval of 120 s of ambient-only illumination. **(D)** Course direction (radians) versus time relative to the fly entering the trigger volume for flash (top row) and sham (bottom row) trigger events. The shaded region denotes the period of red LED illumination during flash events. N = 60-78 flies for each genotype: *CRZ-Gal4*, flash n = 412 trajectories, sham n = 198 trajectories; *empty-Gal4*, flash n = 344 trajectories, sham n = 379 trajectories; *DNg27-Gal4*, flash n = 519 trajectories, sham n = 491 trajectories; *empty-split-Gal4*, flash n = 769 trajectories, sham n = 711 trajectories. A course direction of 0 indicates upwind and *π*/-*π* is downwind. **(E)** Horizontal ground speed velocity (m/s) versus time relative to the fly entering the trigger volume for flash (top row) and sham (bottom row) trigger events. The blue line represents the median groundspeed velocity over all trajectories. Fly and trajectory numbers are the same as in panel (D). **(F)** Altitude (m) versus time relative to the fly entering the trigger volume for flash (top row) and sham (bottom row) trigger events. The blue line represents the median altitude over all trajectories. Fly and trajectory numbers are the same as in panel (D).

**Figure 8 Supplement 2:**
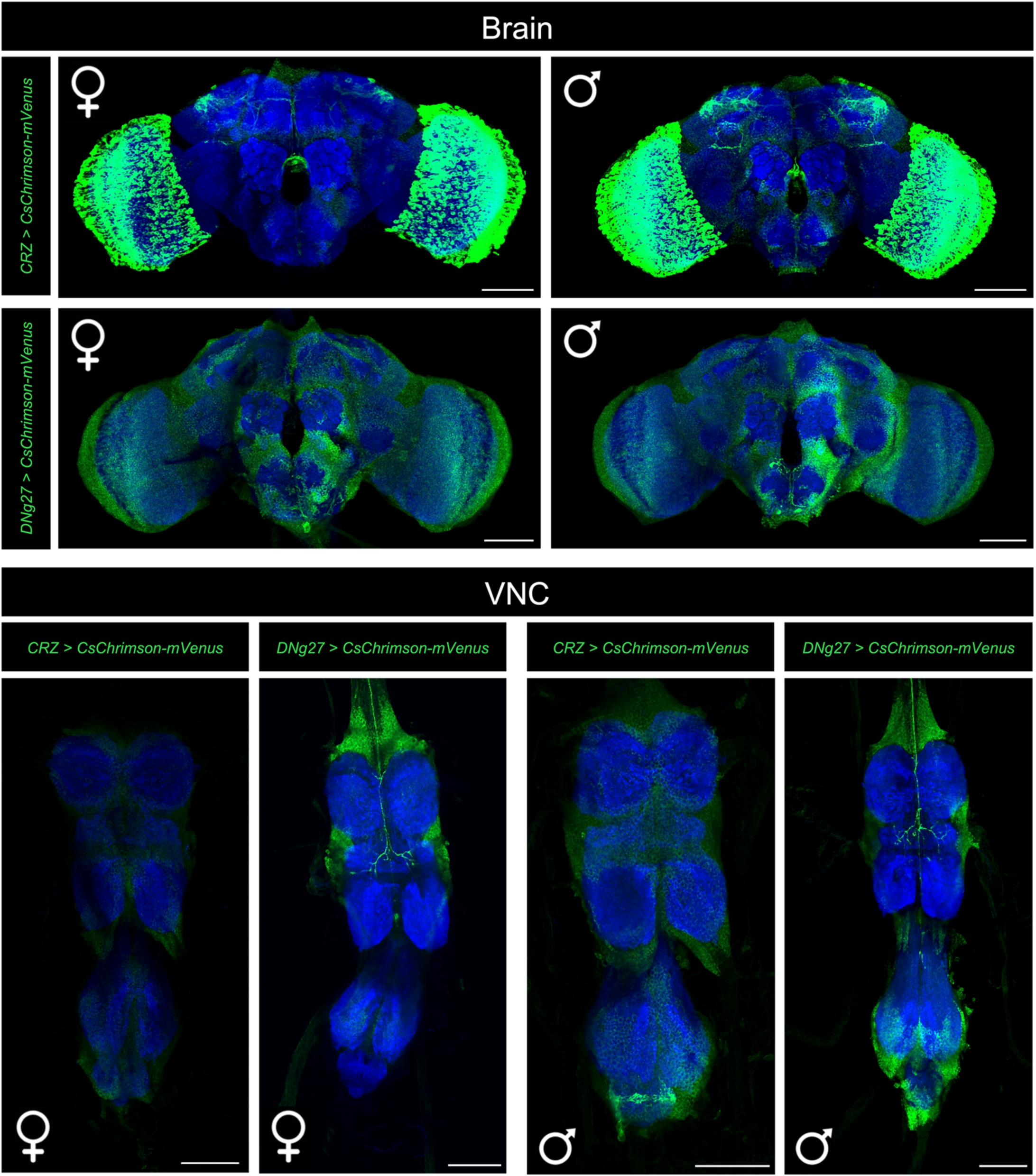
Neurons labelled by CRZ-Gal4 and DNg27-Gal4 in males and females. *CRZ-Gal4* drives CsChrimson-mVenus expression in l-NSC^CRZ^ and optic lobe neurons in both males and females. Additionally, it drives expression in neurons in the abdominal ganglion of the ventral nerve cord in males. *DNg27-Gal4* (a split-Gal4 driver) drives weak CsChrimson-mVenus expression in a pair of descending neurons in both males and females.

**Figure 9 Supplement 1:**
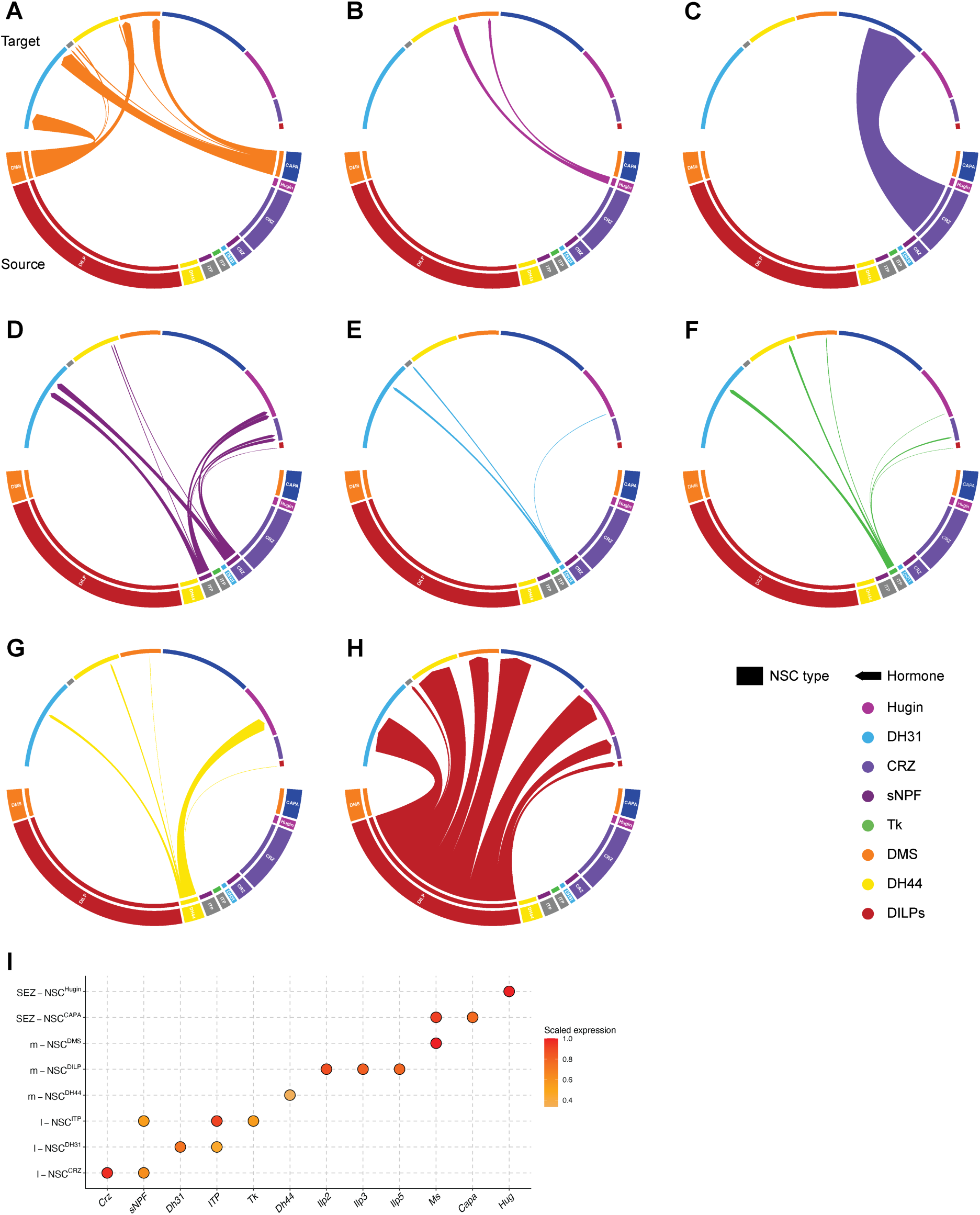
Putative paracrine interconnectivity between NSC. NSCs classes targeted by **(A)** myosuppressin (DMS), **(B)** Hugin, **(C)** corazonin (CRZ), **(D)** short neuropeptide F (sNPF), **(E)** diuretic hormone 31 (DH31), **(F)** tachykinin (TK), **(G)** diuretic hormone 44 (DH44) and **(H)** insulin-like peptides (DILPs). Ion transport peptide and CAPA pathways are not included because their receptors were not detected in these transcriptomes. **(I)** Dot plot showing the neuropeptides expressed in each NSC class following thresholding. The expression has been scaled and was used to generate the connectivity diagrams in Figure 9C and Figure 9 Supplement 1A-H.

**Figure 9 Supplement 2:**
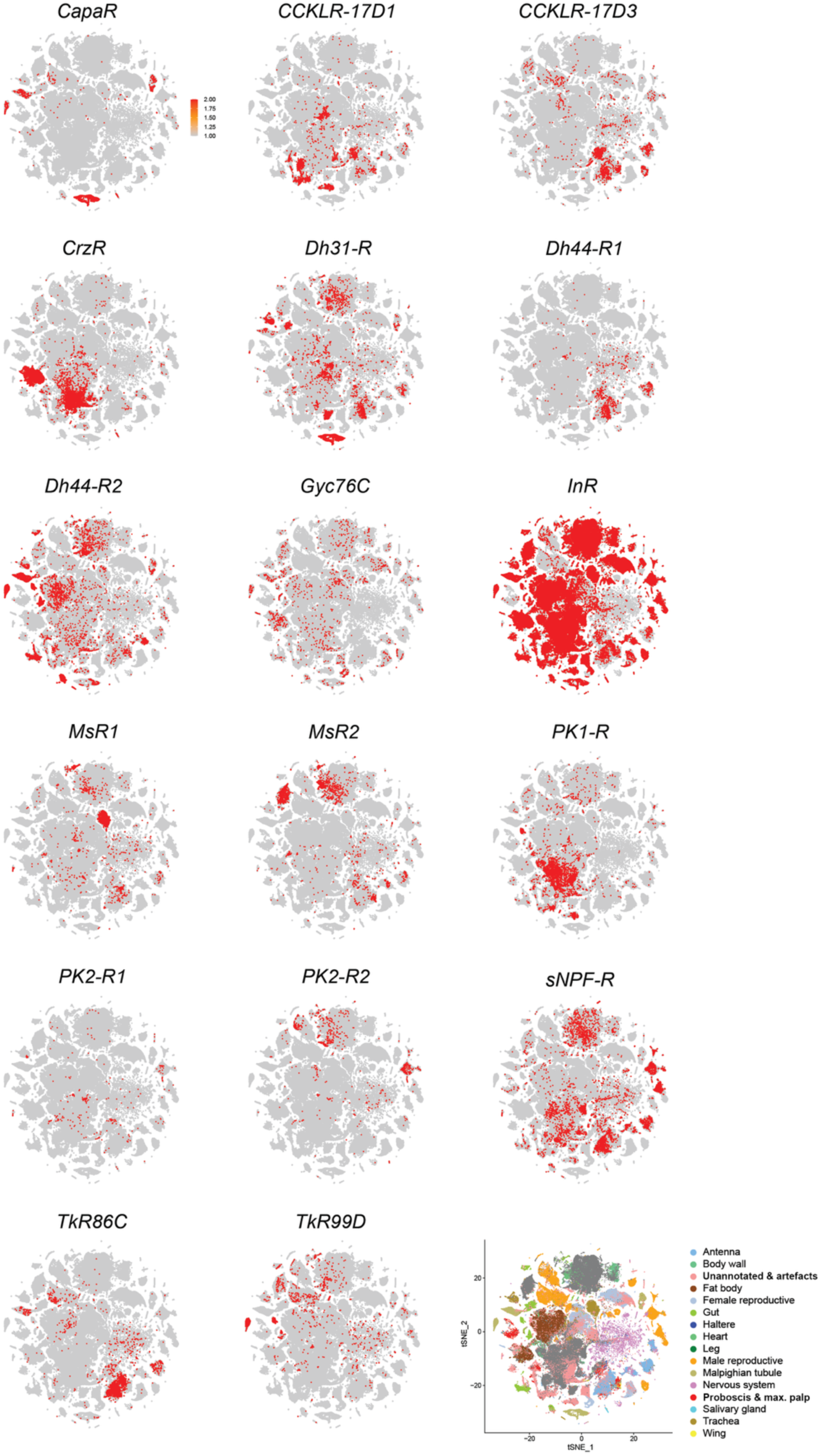
Expression of receptors for hormones released from brain NSC. t-SNE plots showing expression of hormone receptors across single-cell transcriptomes from all *Drosophila* tissues (Li *et al*., 2022). Note that some receptors such as *InR* and *sNPF-R* are broadly expressed whereas others such as *CapaR* and *PK2-R1* are sparsely expressed.

**Figure 9 Supplement 3:**
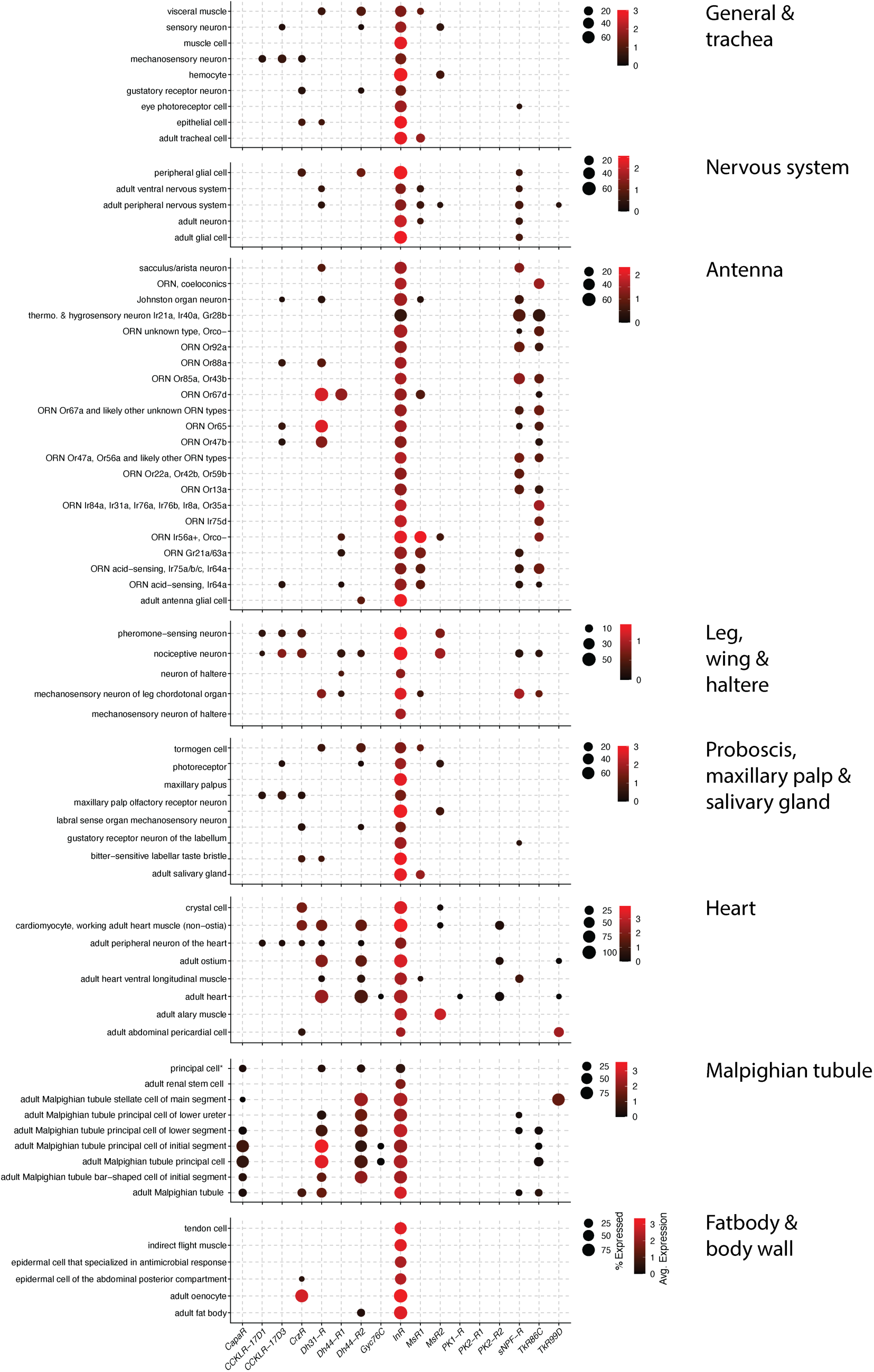
Expression of hormone receptors in peripheral tissues. Dot plots showing expression of hormone receptors in different tissues at single-cell resolution. Expression of only those receptors whose corresponding neuropeptides are expressed in brain NSC are shown.

**Figure 9 Supplement 4:**
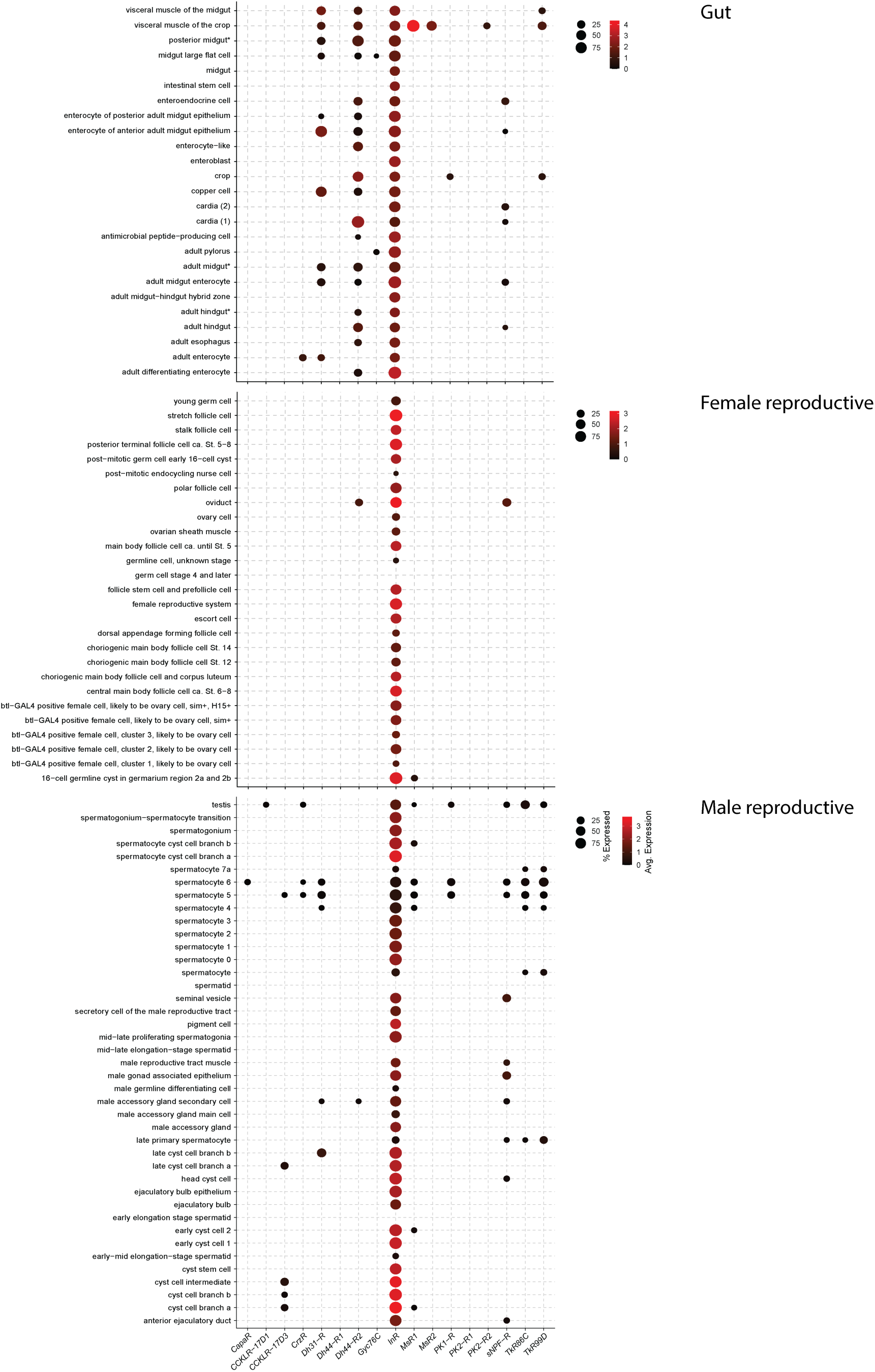
Expression of hormone receptors in the gut and reproductive tissues. Dot plots showing expression of hormone receptors in the gut and reproductive tissues at single-cell resolution. Expression of only those receptors whose corresponding neuropeptides are expressed in brain NSC are shown.

## Supplementary Tables

**Supplementary Table 1:** Fly strains used in this study.

| Fly strain | BDSC Stock number / Reference |
| --- | --- |
| *DH44-Gal4* | 51987 |
| *DILP2-Gal4* | (Ikeya *et al.*, 2002) |
| *DILP3-Gal4* |  |
| *DILP5-Gal4* | 66008 |
| *MS-T2A-Gal4* | 84652 |
| *Gr64a-Gal4* | 57661 |
| *CRZ-Gal4* | 51976 |
| *R24F06-p65.AD/CyO; R45E06-DBD/TM6B (DNg27-split-Gal4)* | 75837 |
| *empty-split-Gal4* | 79603 |
| *empty-Gal4* | 68384 |
| *w<sup>1118</sup>* | 5905 |
| *UAS-retro-Tango* | (Sorkac *et al.*, 2023) |
| *JFRC81-10xUAS-IVS-Syn21-GFP-p10 (EGFP)* | (Pfeiffer *et al.*, 2012) |
| *10xUAS-myr::GFP (myrGFP)* | (Pfeiffer *et al.*, 2010) |
| *UAS-NLS-mCherry* | 38425 |
| *20xUAS-CsChrimson::mVenus* | Janelia Farms |
| *UAS-Kir2.1::EGFP* | 6595 |

**Supplementary Table 2:**
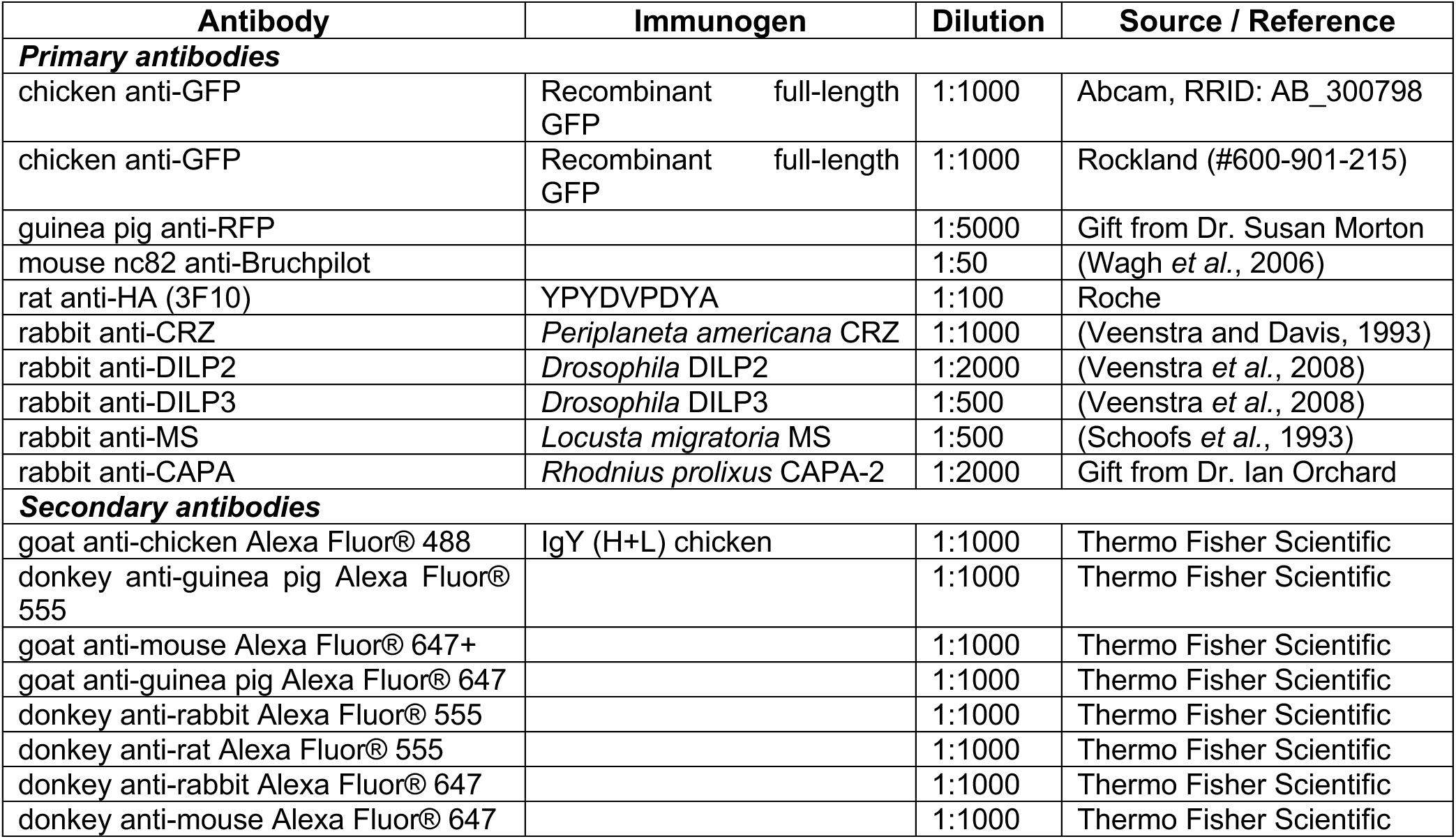
Antibodies used for immunohistochemistry in this study.

**Supplementary Table 3:** Cell IDs of NSC identified in FlyWire, BANC and maleCNS connectomes (see separate file)

